# Hypoimmunogenic human motor neurons induced from iPSCs in vivo substantially ameliorate ALS disease in large animal models

**DOI:** 10.1101/2025.09.03.673895

**Authors:** Na Zhang, Yang Yang, Mengqi Chen, Qingjian Zou, Longquan Quan, Jiayuan Huang, Quanjun Zhang, Yu Zhao, Zhen Ouyang, Zhongtian Zhang, Yuning Song, Min Chen, Kun Zhang, Amar Deep Sharma, Michael Ott, Xiaojiang Li, Zhanjun Li, Wenguang Xie, Zhen Dai, Liangxue Lai

## Abstract

Stem cell-based therapy holds great potential for substituting degenerated motor neurons (MNs) in amyotrophic lateral sclerosis (ALS). Missing protocols for advanced differentiation of transplanted cells into MNs, immune rejection, and the lack of suitable ALS models for preclinical trials have slowed the development of effective therapies. Here, we employed multiplex genetic-editing to generate a novel human pluripotent stem cell line containing doxycycline (Dox)-inducible MNs-specific transcription factors and comprehensively modified immunomodulatory genes. We transplanted these cells into the spinal cord of ALS large animal models (SOD1^G93A^ pigs and TIA1^P362L^ rabbits), which faithfully recapitulate pathologies and symptoms observed in ALS patients. The transplanted cells could efficiently differentiate into functional MNs upon Dox treatment in vivo, distribute throughout the spinal cord and motor cortex via extensive migration, survive long-term without the need for immunosuppression. Notably, these MNs integrated into host neural circuits, as evidenced by their long projection of peripheral axons to target muscle and reformation of neuromuscular junctions. As result, pathologies and motor deficits were substantially ameliorated in both animal models.

**One Sentence Summary:** Hypoimmunogenic human motor neurons induced from iPSCs *in vivo* reform neuromuscular junctions and ameliorate ALS disease in pig and rabbit models.

## INTRODUCTION

Amyotrophic lateral sclerosis (ALS) is a multifactorial neurodegenerative disease with 90% of cases diagnosed as sporadic and only 10% as inherited (*1, 2*). The pathophysiology is characterized by progressive loss of upper and lower motor neurons (MNs) in the spinal cord, brainstem and cerebral cortex, leading to paralysis and death within 3-5 years of onset. Despite advances in our understanding of the molecular and cellular mechanisms, a cure for ALS remains elusive to date (*3, 4*). Cell therapy represents a promising approach for treating various forms of ALS due to its potential to replace the dying MNs. Over the past two decades multiple studies have investigated cell therapies for ALS using different types of cells such as hematopoietic stem cells, mesenchymal stem cells, neural progenitor cells, and glial progenitor cells and different administration routes (intravenous, intracerebroventricular, intrathecal, and intranasal)(*5–14*) in ALS rodent models and in patients. However, all of these studies have yielded limited benefits in treating ALS due to failure of replacing degenerated MNs in the long term.

There are three major challenges in cell replacement therapy for ALS: (1) the incomplete differentiation of MNs *in vivo* has provided a crucial hurdle for cell integration. Most transplants remained predominantly undifferentiated (*15*) or developed into astrocytes rather than MNs (*1, 15, 16*); (2) vigorous immune rejection against the transplanted cells prevented the long-term survival of grafts (*17–19*); (3) rodent models have been criticized to not reflect the core phenotype seen in ALS patients, as they exhibit a limited loss of MNs (*20–24*). To address the issue of MN differentiation, we have recently reported engineered human iPSCs with Dox-inducible expression of three MN-specific transcription factors (*NGN2, ISL1, LHX3*) and forced anti-apoptotic gene expression (*Bcl-XL*) (NILB-iPSCs). The cells were previously manufactured at large scale *in vitro* and were successfully transplanted and induced into MNs *in vivo* in immunodeficient mice (*25*). Genetic modification of the immunomodulatory genes in stem cells could enhance immune tolerance of their derivatives in allogeneic or xenogeneic settings (*19, 26–35*). Additionally, we and other groups have showed that some pathological features resembling human ALS disease only appear in large animal models but not in small rodents (*23, 24, 36, 37*) such as intranuclear inclusions, cytoplasmic distribution of TAR DNA-binding protein 43 (TDP-43) and massive MNs degeneration, which emphasize an urgent demand for utilizing large animal models to study ALS therapies (*38*).

Based on the above concepts, we hypothesized that transferring hypoimmunogenic human iPSCs with Dox-inducible MN-specific transcription factors into ALS large animal models could support MN replacement in ALS. We found that our cells were able to differentiate into MNs upon induction of Dox, migrate along the spinal cord and motor cortex, and survive long term without using immunosuppression in ALS large animal models (transgenic SOD1^G93A^ pigs and point mutation TIA1^P362L^ rabbits) and without evidence of tumor formation. Most importantly, for the first time, we observed successful integration of these MNs into host neural circuits, manifested by their long-distance peripheral axonal projection and reformation of neuromuscular junctions (NMJs) between axon terminals of human stem cell-derived MNs and host muscle fibers, leading to the rescue of various ALS pathological and biochemical features including TDP-43 mislocalization, muscle atrophy, neurotoxic microenvironments and improved electromyographic activity in both pig and rabbit models. Thus, our results provide convincing evidence that human stem cell-derived MNs successfully replace degenerated MNs of ALS hosts and remarkably ameliorate disease progression.

## RESULTS

### Generation of hypoimmunogenic human iPSCs through multiplex genetic modifications of immunomodulatory factors

To address the challenges from either innate or adaptive immune rejections triggered by allogenic or xenogeneic neural stem cells (NSCs) transplantation (*39, 40*), as well as hostile neuroinflammation environments in ALS hosts (*41*), we employed a combined strategy to engineer NILB-iPSCs to confer them superior immune tolerance, resulting in a hypoimmunogenic cell line named as HIP-NILB-iPSCs (**Fig. 1A and fig. S1A)**. These modifications included: (i) disruption of HLA-I and II expression by knocking out β2-microglobulin (*B2M^-/-^*) and class II major histocompatibility complex trans-activator (*CIITA^-/-^*) to prevent T cells-mediated immune rejection(*28, 42*) **(fig. S1B)**; (ii) overexpression of immune checkpoint inhibitors to overcome ‘missing-self’ killing response caused by HLA-I ablation, such as *CD47* for natural killer (NK) cells and macrophages,(*35, 43*) *CD24* for macrophages,(*44*) programmed cell death ligand 1 (*PD-L1*) for PD-1^+^ NK cells, PD-1^+^ macrophages and T cells(*31, 45*) **(fig. S1C)**; (iii) overexpression of *CTLA4-Ig* to enhance an immune privilege by targeting allogeneic or even xenogeneic T cells, dendritic cells, and B cells(*19, 33*) **(fig. S1C)**; (iv) knockout of interleukin 6 signal transducer (*IL6ST^-/-^*) and tumor necrosis factor receptor 1 (*TNFRSF1A^-/-^*) to enhance resistance to neuroinflammatory microenvironments in ALS hosts, which are key mediators of inflammatory responses in the central nervous system (CNS) (*46*) **(fig. S1B)**. Both the deletion of *B2M/CIITA/IL6ST/TNFR1* and the overexpression of *CD47/CD24/PD-L1/CTLA4-Ig* genes in the final HIP-NILB-iPSCs were verified by western blot analysis **(fig. S1D)**.

**Fig 1.**
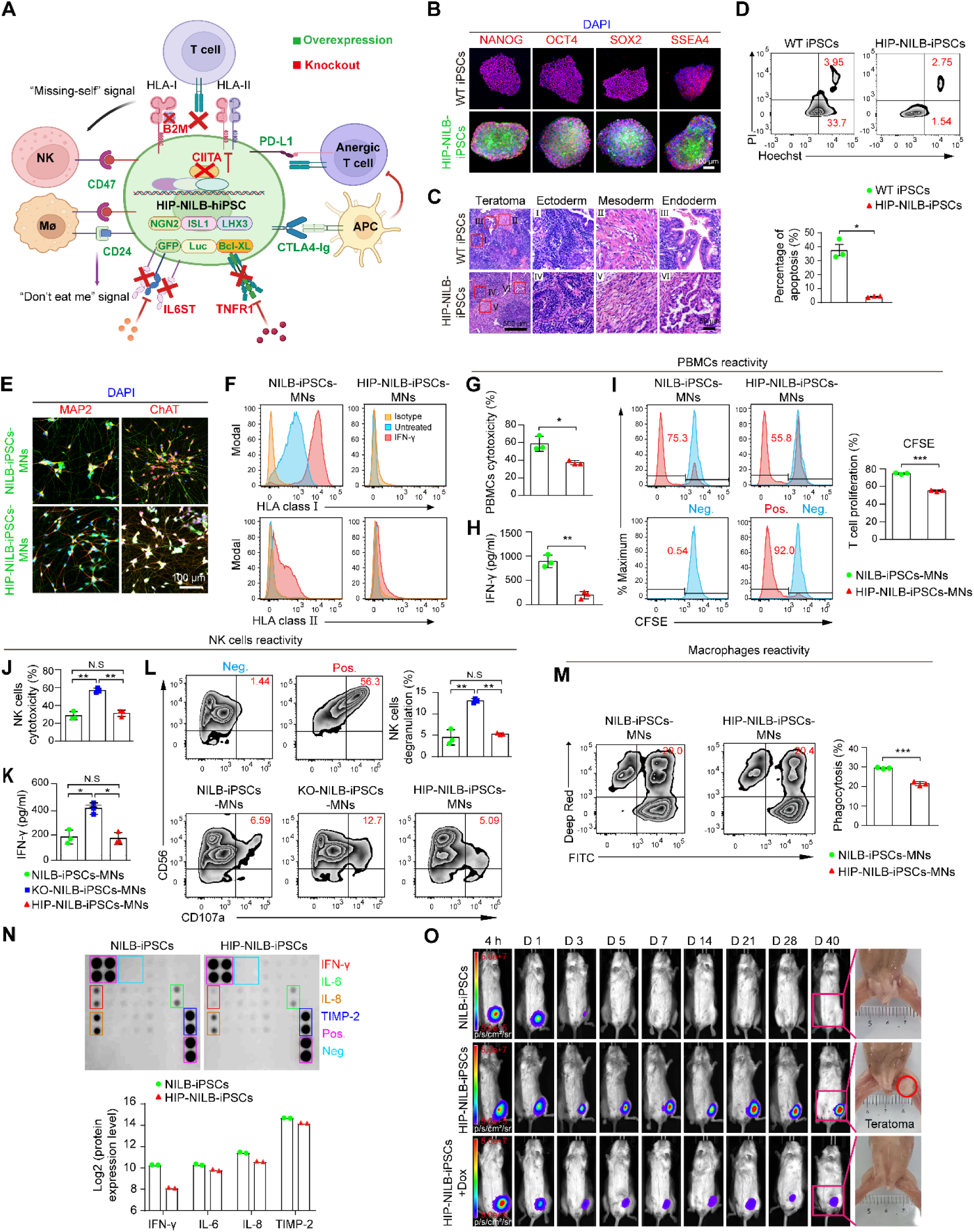
Generation and characterization of hypoimmunogenic HIP-NILB-iPSCs. **A)** Schematic diagram of engineered HIP-NILB-iPSCs created by BioRender. (**B)** Immunofluorescence staining of NANOG, OCT4, SOX2 and SSEA4 in WT iPSCs and HIP-NILB-iPSCs. Scale bar, 100 μm. **(C)** H&E staining of three germ layers in WT iPSCs and HIP-NILB-iPSCs-derived teratoma. Framed areas represent different germ layers regions that were enlarged in I-III. Scale bar, 500 μm or 50 μm. **(D)** Apoptosis assay of HIP-NILB-iPSCs and its quantification. WT iPSCs were used as a control. **(E)** Immunofluorescence staining of MAP2 and ChAT in HIP-NILB-iPSCs induced MNs (HIP-NILB-iPSCs-MNs). NILB-iPSCs induced MNs (NILB-iPSCs-MNs) were used as a control. Scale bar, 100 μm. **(F)** HLA class I and II expression in NILB-iPSCs-MNs and HIP-NILB-iPSCs-MNs with or without IFN-γ treatment. Isotype is a negative control with matched primary antibody. **(G)** PBMCs cytotoxicity against HIP-NILB-iPSCs-MNs by measuring LDH release. PBMCs cocultured with NILB-iPSCs-MNs were used as a control for G-I. **(H)** ELISA assay for IFN-γ secretion in PBMCs cocultured with HIP-NILB-iPSCs-MNs. **(I)** CFSE analysis of T cells (CD3^+^) proliferation in PBMCs when cocultured with HIP-NILB-iPSCs-MNs. PBMCs is negative control (Neg.), PBMCs activated by PHA is positive control (Pos.). **(J)** Primary NK cells cytotoxicity against HIP-NILB-iPSCs-MNs by measuring LDH release. NK cells cocultured with NILB-iPSCs-MNs were used as a control for J-L. **(K)** ELISA assay for IFN-γ secretion in NK cells cocultured with HIP-NILB-iPSCs-MNs. **(L)** NK degranulation assay by quantifying CD107a surface expression in CD56^+^ NK cells cocultured with HIP-NILB-iPSCs-MNs. NK cells, Neg., NK cells treated with PHA, Pos.. **(M)** The phagocytic activity of macrophages against NILB-iPSCs-MNs and HIP-NILB-iPSCs-MNs is characterized by Deep Red Cell Tracker-labeled THP-1 derived M1 macrophages. **(N)** Inflammatory-factor array indicates low inflammatory responses of HIP-NILB-iPSCs upon LPS stimulation. **(O)** BLI signals over time for NILB-iPSCs, HIP-NILB-iPSCs without or with Dox-treatment (HIP-NILB-iPSCs +Dox) engrafted in allogeneic humanized-PBMCs NCG mice (n=5). HIP-NILB-iPSCs +Dox group represents 12 hours pre-induction of Dox *in vitro* prior to transplantation, and continued 3 days Dox induction *in vivo* after transplantation. Experiments were independently repeated at least three times. Data are presented as mean ± SEM, p values were determined using a two-tailed, unpaired Student’s *t-test* (g, h, i, j, k, l, m) or Welch’s t-test (D), \**p* < 0.05, \*\**p*< 0.01, \*\*\**p* < 0.001, N.S, not significant.

We proceeded to investigate whether these genomic alterations exerted any effects on HIP-NILB-iPSCs’ pluripotency. To this end, we passaged them more than 20 times and found normal karyotype architectures **(fig. S1E)** and pluripotency, similar to those of wild type (WT) iPSCs, as revealed by expression of NANOG, OCT4, SOX2 and SSEA4 (**Fig. 1B**), as well as global expression profiles **(fig. S1F)**. Teratoma formation assays in immunodeficient mice indicated retained ability to differentiate into three germ layers (**Fig. 1C**). Consistent with our previous findings (*25*), HIP-NILB-iPSCs exhibited lower levels of apoptosis during passage (**Fig. 1D**) and were able to differentiate into MNs *in vitro* upon Dox induction with nearly all cells expressing neuron marker MAP2 and MNs marker ChAT (**Fig. 1E**). Moreover, we performed bulk RNA sequencing to characterize their differentiation process at different culture time points upon Dox treatment, which indicated progressive downregulation of pluripotency genes and upregulation of spinal MNs differentiation and axonogenesis genes **(fig. S1G)**. Taken together, these results demonstrated that the introduced genomic alterations do not compromise the karyotype, pluripotency, anti-apoptosis and MNs differentiation potential of HIP-NILB-iPSCs.

### Validation of superior immune tolerance of Human HIP-NILB-iPSCs in allogeneic recipients

Next, we sought to determine whether HIP-NILB-iPSCs and HIP-NILB-iPSCs-MNs were able to evade both adaptive and innate immune attacks. It is worthy to mention that we combined multiple hypo-immunogenic genes which have been reported and validated in previous studies (*19, 28, 31, 33, 35, 44–46*), thereby we did not further define effects of individual hypo-immunogenic genes. Here, we first analyzed surface expression of HLA-I and HLA-II in HIP-NILB-iPSCs and HIP-NILB-iPSCs-MNs in the presence of interferon-gamma (IFN-γ). The experiment revealed ablation of HLA-I and HLA-II in HIP-NILB-iPSCs and HIP-NILB-iPSCs-MNs, but not in respective controls (**Fig. 1F and fig. S2A)**. Co-culturing HIP-NILB-iPSCs or HIP-NILB-iPSCs-MNs with human peripheral blood mononuclear cells (PBMCs) for 5 days resulted in significantly lower lactate dehydrogenase (LDH) release (**Fig. 1G and fig. S2B)** than respective controls, indicating reduced lysis of HIP-NILB-iPSCs and HIP-NILB-iPSCs-MNs. Secretion of IFN-γ by PBMCs also decreased when co-cultured with HIP-NILB-iPSCs or HIP-NILB-iPSCs-MNs (**Fig. 1H and fig. S2C)**, suggesting a reduced immune response induced by them. Since T cells represent the most abundant cell type and central mediators in PBMCs for immune rejection, we examined whether HIP-NILB-iPSCs and HIP-NILB-iPSCs-MNs evade T cell-mediated rejection. By co-culturing IFN-γ pretreated HIP-NILB-iPSCs or HIP-NILB-iPSCs-MNs with carboxyfluorescein succinimidyl ester (CFSE)-labeled PBMCs, we observed fewer proliferating T cells compared to respective controls (**Fig. 1I and fig. S2D)**, suggesting HIP-NILB-iPSCs and HIP-NILB-iPSCs-MNs immune from allogeneic T cells.

To explore whether HIP-NILB-iPSCs and HIP-NILB-iPSCs-MNs could evade innate immune attacks, we conducted immunoassays by co-culturing them with allogeneic NK cells or macrophages. As HLA-I ablation could trigger a strong ‘missing self’ killing response by NK cells, we respectively cocultured HIP-NILB-iPSCs and HIP-NILB-iPSCs-MNs with primary NK cells and measured LDH levels in the resultant co-culture medium. As expected, elevated NK cells’ cytotoxicity that incubated with KO-NILB-iPSCs (*B2M/CIITA/IL6ST/TNFR1* knock-out without *CD47/CD24/PD-L1/CTLA4-Ig* overexpression) or KO-NILB-iPSCs-MNs was compromised when incubated with HIP-NILB-iPSCs or HIP-NILB-iPSCs-MNs (**Fig. 1J and fig. S2E**), suggesting HIP-NILB-iPSCs and HIP-NILB-iPSCs-MNs evade HLA-I ablation triggered ‘missing self’ killing response by NK cells successfully. This finding was corroborated by IFN-γ secretion (**Fig. 1K and fig. S2F)** and NK cells degranulation analysis (**Fig. 1L and fig. S2G)**. Moreover, we performed the co-cultures of HIP-NILB-iPSCs with the human microglia HMC3 cells, the resident macrophages in the CNS. As indicated by LDH analysis and CD68 expression, HMC3 cells exhibited lower cytotoxicity and activation when co-cultured with HIP-NILB-iPSCs compared to controls **(fig. S2, H and I)**. A similar phagocytosis trend was observed by co-culturing HIP-NILB-iPSCs and HIP-NILB-iPSCs-MNs with macrophages derived from human monocytic cell line THP-1, respectively (**Fig. 1M and fig. S2J)**. Thus, these findings demonstrate that HIP-NILB-iPSCs and HIP-NILB-iPSCs-MNs escape both NK cell and macrophage mediated rejection.

Inhibition of *IL6ST* (*47*) and deficiency of *TNFRSF1A* (*48*) lead to defective inflammatory responses and fewer activated astrocytes upon lipopolysaccharide (LPS) administration. We thus evaluated the anti-inflammatory capacity of HIP-NILB-iPSCs. We treated them with LPS and analyses inflammatory cytokines in the supernatant. HIP-NILB-iPSCs exhibited reduced levels of inflammatory cytokines compared to NILB-iPSCs, suggesting improved resistance to an inflammatory microenvironment (**Fig. 1N**). Pathway analysis revealed enrichment of pathways associated with decreased antigen presentation, T cell cytotoxicity, type I interferon, and IFN-γ signaling in HIP-NILB-iPSCs **(fig. S2K)**. Finally, we compared the survival capacity of HIP-NILB-iPSCs containing hypoimmunogenic modifications to NILB-iPSCs with normal immunogenicity, in humanized immune system mouse model. As expected, NILB-iPSCs grafts were completely rejected within three days in allogeneic humanized-PBMCs NCG (huPBL-NCG) mice, while HIP-NILB-iPSCs survived for long term and formed teratoma at day 40 after grafting (**Fig. 1O**), confirming their superior immune tolerance *in vivo*. However, upon Dox induction, HIP-NILB-iPSCs exhibited modest and stable BLI signal levels, but no longer formed teratoma, implying their superior immune tolerance and complete differentiation, excluding the risk of tumorigenesis (**Fig. 1O**). Thus, these results along with our co-culture experiments suggest that our combined immunomodulatory strategies indeed confer superior immune tolerance capability on HIP-NILB-iPSCs.

### Survival and distribution of HIP-NILB-iPSCs-derivatives in SOD1^G93A^ pigs

A recent study has reported that poor survival of human NSCs in mouse brain could be markedly improved by immunosuppressants for both innate and adaptive immune responses (*39*). Since we have demonstrated that HIP-NILB-iPSCs can evade either innate or adaptive immune rejections, we next studied their survival and differentiation in the SOD1^G93A^ ALS pig model, which are phagocytically tolerant to human cells due to cross-species binding of porcine SIRPα to human CD47 (*49, 50*), and can more faithfully recapitulate key pathological features seen in the patients (*23, 24, 36, 37*). Four to five-year-old transgenic ALS pigs were used to determine the therapeutic effect of HIP-NILB-iPSCs. The HIP-NILB-iPSCs were pretreated with Dox for 12 hours *in vitro* to initiate differentiation. Subsequently, they were intraspinally injected into lumbar segments (L2-L5) in a 100 μL volume at three sites at a dose of 100,000 cells in 1 μL. Intraperitoneal Dox treatment for three days followed the injection of cells (**Fig. 2A**). This strategy was implemented due to the fact that long-term Dox treatment may induce toxicity in pigs, whereas the Tet-ON inducible system remains active for several days even after Dox withdrawal, allowing HIP-NILB-iPSCs to differentiate into functional MNs (*51*). It is noteworthy that we did not use any immunosuppressive drugs during the whole process of treatment.

**Fig 2.**
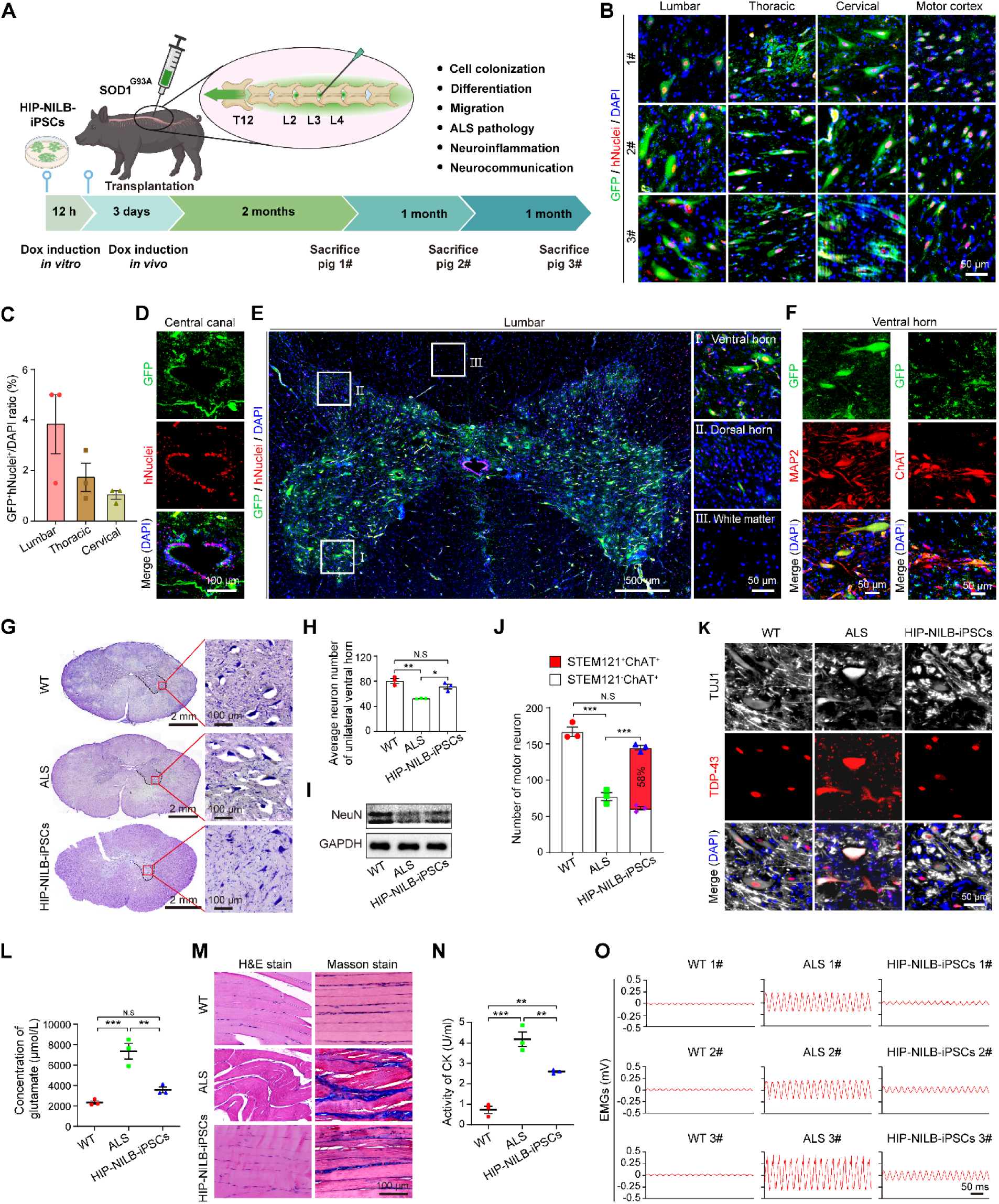
HIP-NILB-iPSCs derived MNs survived long term and ameliorate disease in SOD1^G93A^ ALS pigs. **(A)** Schematic representation of the experiment design created by BioRender. **(B)** Representative images of merged GFP and hNuclei labelling HIP-NILB-iPSCs-derivatives in the lumbar, thoracic, cervical spinal cords and motor cortex of three HIP-NILB-iPSCs treated pigs. Scale bar, 50 μm. **(C)** Quantification of the ratio of GFP^+^hNuclei^+^ cells to total cells (DAPI^+^) in the lumbar, thoracic and cervical spinal cords of three HIP-NILB-iPSCs treated pigs (n=3). **(D)** Representative images show HIP-NILB-iPSCs-derivatives surround the central canal in the lumbar spinal cord of pig-2# (3 months after grafting), which were present in all three treated pigs. Scale bar, 100 μm. **(E)** Representative images show HIP-NILB-iPSCs-derivatives predominantly localized in the ventral horn (Ⅰ) rather than in the dorsal horn (Ⅱ), and white matter (Ⅲ) of the lumbar spinal cord of pig-2#, which were present in all three treated pigs. Scale bars, 500 μm or 50 μm. **(F)** Representative immunofluorescence images show expression of MAP2 and ChAT in HIP-NILB-iPSCs-derivatives located in ventral horn of the lumbar spinal cord of pig-2#, which were present in all three treated pigs. Scale bar, 50 μm. **(G)** Nissl staining of pig lumbar spinal cord in WT, ALS and HIP-NILB-iPSCs treated pigs and magnified images for the ventral horn area. Scale bars, 2 mm or 100 μm. **(H)** Quantification of neuron number in unilateral ventral horn of WT, ALS and HIP-NILB-iPSCs treated pigs (n=3). **(I)** Western blot analysis of NeuN in the lumbar spinal cord of WT, ALS and HIP-NILB-iPSCs treated ALS pigs (n=3). **(J)** Quantitation of STEM121^+^ChAT^+^ human MNs and STEM121^-^ChAT^+^ pig MNs cell numbers in the lumbar spinal cord of WT, ALS and HIP-NILB-iPSCs treated pigs (n=3). **(K)** Representative immunofluorescence images displaying nuclear localization of TDP-43 in TUJ1^+^ neurons in HIP-NILB-iPSCs treated pig-2#, which were present in all three treated pigs. WT and ALS pigs were used as controls. Scale bar, 50 μm. **(L)** ELISA of serum glutamate in WT, ALS and HIP-NILB-iPSCs-treated pigs (n=3). **(M)** Representative H&E and Masson staining show mild atrophy and fibrosis in gastrocnemius from hindlimbs of pig-2#, which were present in all three treated pigs. WT and ALS pigs were used as controls. Scale bar, 100 μm. **(N)** Creatine kinase activity in the serum of WT, ALS and HIP-NILB-iPSCs treated pigs (n=3). **(O)** Representative EMG signals showing spontaneous activities in gastrocnemius muscle of WT, ALS and HIP-NILB-iPSCs treated pigs (n=3). Statistical analysis was performed with one-way ANOVA followed by Tukey’s multiple comparisons. Data was mean ± SEM. \**p* < 0.05, \*\**p* < 0.01, \*\*\**p* < 0.001, N.S, not significant.

To study the implantation efficiency of HIP-NILB-iPSCs after spinal injections in ALS pigs (n = 3), we sacrificed the animals at two (Pig-1#), three (Pig-2#), and four (Pig-3#) months after grafting, respectively. Strikingly, we observed numerous green fluorescent protein (GFP^+^) labeled derivatives of HIP-NILB-iPSCs in the lumbar spinal cords of all three ALS pigs (**Fig. 2B and fig. S3, A and B)**. The cellular identity was also validated by immunofluorescence staining for human nuclei (hNuclei) (**Fig. 2B and fig. S3, A and B)** and STEM121 **(fig. S3C)**, but no signals were detected in the sham control **(fig. S3D).** Previous studies have documented ectopic deposits of implanted cells along the central canal and subpial region in rat models (*52, 53*). To test whether it also occurs in ALS pigs, we detected the spread of HIP-NILB-iPSCs derivatives, which displayed widespread distributions of GFP^+^hNuclei^+^ cells in the thoracic spinal cord, cervical spinal cord, and cerebral motor cortex (**Fig. 2B and fig. S3, A and B)**, suggesting their extensive migration in the CNS of ALS pigs despite exhibiting gradually reduced ratio of HIP-NILB-iPSCs derivatives (GFP^+^hNuclei^+^ cells) along the lumbar, thoracic to cervical spinal cord, which accounted for 3.83%, 1.73% and 1.07% of all cells in respective spinal cords (**Fig. 2C**). To further determine whether they migrated into other organs, we performed PCR analysis of human-specific *Alu* gene, which revealed HIP-NILB-iPSCs-derivatives engrafted in CNS uniquely, but not in other organs **(fig. S3E)**. Mechanistically, we found that many GFP^+^hNuclei^+^ cells surrounded the central canal of the spinal cord (**Fig. 2D**), indicating they might migrate along the central canal.

Next, we characterized whether the engrafted HIP-NILB-iPSCs exhibit features of MNs. The expression of neuronal markers including TUJ1, MAP2, HB9 and ChAT was observed in the grafted GFP^+^ cells of the lumbar, thoracic, cervical spinal cord and cerebral motor cortex (**fig. S4, A to D**), suggesting Dox treatment successfully induced HIP-NILB-iPSCs into MNs *in vivo*. By quantifying the ratio of ChAT^+^STEM121^+^ and ChAT^-^STEM121^+^ cells in the lumbar spinal cord, we found 91.3% HIP-NILB-iPSCs-derivatives are MNs, with a small amount being non-MNs, which might be induced by microenvironment *in vivo* (**fig. S5A**). To characterize their lineage identity, we performed co-stains of GFP with Nestin, GFAP, Oligo2, and indeed observed that a few HIP-NILB-iPSCs differentiated into neural progenitor cells, astrocytes, or oligodendrocytes (**fig. S5, B to D**). We observed a small amount of Nestin^+^GFP^+^ cells in Pig-1# and Pig-2# (two or three months after grafting) and very few Nestin^+^GFP^+^ cells in Pig-3# (four months after grafting). As the human neuron progenitor cells have proliferation and differentiation capacities, thus, these Nestin^+^GFP^+^ cells are supposed to eventually differentiate into post-mitotic MNs, implying their proliferation would diminish with their maturation over time as we observed in the treated pigs. As expected, Ki67^+^ HIP-NILB-iPSCs-derivatives exhibited the same trend with Nestin^+^GFP^+^ cells in ALS pigs (**fig. S5E**). Moreover, we investigated the proliferation of HIP-NILB-iPSCs-derivatives *in vitro* and found no Ki67^+^ cells at day 5 upon Dox induction, which revealed their post-mitotic state (**fig. S5F**)

The neuropathological hallmarks of ALS primarily involve the degeneration of MNs in the motor cortex and spinal ventral horn (*20*). To determine whether HIP-NILB-iPSCs-MNs could directly replace the dying MNs in these regions, we examined the spatial distribution of the derivatives. As expected, a number of GFP^+^hNuclei^+^ cells were located in the motor cortex (**Fig. 2B and fig. S3, A and B)** and enriched in spinal ventral horn areas of ALS pigs (**Fig. 2E**). To confirm their maturity, we performed double immunofluorescence analysis of GFP and MNs markers, and found that the majority of GFP^+^ cells in the ventral horn areas co-expressed MAP2 and ChAT (**Fig. 2F and fig. S6, A and B)**. Given evidences from pioneer studies have documented massive loss of alpha MNs (α-MNs) and less vulnerability of gamma MNs (γ-MNs) in ALS patients and animal models (*54, 55*), this drove us to investigate whether transplanted HIP-NILB-iPSCs could differentiate into α-MNs in ALS pigs, which represents the key event for the cell replacement therapy of ALS disease. Thus, we performed co-stains of ERR3, NeuN with GFP, which showed that HIP-NILB-iPSCs selectively differentiated into α-MNs (NeuN^+^ERR3^-^) rather than γ-MNs (NeuN^-^ERR3^+^) in ALS pigs **(fig. S6C)**. This finding was corroborated by the transcriptomic results of HIP-NILB-iPSCs differentiated MNs upon Dox induction for 7 days in vitro **(fig. S6D)**. Therefore, these findings demonstrate that HIP-NILB-iPSCs could differentiate into α-MNs that are functional for treating ALS, survive for long term and migrate to the defective regions in ALS pigs.

### HIP-NILB-iPSCs treatment remarkably ameliorates disease in SOD1^G93A^ pigs

To address whether HIP-NILB-iPSCs-MNs exert impacts on disease progression of ALS, we conducted a series of experiments to examine the typical pathological features in ALS pigs treated with HIP-NILB-iPSCs, untreated ALS pigs and WT pigs. As revealed by Nissl staining, we found a significant increase of neurons in the ventral horn of the spinal cord in HIP-NILB-iPSCs treated ALS pigs, compared to untreated ALS pigs (**Fig. 2G and H**). Western blot analysis showed the elevated protein levels of the neuron marker NeuN in treated pigs (**Fig. 2I**). Consistently, co-stains of human STEM121 with ChAT antibody displayed that substantial decrease of host MN number happened no matter in HIP-NILB-iPSCs-treated or in untreated ALS pigs. However, the human stem cell-derived MNs (STEM121^+^ChAT^+^) almost fully made up for loss of host ChAT^+^ MNs, accounting for about 58% of total ChAT^+^ MNs in spinal cords of the HIP-NILB-iPSCs-treated ALS pigs (**Fig. 2J**). We then detected the location of TDP-43 in MNs, another major pathological hallmark of ALS, which leads to a loss of function in the nucleus and a gain of toxicity in the cytoplasm. TDP-43 was mainly distributed in the nuclei in HIP-NILB-iPSCs treated ALS pigs and WT pigs, whereas it was located in the cytoplasm of neurons in untreated ALS pigs (**Fig. 2K and fig. S7A)**. Additionally, we observed decreased serum glutamate levels, a neurotoxic amino acid associated with ALS, indicating the improved functionality of MNs in HIP-NILB-iPSCs treated ALS pigs (**Fig. 2L**). To address whether the restoration of neurons affects muscles, we performed H&E and Masson staining and found a remarkable attenuation of gastrocnemius muscle atrophy and fibrosis in HIP-NILB-iPSCs treated ALS pigs (**Fig. 2M**). The reduced serum creatine kinase (CK) levels also confirmed the diminished muscle injury (**Fig. 2N**). Consistently, the electromyography (EMG) signals revealed a significant improvement of the gastrocnemius muscle fibrillation in the treated ALS pigs, compared to untreated ALS pigs (**Fig. 2O**), indicating a recovery of hindlimb muscle denervation upon treatment with HIP-NILB-iPSCs.

To investigate whether axons of HIP-NILB-iPSC-MNs extend their long peripheral projection to the gastrocnemius muscle of ALS pigs, co-staining of α-BTX, NF-H and GFP was employed to examine the neuromuscular junctions (NMJs). Strikingly, the NMJs composed of donor axonal terminal of HIP-NILB-iPSCs-MNs (NF-H^+^) and postsynaptic receptors (α-BTX^+^) on host muscle fiber were observed in pig-2# and pig-3# (3 and 4 months after HIP-NILB-iPSCs grafting) (**Fig. 3A**). Of note, we did not see any reformed NMJs in pig-1# (2 months after HIP-NILB-iPSCs grafting), which might be reasoned by inadequate grafting time for extension of axon projection. Thus, these findings provided convincible evidences for that HIP-NILB-iPSCs-MNs were able to project their long-distance axons to the peripheral gastrocnemius muscle and participate in reformation of NMJs in ALS pigs, implying functional integration of HIP-NILB-iPSCs-MNs ameliorated ALS disease.

**Fig 3.**
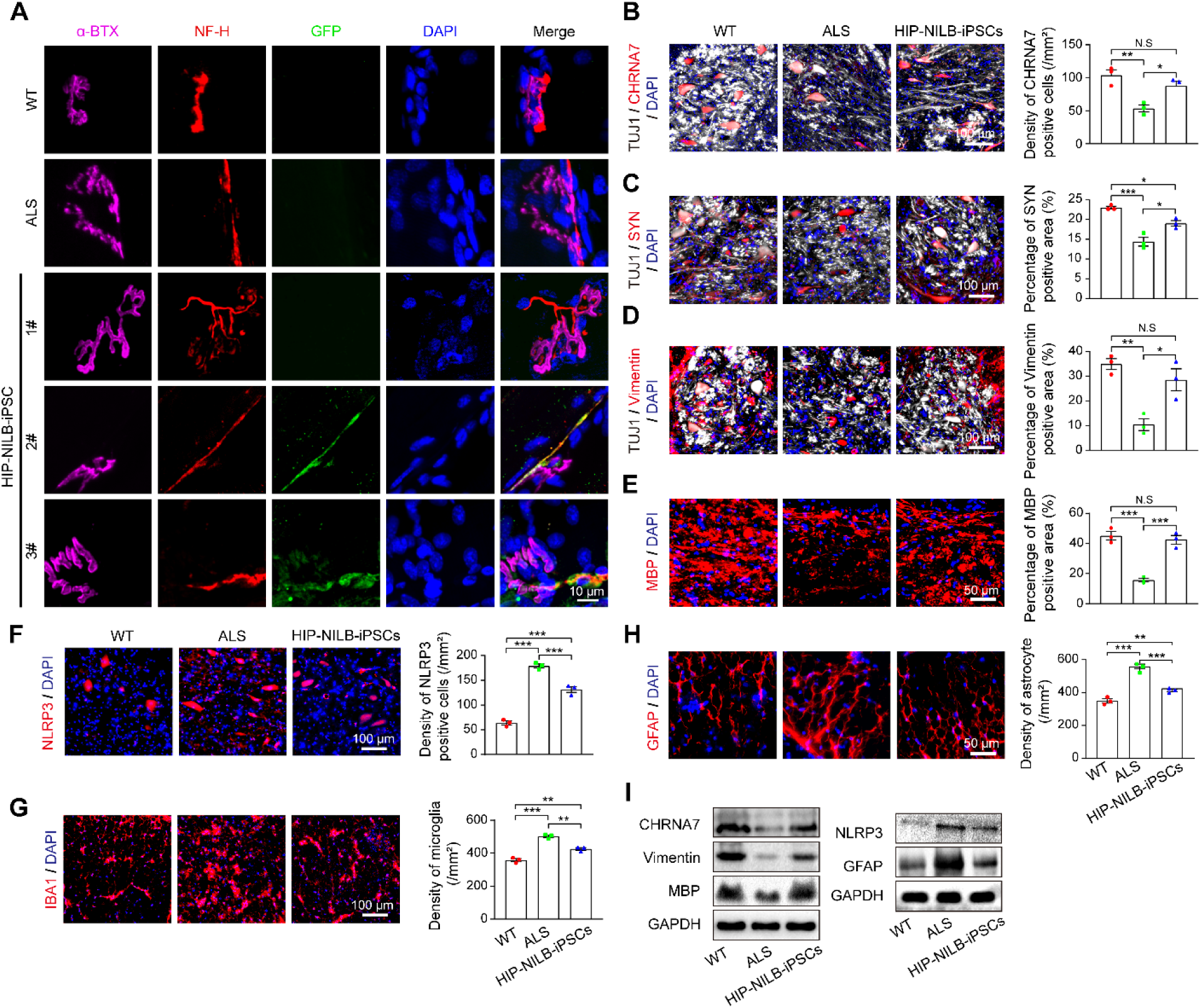
HIP-NILB-iPSCs derived MNs integrated into neural circuits and attenuated neuron degeneration and neuroinflammation in SOD1^G93A^ ALS pigs. **(A)** Representative immunofluorescence images of a-BTX/NF-H/GFP/DAPI showing reformed neuromuscular junctions between HIP-NILB-iPSCs-derived MNs and host gastrocnemius muscle fiber in Pig-2# and Pig-3# (three and four months after grafting), but not in Pig-1# (two months after grafting), WT or ALS controls. Scale bar, 10 μm. **(B-H)** Representative immunofluorescence images for (B) CHRNA7, (C) SYN, (D) Vimentin, (E) MBP, (F) NLRP3, (G) IBA1 and (H) GFAP in the lumbar spinal cord of WT, ALS and HIP-NILB-iPSCs treated pigs, and their quantifications (n=3). Scale bars, 100 μm or 50 μm. **(I)** Western blot analysis of CHRNA7, Vimentin, MBP, NLRP3 and GFAP in lumbar spine cord of WT, ALS and HIP-NILB-iPSCs treated pigs. The statistical analysis was performed by one-way ANOVA following Tukey’s multiple comparisons. Data was shown as mean ± SEM. \**p* < 0.05, \*\**p* < 0.01, \*\*\**p* < 0.001, N.S, not significant.

Next, we attempted to underline the efficacy of treatment by examining molecular changes related to neuron regeneration and neuroinflammation. To investigate whether the increased number of neurons observed in the spinal cord would support functional neuro-communication, we performed double immunofluorescence staining of TUJ1 with cholinergic receptor nicotinic alpha 7 subunit (CHRNA7), a neuronal acetylcholine receptor modulating neurotransmission, and synaptophysin (SYN), a neuronal synaptic vesicle regulating neurotransmitter release. The expression of CHRNA7 and SYN exhibited a significant increase in the spinal cord of HIP-NILB-iPSCs treated ALS pigs compared to untreated ALS pigs (**Fig. 3, B and C and fig. S7, B and C)**, suggesting improved neurotransmission, synaptogenesis upon HIP-NILB-iPSCs treatment. Moreover, we examined the expression of vimentin, a marker for neural stem cells, and myelin basic protein (MBP), a marker for stabilizing and compacting myelin, which showed elevated expression (**Fig. 3, D and E and fig. S7, D and E)**, and further validated enhanced neuron regeneration and functional neurocommunication in HIP-NILB-iPSCs treated ALS pigs. Furthermore, we evaluated the neuroinflammation microenvironment by examining the expression of key neuroinflammatory modulators, including inflammasome marker NLRP3, microglia marker IBA1, and astrocyte marker GFAP. HIP-NILB-iPSCs treated ALS pigs exhibited lower expressions of NLRP3, IBA1, and GFAP (**Fig. 3, F to H and fig. S7, F and G)**, suggesting that HIP-NILB-iPSCs’ treatment mitigates the inflammatory environment rather than exacerbates it. Consistently, the changes of the above markers were further confirmed using western blot analysis (**Fig. 3I**). Lastly, we observed comparable decreased density of CD86^+^ M1 pro-inflammatory microglia and C3^+^ A1 neurotoxic reactive astrocyte in HIP-NILB-iPSCs-treated ALS pigs compared with untreated ALS pigs (**fig. S8, A and B**), while the density of CD206^+^ M2 anti-inflammatory microglia and S100A10^+^ A2 neuroprotective astrocyte increased (**fig. S8, C and D**), further validating improved microenvironment in HIP-NILB-iPSCs-treated ALS pigs.

### HIP-NILB-iPSCs-MNs demonstrate long-term survival and integration into neural circuits in TIA1P362L rabbits

To extend the therapeutic application of HIP-NILB-iPSCs to other ALS models and gain convincing insights into their clinical translation, we next treated TIA1^P362L^ ALS rabbits with HIP-NILB-iPSCs (*56*). These rabbit models precisely mimic the ALS causative TIA1^P362L^ point mutation in ALS patients, avoiding variations of transgene expression in SOD1^G93A^ transgenic models, and exhibit characteristic symptoms seen in ALS patients (*57*), including loss of MNs, cytoplasmic TDP-43 aggregation, abnormal electromyographic activity, muscular dystrophy, and even motor deficits. To minimize individual therapeutic differences upon HIP-NILB-iPSCs transplantation in ALS rabbits, we selected six TIA1^P362L^ ALS rabbits at ages of 8-10 months, which clearly manifested movement deficits (**Fig. 4A**). Following the above-optimized delivery procedure, we intraspinally transplanted 4,000,000 HIP-NILB-iPSCs into each ALS rabbit without using immunosuppression. As in stem cell-treated ALS pigs, we observed numerous GFP^+^hNuclei^+^ cells in the lumbar cord, as well as the thoracic cord, the cervical and cerebral motor cortex of ALS rabbits at two months after grafting (**Fig. 4B and fig. S9A)**, but no signals were detected in the sham control **(fig. S9B)**. The immunofluorescence staining for STEM121 **(fig. S9C)** and PCR analysis of human-specific *Alu* gene (**Fig. 4C**) further validated the cellular identity and engraftment in CNS. The ratio of GFP^+^hNuclei^+^ cells accounted for 7.47%, 3.70% and 2.13% of total cell numbers in the lumbar, thoracic and cervical spinal cord of stem cell-treated ALS rabbits, respectively (**Fig. 4D**), indicating successful engraftment and migration of HIP-NILB-iPSCs-derivatives in CNS of ALS rabbits. Enrichments of GFP^+^hNuclei^+^ cells were observed in the central canal (**Fig. 4E**) and ventral horn of the lumbar spinal cord (**Fig. 4F**), suggesting their migration to defective regions, and potential replacement of degenerating MNs in ALS rabbits. In contrast, when NILB-iPSCs with intact immunogenicity were transplanted into ALS rabbits, no or very few human cells survived in lumbar spinal cord of ALS rabbits at one or two months after grafting **(fig. S9D)**, confirming the necessity of hypoimmunogenic modification for the long-term survival and thereby precluding us from further functional investigations of NILB-iPSCs treated rabbits.

**Fig 4.**
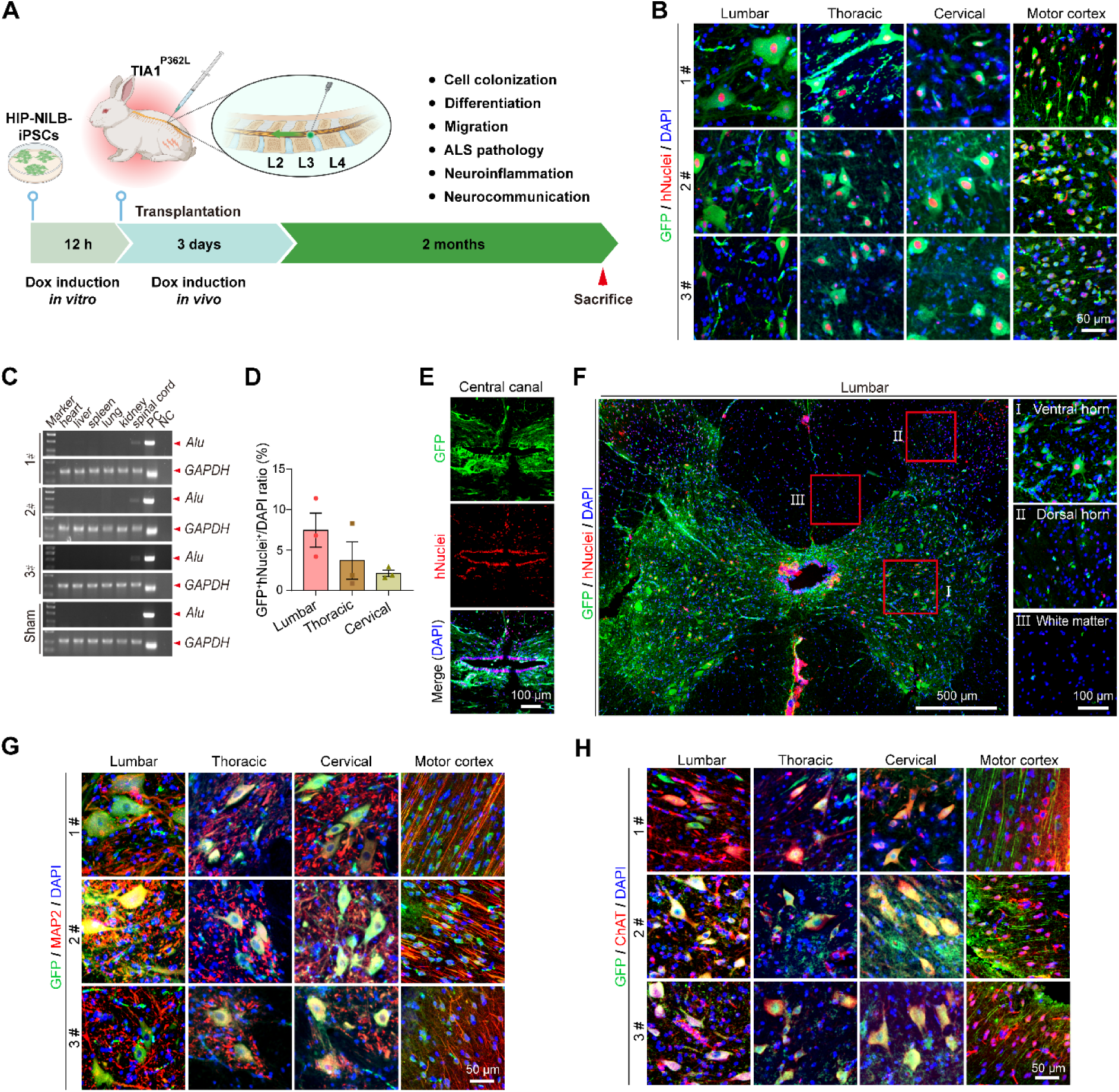
HIP-NILB-iPSCs derived MNs survived long term in TIA1^P362L^ rabbits. **(A)** Schematic representation of the experiment design created by BioRender. **(B)** Representative images of merged GFP and hNuclei labelling HIP-NILB-iPSCs-derivatives in the lumbar, thoracic, cervical spinal cords and motor cortex of three HIP-NILB-iPSCs treated rabbits. Scale bar, 50 μm. **(C)** Human specific *Alu* genome PCR analysis of various organs and spinal cords of three HIP-NILB-iPSCs treated rabbits and the sham control. **(D)** Quantification of the ratio of GFP^+^hNuclei^+^ cells to total cells (DAPI) in the lumbar, thoracic and cervical spinal cords of HIP-NILB-iPSCs treated rabbits (n=3). **(E)** Representative immunofluorescence images show HIP-NILB-iPSCs-derivatives aggregates around the central canal of lumbar spinal cord of HIP-NILB-iPSCs treated rabbits. Scale bar, 100 μm. **(F)** Representative images show HIP-NILB-iPSCs-derivatives predominantly localize in the ventral horn (I) rather than in the dorsal horn (Ⅱ) and white matter (Ⅲ), of lumbar spinal cord of HIP-NILB-iPSCs treated rabbits. Scale bars, 500μm or 100 μm. **(G-H)**Representative immunofluorescence images for (G) MAP2 and (H) ChAT in the lumbar, thoracic, cervical spine and motor cortex of three HIP-NILB-iPSCs treated rabbits. Scale bar, 50 μm.

To determine whether these HIP-NILB-iPSCs matured into functional MNs, we examined the expression of MNs markers in these HIP-NILB-iPSCs-derivatives. Indeed, these GFP^+^ cells co-expressed MAP2 and ChAT (**Fig. 4, G and H, and fig. S10, A and B)**. In line with lineage identities of HIP-NILB-iPSCs-derivatives in ALS pigs, we observed 92% HIP-NILB-iPSCs-derivatives are MNs **(fig. S11A)**, with only a few cells are GFAP^+^ astrocytes, Oligo2^+^ oligodendrocytes **(fig. S11, B and C)**. Of note, no Nestin^+^ or Ki67^+^ HIP-NILB-iPSCs-derivatives were found in HIP-NILB-iPSCs-treated ALS rabbits **(fig. S11, D and E)**, suggesting their post-mitotic state and maturation. Consistently, we examined subtypes of HIP-NILB-iPSCs-MNs, and found that most of them are α-MNs (NeuN^+^ERR3^-^) but not γ-MNs (NeuN^-^ERR3^+^) **(fig. S11F)**, implying their replacement of degenerating α-MNs in ALS rabbits.

To define whether direct connections have been formed between donor and host neurons, we infected HIP-NILB-iPSCs with LV-EF1a:G and subsequently transplanted them into ALS rabbits together with rabies-dG-tdTomato using the optimized procedure. At fourteen days post-transplantation, we observed the transport of dense tdTomato-labeled rabies virus to neighboring host neurons (GFP^-^tdTomato^+^ cells indicated by white arrow), indicating that HIP-NILB-iPSC-MNs received afferent projections from the host neurons (**Fig. 5A**).

**Fig 5.**
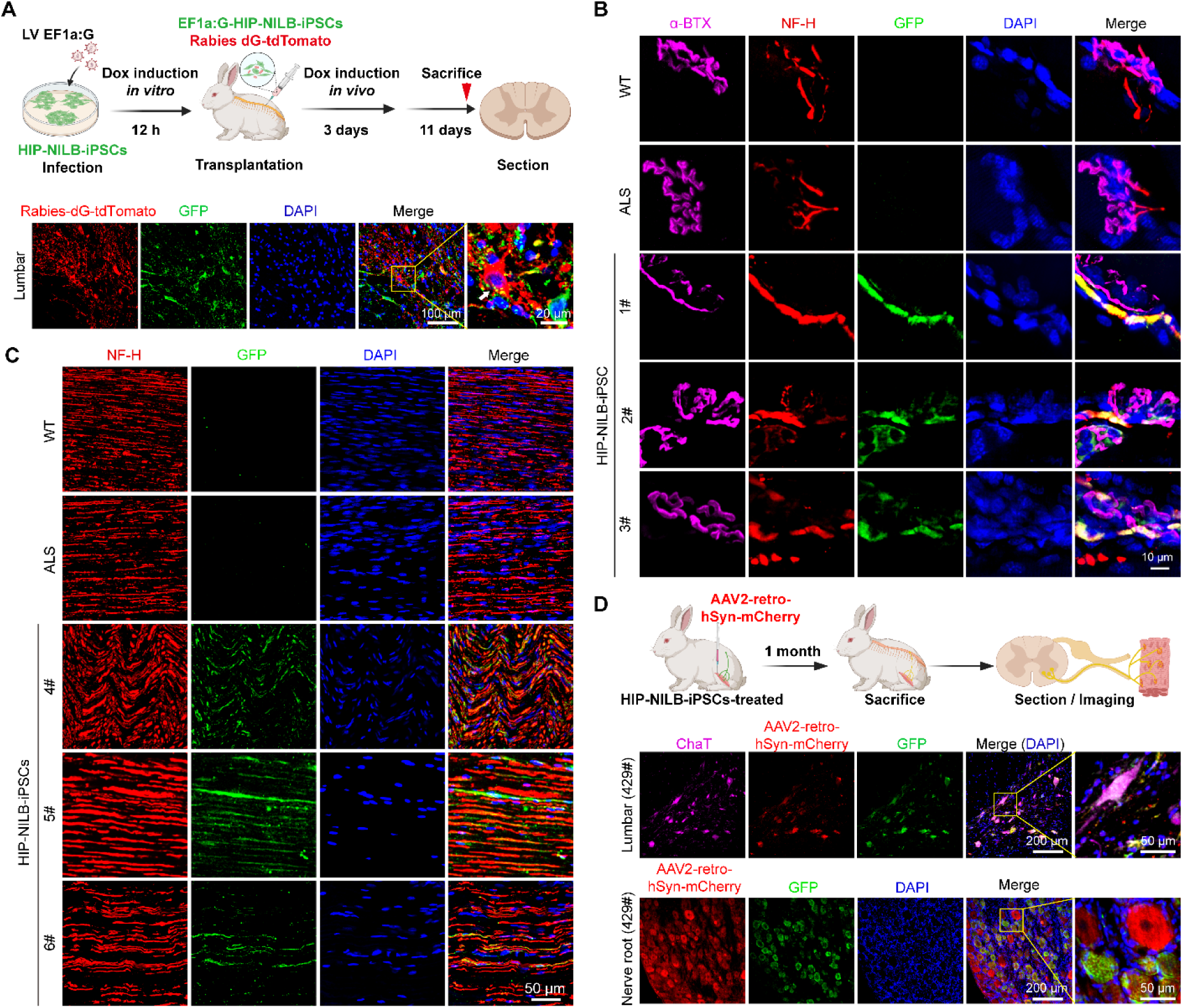
HIP-NILB-iPSCs-derived MNs integrated into neural circuits in TIA1^P362L^ ALS rabbits. **(A)** Scheme and representative images for rabies-virus-based monosynaptic tracing show retrograde labeling of inputs from HIP-NILB-iPSCs-MNs to host neurons (n=3). The schematic diagram created by BioRender. The host neuron was pointed out with a white arrow in the framed region. Scale bars, 100 μm or 20 μm. **(B)** Representative immunofluorescence images of α-BTX^+^NF-H^+^GFP^+^DAPI^+^ showing reformed NMJs between HIP-NILB-iPSCs-derived MNs and host gastrocnemius muscle fibers in all three HIP-NILB-iPSCs treated rabbits (Rabbit 1#, 2# and 3#), but not in WT or ALS controls. Scale bar, 10 μm. **(C)** Representative immunofluorescence images of NF-H^+^GFP^+^DAPI^+^ showing reformed neurofilament of HIP-NILB-iPSCs-MNs within sciatic nerve in HIP-NILB-iPSCs treated rabbits, but not in WT or ALS controls. Scale bar, 50 μm. **(D)** Scheme and representative images of retrograde tracing using AAV2-retro-hSyn-mcherry-virus from the gastrocnemius muscle to the spinal cord show labeling of both HIP-NILB-iPSCs-MNs (mCherry^+^ChAT^+^GFP^+^) in the lumbar spinal cord and axons (mCherry^+^GFP^+^) in spinal nerve root of HIP-NILB-iPSCs treated rabbits (n=3). The schematic diagram created by BioRender. Scale bars, 200 μm or 50 μm.

To investigate whether axon of HIP-NILB-iPSCs-MNs reached muscle fibers, we performed immunofluorescence staining on gastrocnemius muscles from HIP-NILB-iPSCs treated rabbits (Rabbit 1^#^, 2^#^ and 3^#^). The results revealed the reformation of NMJs between donor axon terminals from HIP-NILB-iPSCs-derived MNs and postsynaptic receptors on host muscle fibers in the gastrocnemius muscles at two months after grafting (**Fig. 5B**). Of note, we did not collect sciatic nerve tissues of above three treated ALS rabbits. To further verify that HIP-NILB-iPSCs-MNs indeed projected out of vertebral canal, we treated three more ALS rabbits (Rabbit 4^#^, 5^#^ and 6^#^) and collected their sciatic nerve tissues at more than three months later after grafting and stained them with NF-H and GFP antibodies, confirming that human neurofilaments existed in sciatic nerve of HIP-NILB-iPSCs-treated ALS rabbits (**Fig. 5C**). Similarly, we observed the reformation of new NMJs in the gastrocnemius muscles of these treated ALS rabbits **(fig. S12A)**. Additionally, we injected AAV2-retro-hSyn-mCherry viruses into gastrocnemius muscle of ALS rabbits at two months after HIP-NILB-iPSCs grafting to trace lower MNs projecting to gastrocnemius muscle.(*58, 59*) One month after AAV2-retro-hSyn-mCherry virus injection, we observed that many ChAT^+^mCherry^+^GFP^+^ cells were located in the ventral horn of lumbar spinal cord (**Fig. 5D and fig. S12, B and C)**, and mCherry^+^GFP^+^ axons existed in the lumbar nerve roots (**Fig. 5D**). In summary, our data demonstrate that HIP-NILB-iPSC-MNs survive long term and integrate into neural circuits in ALS rabbits.

### Rescue of TIA1P362L ALS rabbits by HIP-NILB-iPSCs treatment

Next, we compared the pathological features between HIP-NILB-iPSCs treated or untreated ALS rabbits, and WT rabbits, which revealed a significant increase in the number of neurons in the ventral horn of the spinal cord in the HIP-NILB-iPSCs treated ALS rabbits (**Fig. 6, A and B)**. Additionally, the treated rabbits exhibited a higher gray-to-white matter ratio compared to the untreated ALS rabbits (**Fig. 6B**). This increase was further confirmed by western blot analysis, which showed an elevation in NeuN expression (**Fig. 6C**). By quantifying ChAT^+^ MNs, substantial decrease of host MN number was observed no matter in HIP-NILB-iPSCs-treated or in untreated ALS rabbits. However, the human stem cell-derived MNs (STEM121^+^ChAT^+^) almost fully made up for loss of host ChAT^+^ MNs, accounting for about 40.6% of total ChAT^+^ MNs in spinal cords of the HIP-NILB-iPSCs-treated ALS rabbits (**Fig. 6D**). The restoration of neurons was corroborated by the analysis of TDP-43 pathology (**Fig. 6E and fig. S13A**). The insoluble TDP-43 in the pellet of HIP-NILB-iPSCs-treated ALS rabbits also significantly decreased, accompanied by reduced C-terminal fragments CTF35 and CTF25, two cleaved fragments of TDP-43 that provide seeds for aggregation of the full-length TDP-43 (**Fig. 6F**). Furthermore, decreased glutamate levels (**Fig. 6G**), as well as alleviated muscle injury indicated by H&E staining, Masson staining, and CK assays (**Fig. 6, H and I**), were observed in the HIP-NILB-iPSCs treated ALS rabbits.

**Fig 6.**
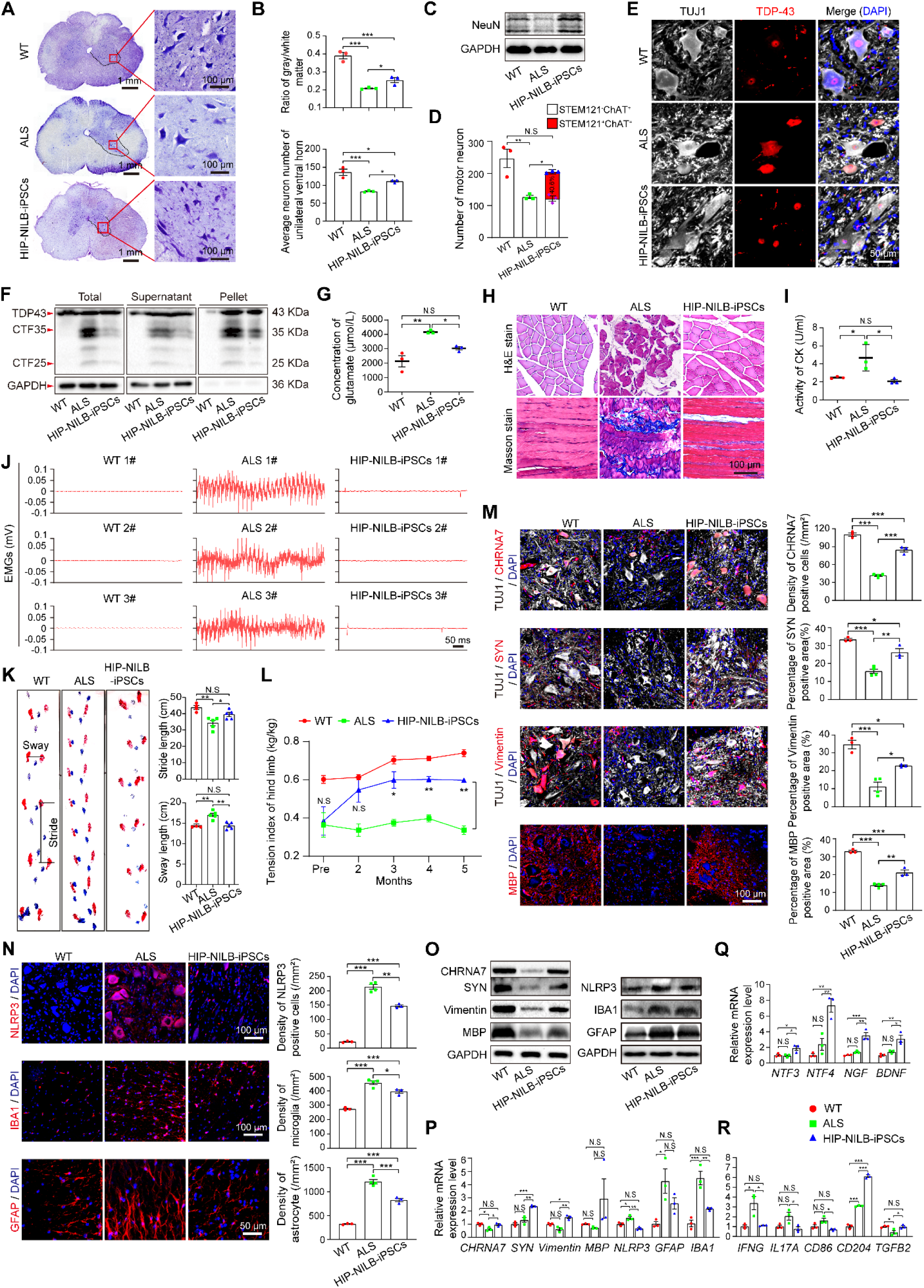
Rescue of TIA1^P362L^ ALS rabbits by HIP-NILB-iPSCs derived MNs. **(A)** Nissl staining of lumbar spinal cord in WT, ALS and HIP-NILB-iPSCs treated rabbits, and representative magnified images for ventral horn area (n=3). Scale bars, 1 mm or 100 μm. **(B)** Quantifications of gray to white matter ratio and average neuron number in the unilateral ventral horn (n=3). **(C)** Western blot analysis of NeuN expression in spinal cords of WT, ALS and HIP-NILB-iPSCs treated rabbits (n=3). **(D)** Quantitation of STEM121^+^ChAT^+^ human MNs and STEM121^-^ChAT^+^ rabbit MNs cell numbers in the lumbar spinal cord of three HIP-NILB-iPSCs treated rabbits. WT and ALS rabbits were used as controls. **(E)** Representative immunofluorescence images displaying nuclear localization of TDP-43 in TUJ1^+^ neurons in HIP-NILB-iPSCs treated. WT and ALS rabbits were used as controls. Scale bar, 50 μm. **(F)** Western blot analysis of the soluble and insoluble TDP-43 in supernatants and pellets of rabbit lumbar spinal cords. **(G)** ELISA assay for serum glutamate concentration in WT, ALS and HIP-NILB-iPSCs treated rabbits (n=3). **(H)** Representative H&E and Masson staining images showing the atrophy and fibrosis of gastrocnemius muscle in WT, ALS and HIP-NILB-iPSCs treated rabbits (n=3). Scale bar, 100 μm. **(I)** Creatine kinase activity in the serum of WT, ALS and HIP-NILB-iPSCs treated rabbits (n=3). **(J)** Representative EMG signals showing spontaneous activities in gastrocnemius muscle of WT, ALS and HIP-NILB-iPSCs treated rabbits. **(K)** Gait analysis and quantification of stride and sway length in hind limbs of WT (n=4), ALS (n=5) and HIP-NILB-iPSCs treated rabbits (n=5) at two months after grafting. **(L)** Analysis of tension index of hind limbs over time in WT (n=4), ALS (n=3) and HIP-NILB-iPSCs treated rabbits (n=3). The statistics analysis was performed with two-way ANOVA followed by Tukey’s multiple comparisons, and the significant differences were labeled between ALS and HIP-NILB-iPSCs group in different time points. **(M)** Representative immunofluorescence staining of CHRNA7, SYN, Vimentin and MBP in lumbar spinal cord of WT (n=3), ALS (n=4) and HIP-NILB-iPSCs treated rabbits (n=3), and their quantifications. Scale bar, 100 μm. **(N)** Representative immunofluorescence staining of NLRP3, IBA1 and GFAP in lumbar spinal cord of WT (n=3), ALS (n=4) and HIP-NILB-iPSCs treated rabbits (n=3), and their quantifications. Scale bars, 100 μm or 50 μm. **(O-P)** (O) Western blot analysis and (P) Real-time PCR analysis of CHRNA7, SYN, Vimentin, MBP, NLRP3, IBA1, GFAP in lumbar spinal cords of WT (n=3), ALS (n=3) and HIP-NILB-iPSCs treated rabbits (n=3). **(Q-R)** Real-time PCR analysis of (Q) neurotrophic factors (*NTF3*, *NTF4*, *NGF*, *BDNF*) and (R) inflammatory factors (*IFNG*, *IL17A*, *CD86*, *CD204* and *TGFB2*) in lumbar spinal cords of WT (n=3), ALS (n=3) and HIP-NILB-iPSCs treated rabbits (n=3). The statistical data were analyzed by one-way ANOVA followed by Dunnett’s T3 multiple comparisons (N-NLRP3) or Tukey’s multiple comparisons (b, d, g, i, k, m, n-IBA1, n-GFAP, p, q, r) unless mentioned otherwise. Data was mean ± SEM. \**p* < 0.05, \*\**p* < 0.01, \*\*\**p* < 0.001, N.S, not significant.

To evaluate the functional integration of HIP-NILB-iPSCs-MNs into neural host circuits in ALS rabbits, we assessed the gastrocnemius muscle fibrillation by recording EMG signals. The HIP-NILB-iPSCs-treated ALS rabbits displayed a significant improvement in muscle fibrillation compared to untreated ALS rabbits (**Fig. 6J**), indicating recovery of hindlimb muscle denervation. More importantly, we observed a remarkable alleviation of movement deficits, including increased stride length and decreased sway length, in HIP-NILB-iPSCs-treated ALS rabbits, compared to ALS rabbits (**Fig. 6K**). Moreover, we examined muscle strength of hind limbs in the rabbit model over time, which revealed gradual improvements of HIP-NILB-iPSCs-treated ALS rabbits compared to untreated ALS rabbits (**Fig. 6L**).

Finally, we examined the molecular signature changes associated with neuropathological features in HIP-NILB-iPSCs-treated ALS rabbits. In accordance with improved synaptogenesis and neuron regeneration observed in HIP-NILB-iPSC-treated ALS pigs, we found increased expression of CHRNA7, SYN, MBP, and Vimentin in the treated ALS rabbits, indicating enhanced neurotransmission and neuron regeneration (**Fig. 6M and fig. S13, B to E)**. The HIP-NILB-iPSCs treated rabbits exhibited reduced neuroinflammation, as evidenced by lower expression levels of IBA1, GFAP, and NLRP3 (**Fig. 6N and fig. S13, F and G)**. This finding was further corroborated by reduced density of CD86^+^ M1 pro-inflammatory microglia and C3^+^ A1 neurotoxic reactive astrocyte **(fig. S14, A and B)**, and increased density of CD206^+^ M2 anti-inflammatory microglia and S100A10^+^ A2 neuroprotective astrocyte **(fig. S14, C and D)**. The changes in these markers were further validated by western blot and qRT-PCR analysis (**Fig. 6, O and P)**. Furthermore, qRT-PCR analysis of multiple neurotrophic factors, including *NTF3, NTF4, NGF, BDNF,* and neuron inflammation factors, including *IFNG, IL17A, CD86, CD204* and *TGFB2*, confirmed improved neurotrophy and decreased inflammation response in the HIP-NILB-iPSCs treated ALS rabbits (**Fig. 6, Q and R**). Overall, these findings demonstrate that long-term survival of hypoimmunogenic human MNs successfully rescue the pathological phenotype in TIA1^P362L^ rabbits.

## DISCUSSION

Over the past twenty years, extensive studies have attempted to replace degenerated MNs in ALS hosts through cell therapy. Despite some studies reporting mild benefits through modulating the host microenvironment (*60*) or repairing pathological glial cells (*1, 61*), the overall outcomes remained poor, which could be attributed to the vigorous immune rejection of the host, limited differentiation of transplanted cells into functional MNs, especially the failure of their long projections to muscle targets. Here, we present a novel approach that engineers human iPSCs to be hypoimmunogenic and inducible into MNs *in vivo*, which also represent ‘off-the-shelf’ universal cells, as they could overcome batch variability. More importantly, we demonstrate for the first time that hypoimmunogenic human MNs can survive long term without using any immunosuppression, effectively replenish dying MNs, integrate into host neural circuits, and substantially reduce disease burden in large animal models of ALS.

The utilization of large animal models in clinical translations is of utmost importance due to their similarities to humans in terms of CNS anatomy and life expectance (*38, 62*). In our study, the SOD1^G93A^ pigs were developed previously by transferring mutant human *SOD1* genes into the pig genome (*23*), while the TIA1^P362L^ rabbit models faithfully mimic human ALS causative TIA1^P362L^ missense mutation. Both animal models exhibit key pathological features that does not appear in small rodents, including MNs loss, TDP-43 cytoplasm mislocalization. ALS pigs did not exhibit noticeable movement deficits like the F0 generation, possibly due to variations in transgene copies during breeding, while the rabbit models showed crippled movement across different individual offspring, which allow us to evaluate the treatment effects on motor deficits. Here, we achieved consistent treatment effects between the two different species of animal models, despite their size differences, and among all treated ALS animals (three ALS pigs and twelve ALS rabbits), such as long-term survival, widespread migration and efficient differentiation into functional α-MNs of the grafts, reformation of new NMJs, and remarkable amelioration of ALS pathologies. Hence, the low variabilities in animal transplantations makes us believe that when our approach is translated to clinical trials for human ALS patients, similar outcomes should be achieved.

Previous studies have demonstrated immune tolerance of hypoimmunogenic human iPSCs (*30, 35, 63, 64*) and embryonic stem cells (*19*) in allogenic or xenogeneic settings. But these cells have been engineered limited immune-related genes and remained elusive in neurodegenerative diseases therapy. To tackle the challenges of immune rejection and neuroinflammation in ALS hosts, we implemented a series of genomic alterations to evade both adaptive and innate immune attacks including T cell-, NK-, and microphage-mediated immune rejection, the ‘missing self’ killing response, as well as the inflammatory environment. As a result, these multiple modified human iPSC can survive long term after injecting into the spinal cord of pig and rabbit models without the use of any immunosuppression. In addition, it must be mentioned that the long-term persistence of HIP-NILB-iPSCs-MNs in xenogeneic ALS large animal models may rather be underestimated and possibly leads to more benefits when tested in allogeneic hosts, i.e. human patients.

In our study, we deeply investigated whether the integration of human stem cell-derived MNs or the trophic effects of transplants result in amelioration of ALS disease. When HIP-NILB-iPSCs infected with LV-EF1a:G were transplanted into ALS rabbits together with rabies-dG-tdTomato, the transport of dense tdTomato-labeled rabies virus to neighboring host neurons was observed. Co-staining for NF-H and GFP revealed the presence of human neurofilaments in the sciatic nerve of ALS rabbits treated with HIP-NILB-iPSCs. Using AAV2-retro-hSyn-mCherry viruses for retrograde tracing, we observed mCherry^+^GFP^+^ axons in lumbar nerve roots, as well as ChAT^+^mCherry^+^GFP^+^STEM121^+^ cells in the lumbar spinal cord. These findings indicated that axons of human MNs projected out of the vertebral canal and extended toward the gastrocnemius muscles. Fascinatingly, co-staining for α-BTX, NF-H, and GFP in host muscle tissues revealed NMJs formed between human axon terminals and host muscle fibers in the gastrocnemius muscles of both HIP-NILB-iPSCs-treated ALS pigs and rabbits. A thorough analysis of the lineage identities of HIP-NILB-iPSCs derivatives showed that the majority of human stem cell differentiated into MNs. Transcriptome profiling of HIP-NILB-iPSCs-derived MNs induced *in vitro* demonstrated that expression of α-MN-specific genes was elevated, and most transplants differentiated into α-MNs *in vivo*. Furthermore, axon regeneration rate varies from 1-2 mm to 5-8 mm per day with various lesion types, neuron types and species (*65–67*). Here, due to MNs fast differentiation by forced transcription factors expression, *in vivo* neural microenvironment, pig and rabbit species-factors, and ALS disease situation, the MN axons were successfully extended to rabbit and pig gastrocnemius muscle in 2 months and 3 months, respectively, suggesting a high axon growth rate could be achievable in adult animals *in vivo*. Despite their underlying molecular and cellular mechanisms not well documented in this study, we firstly reported newly formed NMJs, indicating successful cell replacement.

We also identified a small quantity of non-MN-cells derived from the transplant, such as astrocytes and oligodendrocyte-lineage cells. These non-MN cells may provide trophic support, improving the neurogenic microenvironment and potentially contributing to the attenuation of ALS phenotypes, as indicated in prior studies (*1, 68, 69*). However, since most transplants differentiated into MNs, survived long-term, and extended axons to the gastrocnemius muscles, direct cell replacement should play a major role in ameliorating ALS symptoms in this study.

Concerns regarding the safety of *in vivo* differentiation of stem cells induced with Dox and the intensive genetic modifications of cells may influence the translatability of our approach. Since the HIP-NILB-iPSCs were pretreated with Dox for 12 hours *in vitro* prior to transplantation, which enables expression of Tet-ON induced genes for several days even after Dox withdrawal (*51*), all cells were capable of responding to Dox and initiating their differentiation. Regarding the proliferation status of stem cell-derivatives post-Dox induction, we did not observe Ki67^+^ human cells *in vitro* and in CNS of HIP-NILB-iPSCs-treated rabbits. Although some Ki67^+^ cells were present in stem cell-treated ALS pigs, their numbers gradually decreased over time post-grafting, mirroring the decline of human neural progenitor cells. Therefore, the identified Ki67^+^ cells likely represent human neural progenitor cells, which possess proliferative and differentiating capabilities and are expected to eventually differentiate into post-mitotic MNs. This implies that the proliferation of HIP-NILB-iPSCs derivatives would diminish over time as they mature into neurons.

Furthermore, no tumorigenesis was observed in any of the HIP-NILB-iPSCs-treated ALS models or in humanized immune system mice. Consequently, the risk of tumorigenesis arising from incomplete differentiation or cancerous mutations in transplanted stem cells appears to be very low. Alternatively, incorporating designed suicide-switch systems into HIP-NILB-iPSCs could provide an additional safeguard against safety concerns. These suicide-switch systems could enable stringent control over cell survival, facilitating the selective elimination of undifferentiated PSCs or the complete removal of all proliferating PSC-derivatives when necessary (*70*). In summary, our results suggest that HIP-NILB-iPSCs delivery and MNs replacement offer a viable and broadly applicable strategy toward an efficient cell therapy of ALS or other MNs degenerative diseases.

## MATERIALS AND METHODS

### Animal assurance

All animal experiments were performed according to the regulations of the Animal Care and Use Committee of the Guangzhou Institutes of Biomedicine and Health (GIBH) (IACUC2017014, IACUC2022034). The parents donated the umbilical cord blood of their child to generate the UH10 iPSCs and provided permission to use it for biomedical research. This study was approved by Human Subject Research Ethics Committee/Institutional Review Board of GIBH (GIBH-IR807-2017014, GIBH-IRB2022-022) and was performed in accordance with the ethical guidelines released by the International Society for Stem Cell Research (ISSCR). The SOD1^G93A^ transgenic ALS pig model was generated in our lab as described previously(*23*). Four to five-year-old WT and ALS pigs were used in the experiment. The TIA1^P362L^ point mutation ALS rabbit model was generated in our lab as described previously(*56*). WT Rabbits were purchased from HuNan TaiPing Biotech. Six-week-olds NCG (NOD/ShiltJGpt-*Prkdc*^em26Cd52^*Il2rg*^em26Cd22^/Gpt) mice were bought from Gempharmatech and housed in controlled conditions, humidity and a light-dark cycle in a specific pathogen-free facility following the ‘Guide for the Care and Use of Laboratory Animals (2011)’.

### Cell lines

Human WT iPSCs (UH10) were previously generated from human umbilical cord blood and were a gift from Professor Guangjin Pan’s laboratory in GIBH. UH10, NILB-iPSCs, KO-NILB-iPSCs and HIP-NILB-iPSCs were cultured on Matrigel (Corning, 354277) coated plates with mTeSR1 medium (Stem Cell Technologies, 85850) containing 10 nM Y-27632 (Sigma, Y0503). Cells were cultured at 37°C with 5% CO_2_ and passaged using Accutase (Stem Cell Technologies, 07922) when reached 90% confluency. Primary NK cells were isolated from PBMC and cultured with NK cell induction regent kit following manufacturer’s instruction (BaSO Biotechnology Co., Ltd, T2020N2). Human microglial (HMC3) cells (BNCC342264) were bought from BNCC (BeNa Cultute Collection), and cultured in MEMα containing 10% FBS. THP-1 (ATCC) and cultured in RPMI 1640 medium (Gibco, 61870036) containing 10% FBS. Human PBMCs were bought from LDEBIO and maintained in 10% FBS RPMI 1640 medium (Gibco, 61870036).

### Generation of hypoimmunogenic HIP-NILB-iPSCs

UH10 iPSCs that contain doxycycline (Dox)-inducible MNs-specific transcription factors (*NGN2^+^, ISL1^+^, LHX3^+^*), Bcl-XL, firefly luciferase and GFP were constructed with PiggyBac plasmids to generated NILB-iPSCs in our lab as described previously(*25*). To generate hypoimmunogenic iPSCs, NILB-iPSCs underwent two rounds of genetic-editing. Firstly, we knock out the *B2M*, C*IITA*, *IL6ST,* and *TNFRSF1A* genes simultaneously using CRISPR-Cas9 technology. In detail, gRNAs targeting human *B2M*, *CIITA*, *IL6ST,* and *TNFRSF1A* coding regions were designed (https://www.benchling.com/crispr, **Table S1**), annealed and linked in tandem while U6 promoters and gRNA scaffolds were added to ensure the equal disruption efficiency in each gene (**fig. S1A**). The sequences were then cloned into pSpCas9(BB)-2A-Puro (PX459) (Addgene, 108292) plasmids. Thus, each gRNA was functioning in a single cassette within this all-in-one plasmid. NILB-iPSCs were transfected with the all-in-one vectors using the electroporation condition program B-016 using the 2B-Nucleofector (Lonza) system. The transfected iPSCs were cultured in mTeSR medium containing 10 nM Y-27632 (Sigma, Y0503) for 24 h, then changed medium and further screened by treating with puromycin (0.5 μg/mL, Sigma, 540411) for a week. Single clones were picked, amplificated and used for Sanger sequencing and identification to generate a *B2M*^-/-^*CIITA*^-/-^*IL6ST*^-/-^*TNFRSF1A*^-/-^ iPSC lines (KO-NILB-iPSCs). Secondly, we overexpressed CD47, CD24, PD-L1 and CTLA-4-Ig in these cells using PB-CAG-BGHpA (Addgene, 92161) piggyBac system. The coding sequences of CD47 and CD24 linked with P2A were synthesized and inserted into the ORF under the control of the CAG promotor. In the opposite direction, *CTLA4*-*IgG1 Fc*-P2A-*PD-L1*-T2A-Puro sequences were cloned driving by human ubiquitin C (UbC) promoter. The resulting plasmids express CD47, CD24, PD-L1, a fusion protein CTLA-4-Ig and puromycin respectively. 5 μg above constructed transposable plasmids and 5 μg piggyBac transposase plasmids PB200PA-1 were co-transfected into *B2M*^-/-^*CIITA*^-/-^*IL6ST*^-/-^*TNFRSF1A*^-/-^ hiPSCs using Lonza Nucleofector system. Selection of the stable clones used 0.5 μg/mL puromycin for 7 days until the appearance of drug-resistant cells, and the survival cells were expanded and validated.

### MNs differentiation

For *in vitro* differentiation, HIP-NILB-iPSCs, KO-NILB-iPSCs or NILB-iPSCs were seeded into plates pre-coated with matrix gel for 24 hours to reach 80% confluence. The mTeSR medium containing 2 μg/mL Dox (Sigma, 324385) was freshed daily for 4 days, followed by the analysis of these differentiated cells. For the transplantation experiments *in vivo*, HIP-NILB-iPSCs or NILB-iPSCs were seeded into plates pre-coated with matrix gel for 24 hours to reach 80% confluence. The fresh mTeSR medium containing 2 μg/mL Dox (Sigma, 324385) was used to culture these cells for 12 hours. Then, the cells were transplanted into animal spinal cords, followed by Dox treatment for 3 days *in vivo*. In detail, rabbits and pigs were intragastric or orally administrated, at a dose of 25 mg/kg/d and 50 mg/kg/d, respectively.

### G-band karyotyping

HIP-NILB-iPSCs were cultured in 6-cm plates until in a logarithmic growth phase, followed by 0.5 μg/ml colcemid treatment (Millipore, 234109-M) for 1 hour. The cells were then trypsinized, centrifuged at 300 *g* for 5 mins, and resuspended with 0.075 M KCl solution, followed by incubating for 30 mins at 37°C. To fix the cells, a fixative solution composed of one part of acetic acid and three parts of methanol was added and incubated for 10 mins at 37°C. After centrifugation, cell pellets were resuspended with 10 ml ice-cold fixative solution, dropped onto a clean glass slide and incubated at 75°C for 3 hours to allow the cell to spread out. The slides were then treated with 0.25% trypsin and stained with Giemsa. The results were acquired and analyzed with an Olympus microscope.

### Immunofluorescence assay

For iPSCs and MNs immunofluorescence staining, cells were fixed in 4% paraformaldehyde (PFA) for 1 hour at 4°C, followed by blocking and permeabilizing in 5% BSA with 0.2% TritonX-100 for 30 mins. For brain and spinal cords, tissues of different sizes were fixed in 4% PFA at 4°C for 36 to 72 hours, and gradiently dehydrated in 10%, 20%, and 30% sucrose solution for 48 hours until the tissues sank. The tissue blocks were then embedded in an optimal cutting temperature (OCT) compound (Sakura, 4583), quickly frozen with liquid nitrogen, and cut into a thickness of 30 μm slices using a cryomacrotome (Leica CM 3050S). Tissue slices were blocked and permeabilized in 5% BSA (Sigma, V900933) with 0.5% TritonX-100 solution for 2 hours at room temperature. After that, specific primary antibodies targeted different proteins were diluted at a certain concentration and incubated onto the slices at 4°C overnight. After washing with 0.1% PBS-T three times, the tissue sections were incubated with the indicated fluorescence-conjugated secondary antibodies for 1 hour at room temperature. DAPI (Beyotime, C1006) at 2 μg/ml was used to label cell nuclei and then the slides were washed for mounting. Images were acquired on LSM 710 confocal microscope (Zeiss) and IXplore SpinSR (Olympus) at a certain magnification. ImageJ software (http://rsb.info.nih.gov/ij/) was used to analyze the images. The primary antibodies were used as follows: chicken anti-GFP polyclonal antibody (ThermoFisher, A10262, 1:300), mouse anti-GFP monoclonal antibody (Proteintech, 66002-1-Ig, 1:300), rabbit anti-GFP polyclonal antibody (Proteintech, 50430-2-AP, 1:300), mouse anti-beta III Tubulin (TUJ1) monoclonal antibody (Abcam, ab78078, 1:500), mouse anti-MAP2 monoclonal antibody (Sigma-Aldrich, M1406, 1:300), Rabbit anti-Choline Acetyltransferase (ChAT) monoclonal antibody (Sigma-Aldrich, ab181023, 1:300), mouse anti-GFAP monoclonal antibody (Santa Cruz, sc-33673, 1:1000), mouse anti-Nuclei monoclonal antibody (Millipore, MAB4383, 1:200), mouse STEM121 antibody (Takara, Y40410, 1:400), mouse anti-Neurofilament 200 monoclonal antibody (Sigma-Aldrich, N5389, 1:400), α-Bungarotoxin, Alexa Fluor® 647 Conjugate (ThermoFisher, B35450, 1:1000), rabbit anti-CHRNA7 polyclonal antibody (Proteintech, 21379-1-AP, 1:200), rabbit anti-Synaptophysin (Syn) monoclonal antibody (Abcam, ab52636, 1:200), rabbit anti-Vimentin polyclonal antibody (Proteintech, 10366-1-AP, 1:200), rabbit anti-Myelin basic protein (MBP) polyclonal antibody (Proteintech, 10458-1-AP, 1:200), rabbit anti-NLRP3 polyclonal antibody (Proteintech, 19771-1-AP, 1:200), rabbit anti-IBA1 polyclonal antibody (Proteintech, 10904-1-AP, 1:500), rabbit anti-TDP43 polyclonal antibody (Proteintech, 10782-2-AP, 1:200). The following fluorescence labeled secondary antibodies were used: CoraLite488 conjugated Affinipure Goat Anti-Rabbit IgG(H+L) (Proteintech, SA00013-2, 1:1000), CoraLite488 conjugated Affinipure Goat anti-mouse IgG(H+L) (Proteintech, SA00013-1, 1:1000), Goat anti-Chicken IgY H&L Alexa Fluor® 488 Conjugate (Abcam, ab150173, 1:1000), Anti-mouse IgG (H+L), F(ab’)2 Fragment (Alexa Fluor® 555 Conjugate) (CST, 4409S, 1:1000), Anti-rabbit IgG (H+L), F(ab’)2 Fragment (Alexa Fluor® 555 Conjugate) (CST, 4413S, 1:1000), Anti-mouse IgG (H+L), F(ab’)2 Fragment (Alexa Fluor® 647 Conjugate) (CST, 4410S, 1:1000), Anti-rabbit IgG (H+L), F(ab’)2 Fragment (Alexa Fluor® 647 Conjugate) (CST, 4414S, 1:1000).

### *Alu* gene detection in rabbits and pigs

The genomic DNA of tissues and HIP-NILB-iPSCs was extracted with FastPure Cell/Tissue DNA Isolation Kit (Vazyme, DC102-01) and *Alu* PCR primers were synthesized as follows: forward 5’-CTGTATACTCAGCTACTAGGGT-3’, reverse 5’-CTCAGGGGTTATCTAAAGTGGC-3’. 1 μg DNA template was amplificated with Taq polymerase (Vazyme, P222) for 35 cycles of 95°C denaturation for 15 s, 60°C annealing for 15 s, and 72°C extension for 8 s. The samples in lumbar spinal cord of grafted rabbits or pigs were used as the experimental positive control, HIP-NILB-iPSCs DNA used as technical positive control, and water added as template was a negative control. *GAPDH* was chosen as an internal reference, and the primers used as follows: human *GAPDH* forward 5’-GTCAAGCTCATTTCCTGGTATGAC-3’, reverse 5’-CAGTGTGGTGGGGGACTGAG-3’; rabbit *GAPDH* forward 5’-GCCTGGAGAAAGCTGCTAAGT-3’, reverse 5’-GGCTCTTACTCCTTGGAGGC-3’; pig *GAPDH* forward 5’-GTCGGAGTGAACGGATTTGGC-3’, reverse 5’-CTTGCCGTGGGTGGAATCAT-3’.

### Real-time PCR and Western blotting

Spinal cord tissues were homogenized with RNAiso Plus (Takara, 9108) in a dounce tissue grinder at 4°C. The total RNA was extracted following standard processes, measured and quality checked using NanoDrop 1000 (Thermofisher). 1 μg total RNA was reverse transcribed to cDNA after eliminating genome DNA using HiScript III RT SuperMix (Vazyme, R323-01). Quantitative PCR was then performed with AceQ Universal SYBR qPCR Master Mix (Vazyme, Q511-02) on CFX96 Real-Time System (Bio-Rad) using primers shown in **Table S1**. The experiments were conducted in triplicate, and the mRNA levels of each target were normalized to *GAPDH* transcripts and calculated by 2^-ΔΔCT^. For western blot, cells were collected and lysed in RIPA buffer (Beyotime, P0013B) with 1mM PMSF (Beyotime, ST2573) and protease inhibitor cocktail (Roche Biochemicals, 04693116001). Spinal cord tissues from rabbits and pigs were homogenized using a high-speed tissue homogenizer followed by centrifuging at 13,000 *rpm* for 10 mins at 4°C, and the supernatants were collected. The total protein concentrations from cell and tissue samples were measured by the BCA protein assay kit (Pierce, A53225) and used for western blot. For soluble and insoluble TDP-43 fractionation, the soluble supernatants and insoluble pellets were separately collected by centrifuging at 13,000 *rpm* for 10 mins at 4°C after 50 μL total protein was reserved. The pellets were subsequently washed with RIPA buffer for three times and solubilized with 8 M urea following centrifugation at 13,000 *rpm* for 10 mins at 4°C. The soluble and insoluble fractions were then used for western blot analysis with TDP-43 antibody (proteintech, 12892-1-AP, 1:1000).

### Bulk RNA-seq

Total RNA was extracted by RNAios Plus (Takara, 9108) and sent for library construction and sequencing on an Illumina HiSeq 3000 sequencer (Genergy Bio-technology). The raw data were tested for quality control by FastQC. Quality control was performed by removing adapter sequences and reads with low complexity or of low quality by using TrimGalore. Clean RNA-seq reads were aligned to GRCh38 (hg38) human reference genome using STAR with default parameters and the number of reads assigned to each gene were quantified using Featurecounts. DEseq2 package was used for variance stabilizing transform (vst) in count matrix, data normalization and differential expression analysis. During the downstream analysis, correlation analysis and Gene Set Enrichment Analysis (GSEA) were performed, ggplot2, pheatmap, ClusterGVis and GseaVis were used for customized visualizations.

### Flow cytometry

Cells were collected and dissociated into single-cell suspension. For surface markers staining, single cell suspensions were directly incubated with fluorescence-labeled diluted antibodies or isotype-matched control antibodies at 4°C for 20 mins in the dark. For intracellular markers, cells were fixed with 4% PFA and permeabilized with 1× Intracellular Staining Permeabilization Wash Buffer Fixation Buffer (BioLegend, 421002), followed by incubation in respective fluorochrome-conjugated antibodies solutions at 4°C for 20-30 mins in the dark. After washing with Cell Staining Buffer (BioLegend, 420201) three times, the samples were loaded to LSR Fortessa SORP (BD), and the data were further analyzed using FlowJo V.10 software. The antibodies used were listed as follows: PE anti-human HLA-A,B,C (BioLegend, 311405, 1:20), PE anti-human HLA-DR,DP,DQ (BioLegend, 361716, 1:20), PE Mouse IgG2a, κ Isotype Ctrl (FC) antibody (BioLegend, 400213, 1:20), PE anti-human CD3 (BD Biosciences, 347347,1:50), FITC anti-human CD107a (BD Biosciences,555800,1:100), FITC Mouse IgG1, κ Isotype Control RUO (BD Biosciences, 555748,1:100), APC anti-human CD56 (BioLegend, 362504, 1:20), PE-Cy5 human CD56 (BD Biosciences, 555517, 1:10), APC Mouse IgG1, κ Isotype Ctrl (FC) antibody (BioLegend, 400122, 1:20), PE anti-mouse CD68 (BioLegend, 137014, 1:100), PE Rat IgG2a, κ Isotype Ctrl (BioLegend, 400508, 1:100), CoraLite® Plus 647-conjugated IBA1 monoclonal antibody (Proteintech, CL647-66827,1:200), CoraLite® Plus 647 Mouse IgG2a Isotype Control (Proteintech, CL647-65244,1:200).

### CFSE-labelled T cell proliferation assay

PBMCs were cultured in RPMI-1640 medium supplemented with 10% FBS for 12 hours and then were labeled with 1 μM CFSE (BestBio, BB-4211) at 37°C in the dark for 10 mins, followed by washed 2 times with FBS-free RPMI-1640 medium. WT iPSCs, Dox-induced HIP-NILB-iPSCs or HIP-NILB-iPSCs were pretreated with IFN-γ (Sigma, SRP3058), followed by incubating with allogenic CFSE-diluted PBMC at 1:1 for 5 days. Suspended PBMCs were collected for flow cytometry. In general, we first gated CD3^+^ T cells using PE-conjugated anti-CD3 antibodies (BD, 347347) and then analyzed the percentage of CD3^+^CFSE^-^ proliferative T cells. PBMCs cultured alone or activated by 2 μg/mL phytohemagglutinin P (PHA-P) (Sigma, L8754) were used as negative or positive controls, respectively.

### Phagocytosis Assay

The human monocytic THP-1 cells were cultured in RMPI-1640 containing 10% FBS and were incubated with 100 ng/mL PMA for 2 days to differentiate into macrophages. After that, THP-1 macrophages were exposed to the RPMI-1640 culture growth medium with LPS (100 ng/mL) and IFN-γ (20 ng/mL) for 48 h reversal to M1 macrophages. The M1 macrophages were stained with Cell-Tracker Deep Red (Invitrogen, C34565). We have generated iPSCs carrying green fluorescent protein (GFP). The iPSCs (2×10^5^/well) were incubated with labeled human macrophages (2×10^5^/well) in 96-well plates for 2 hours at 37°C. Then, the cell mixtures were harvested, and the proportion of Deep Red/GFP double positive cells was analyzed by flow cytometry to evaluate the phagocytic function of the human macrophages.

### Protein array assay

HIP-NILB-iPSCs and NILB-iPSCs cultured in 6-well plates were treated with 100 ng/ml lipopolysaccharide (LPS, Sigma, L5418) for 6 hours to stimulate inflammatory factors secretion. Fresh mediums were changed and medium supernatants were collected after 24 hours. The supernatant was analyzed by a membrane-based human inflammation antibody microarray to measure 20 inflammatory cytokines following the manufacturer’s instructions (RayBiotech, AAH-INF-1-2). In general, the microarray was blocked at room temperature for 1 hour and incubated with 700 µl supernatant overnight at 4°C. After washing, the microarray was incubated with 1 ml diluted biotin-labelled antibody for 2 hours at room temperature, followed by incubation with HRP-conjugated streptavidin for 1 hour in the dark. The detective substrates were applied for coloration. The membranes were scanned and the images were acquired with ImageQuant LA4000 Scanner (GE Healthcare Corporate).

### ELISA for IFN-γ secretion

One million PBMCs or primary NK cells (effector cells) were incubated with WT iPSCs, Dox-induced HIP-NILB-iPSCs or HIP-NILB-iPSCs (target cells) at a ratio of 1:1 for 5 days, respectively. The supernatants were collected and a human IFN-γ ELISA Kit (MultiSciences, EK180) was used to measure the IFN-γ secretion.

### Lactate dehydrogenase (LDH) assay for cell cytotoxicity

Using coculture experiments, we measured LDH released in the medium using an LDH cytotoxicity assay kit (Beyotime, C0016) following standard instructions. In brief, 1 × 10^6^ target cells were mixed with effector cells at a ratio of 1:1 for 5 days. Medium from only effector cells was set as the spontaneous LDH activity group and the effector cells treated with lysis buffer as the maximum LDH activity group. Absorbances at 490 nm and 680 nm (background signal from the instrument) were measured in a 96-well luminometer plate. The percent cytotoxicity was calculated after subtracting the 680 nm absorbance value from the 490 nm absorbance and the formula as follows: (LDH activity from coculture group minus spontaneous LDH activity)/(maximum LDH activity minus spontaneous LDH activity) × 100.

### Survival and tumorigenicity of HIP-NILB-iPSCs in huPBL-NCG

To construct huPBL-NCG model, we intravenously injected 4 × 10^6^ human PBMCs into NCG mice. After 7 days, one million NILB-iPSCs, HIP-NILB-iPSCs or 12 hour-Dox-treated HIP-NILB-iPSCs in 100 μL were subcutaneously transplanted into huPBL-NCG mice at inguinal region, respectively. For 12 hour-Dox-treated HIP-NILB-iPSCs group, HIP-NILB-iPSCs were induced with Dox (2 μg/mL) for 12 hours prior to transplantation, and Dox (25 mg/kg/d) treatment for 3 days was applied to induce their differentiation in *vivo* further. The luciferase signals in each group were detected at 4 hours and 1, 3, 5, 7, 14, 21, 28, 40 days via intraperitoneally injection with 150 mg/kg luciferin substrate (Promega, E1605) in PBS and the average radiance was recorded with the Multi-Mode In Vivo Imaging System (BLTlux, AniView600). After 40 days, all mice were euthanized to assess teratoma formation.

### Creatine kinase (CK) assay and Glutamate assay

Serum from rabbits and pigs was collected and used to detect CK levels (A032-1-1, Nanjing Jiancheng Bioengineering Institute) and glutamate levels (A074-1-1, Nanjing Jiancheng Bioengineering Institute) following the manufacturer’s instructions.

### Pluripotency analysis *in vitro* and *in vivo*

To investigate the pluripotency of HIP-NILB-iPSCs and WT iPSCs *in vitro*, they were cultured on Matrigel-coated dishes until colony formation. NANOG (R&D, AF1997, 1:200), OCT4 (Santa Cruz, sc-5279, 1:200), SOX2 (R&D, MAB2018, 1:250), and SSEA4 (ThermoFisher, 46-8843-41, 1:300) were used for the immunofluorescence assay. For *in vivo* experiment, five million WT iPSCs or HIP-NILB-iPSCs were subcutaneously transplanted into inguinal region of NCG mice. After 21 days, teratoma formations were fixed with 4% PFA, followed by gradient dehydration, embedding, days, teratomas were collected, fixed, embedded and sectioned for H&E staining.

### H&E, Masson, and Nissl staining

The gastrocnemius muscle samples of pigs and rabbits were collected and fixed in 4% PFA for 48 hours followed by dehydrating and clearing. Once the tissue blocks were embedded with liquid paraffin, 4 μm slices were cut and used for H&E staining and Masson staining. For Masson staining, Weigert’s iron hematoxylin was used to highlight the nuclei in dark blue. Instead, the collagen fibers were stained blue with aniline blue solution for 5 mins, and Ponceau fuchsin stained the cytoplasm and muscle fibers red or pink. The mounted slides were finally scanned using the Tissue FAXS viewer (Tissue Gnostics). Nissl staining was used to visualize the distribution and morphology of neurons in spinal cord tissue. Spinal cord slices were collected serially at 30 μm intervals, and dehydrated in an ethanol gradient, followed by defatting in xylene. After rehydration, slices were stained in 1% cresyl violet for 10 mins. Then, slices were dehydrated in absolute ethanol and cleared in xylene. Finally, all slices were mounted in neutral balsam and examined under a bright-field microscope. The neuron number of the ventral horn was calculated by counting the cresyl violet-stained cell body in each slice.

### Gait analysis and muscular strength measurement in rabbit hindlimbs

Gait analysis was performed to assess the motor ability of WT, ALS and HIP-NILB-iPSCs-treated rabbits. In detail, customized runway was laid out with a white paper measuring 3.5 m in length and 0.4 m in width. Prior to the test, rabbits were trained to run down the runway without interruption. The rabbit paws were daubed over non-toxic pigment (forepaws in blue, hindpaws in red) for footprints recording. The stride length (distance between two successive footprints of the same hindpaw) and sway length (distance between the right and left hindpaw) were then measured and analyzed. Rabbit hindlimb muscular strength was measured with a digital force gauge (Hemuele, ZMF-100, range 0.01-100 kg) fitted with a metal hook. In brief, rabbits were held and the hind legs were kept in a straight and relaxed position to ensure accurate measurements and to minimize stress on the rabbits. The hind legs of rabbits were then connected to the metal hook of the force gauge that horizontally fixed on the table. A force stimulator was applied to pinch the skin on the gastrocnemius muscle for once to evoke a flexor reflex and thus exert force on the force gauge. The peak force was then measured and recorded. Each rabbit’s legs are measured 8-10 times, and the tension index was calculated by the ratio of average hind limp strength/body weight.

### Resting EMG recording

Pigs were anesthetized with 2.5% isoflurane in room air via a facemask and the hair on the hindlimb was shaved, and rabbits were held and kept the hind limps relax followed hair shaving. Resting EMG potentials were recorded with two platinum transcutaneous needle-recording electrodes which were positioned into the gastrocnemius muscle with a distance between recording electrodes of ∼1 cm in each muscle. Electrodes were connected to an active multichannel physiologic recorder (MP160, BIOPAC) and a ground electrode was subcutaneously placed in the back. The recorded signals were amplified using the myoelectric amplifier module (EMG100C, BIOPAC) following digitized and stored for analysis using the AcqKnowledge 5.0 software (BIOPAC). The recorded signal was sampled at 50 kHz over 5 mins.

### Transplantation of cells into ALS animals

HIP-NILB-iPSCs or NILB-iPSCs were gently dissociated into single cells with accutase, resuspended in mTeSR medium. After the intravenous administration of propofol (1 mg/kg, Petsun Therapeutics), pigs were anesthetized with 3.5% inhalable isoflurane and placed in a surgery bed. The dura mater was punctured using a bent 30-gauge needle, and a total of 10,000,000 HIP-NILB-iPSCs in 100 μL was injected into the lumbar spinal cord at three points (L2, L3, L4) in a row at a rate of 1 μL per 5 s. Dox (50 mg/kg) was treated for 3 days to induce MNs differentiation through oral administration. For rabbits, after being anesthetized with 3% isoflurane, the surgical area was shaved and cleaned, and a skin incision was made at the L2-L4 vertebral level. The dura overlying the L3 spinal segments punctured and the blunt 30-gauge injection needle containing the HIP-NILB-iPSCs was then inserted into the lumbar spinal cord using a fine XYZ manipulator (SMM 100B, Narishige). One point on the third lumbar vertebra (L3) was chosen for injection. A total of 4,000,000 HIP-NILB-iPSCs or NILB-iPSCs in 20 μL or 20 μL mTeSR medium were injected into each rabbit. Dox (25 mg/kg) treatment via gavage administration was applied for 3 days to induce MNs differentiation.

### Retrograde tracing of MNs through intramuscular injection of the AAV2-retro viruses

To examine whether HIP-NILB-iPSCs-derived MNs project to motor endplates, rAAV2-retro-hSyn-mCherry-WPRE-hGH polyA viruses (Brainvta, PT-0100) were injected into the gastrocnemius muscle of HIP-NILB-iPSCs-treated rabbits at 2 months post-grafting. In general, the rabbits were restrained following clearing the hair above the gastrocnemius muscles and disinfecting the skin with iodine. The viruses were diluted to a final titer of 2.9 × 10^12^ genome copies/mL in 200 μL with physiological saline solution prior to injection. A total of 2.9 × 10^11^ viral particles were injected into gastrocnemius muscle of each leg for five sites and each site for 20 μL at a rate of 2 μL/min. The injection needle being held in place for 30 s, followed by being slowly withdrawn from the gastrocnemius muscle. After 30 days, the rabbits were sacrificed for further analysis.

### Retrograde trans-monosynaptic tracing with rabies virus

The HIP-NILB-iPSCs were cultured to logarithmic growth phase and then infected with 100 µL LV-EF1a:G virus (5 × 10^8^ Tu/ml, BrainVTA) in the presence of infective enhancer A+B (BrainVTA) for 3 days. For rabies tracing, 4,000,000 HIP-NILB-iPSCs in 10 μL that expresses EF1a:G were incubated with Dox for 12 hours, followed by being transplanted into rabbit lumbar spinal cord together with 1 μL RV-dG-tdTomato (2 × 10^8^ IFU/ml, BrainVTA) and treated with Dox (25 mg/kg) via gavage administration for 3 days. After 11 days, the rabbits were sacrificed for further analysis.

### Statistical analysis

All data were expressed as mean ± SEM and analyzed using GraphPad Prism 9. The sample number (n) indicates the number of animals or cellular experiment repeats, as specified in the figure legends. For data with two groups, the mean values from individual groups were compared using either an independent sample unpaired, two-tailed, Student’s *t-test* or Welch’s *t*-*test*. Statistical comparisons between three groups were made using one-way analysis of variance (ANOVA) followed by Tukey’s or Dunnett’s T3 multiple comparisons. Two-way ANOVA with Tukey’s multiple comparisons were made in rabbits over-time tension index analysis. Exact *p* values are described in each figure and between the indicated groups.

## List of Supplementary Materials

Fig. S1 to Fig. S14

Table S1

## Acknowledgments

Schematic diagrams were created by BioRender.

## Funding

National Key Research and Development Program of China (2022YFA1105403)

National Key Research and Development Program of China (2023YFC3404305)

National Key Research and Development Program of China (2022YFF0710601)

The National Natural Science Foundation of China (32400806)

The National Natural Science Foundation of China (32470891)

The Science and Technology Service Network Initiative of the Chinese Academy of Sciences-Huangpu Program (STS-HP-202204)

Guangzhou Key Research Foundation (2023B03J0011)

Hainan Provincial Key Research and Development Program (ZDYF2021SHFZ230)

Major Science and Technology Project of Hainan Province (ZDKJ2021030)

2020 Research Program of Sanya Yazhou Bay Science and Technology City (202002011)

Guangzhou Basic and Applied Basic Research Foundation (2023A04J0730)

Guangzhou Basic and Applied Basic Research Foundation (2023A04J0108)

Science and Technology Planning Project of Guangdong Province (2020B1212060052)

Science and Technology Planning Project of Guangdong Province (2021B1212040016)

Research Unit of Generation of Large Animal Disease Models, Chinese Academy of Medical Sciences (2019-I2M-5-025)

## Author contributions

N.Z., Y.Y., M.C., Q.Z., W.X., and Z.D. performed experiments and analyzed data. L.Q., Q.Z., J.H., Y.Z., and Z.Z. collected samples. Z.L., M.C., and K.Z. gave advises for the project. Z.O., Y.S., A.S., M.O., and X.L. helped to revise the manuscript. W.X., Z.D., and L.L. designed and supervised the project. N.Z., Z.D., and L.L. wrote the manuscript.

## Data and materials availability

All data are available and can be requested by contacting the corresponding author. Raw RNA-seq data are accessible through the CNGB Sequence Archive (CNSA) of China National GeneBank DataBase (CNGBdb) with the accession number CNP0004683 (http://db.cngb.org/cnsa/project/CNP0004683_fdfb1785/reviewlink/). The data of this study were analyzed with standard software that is already available and does not report original code.

## Supplementary Figures

**Fig. S1.**
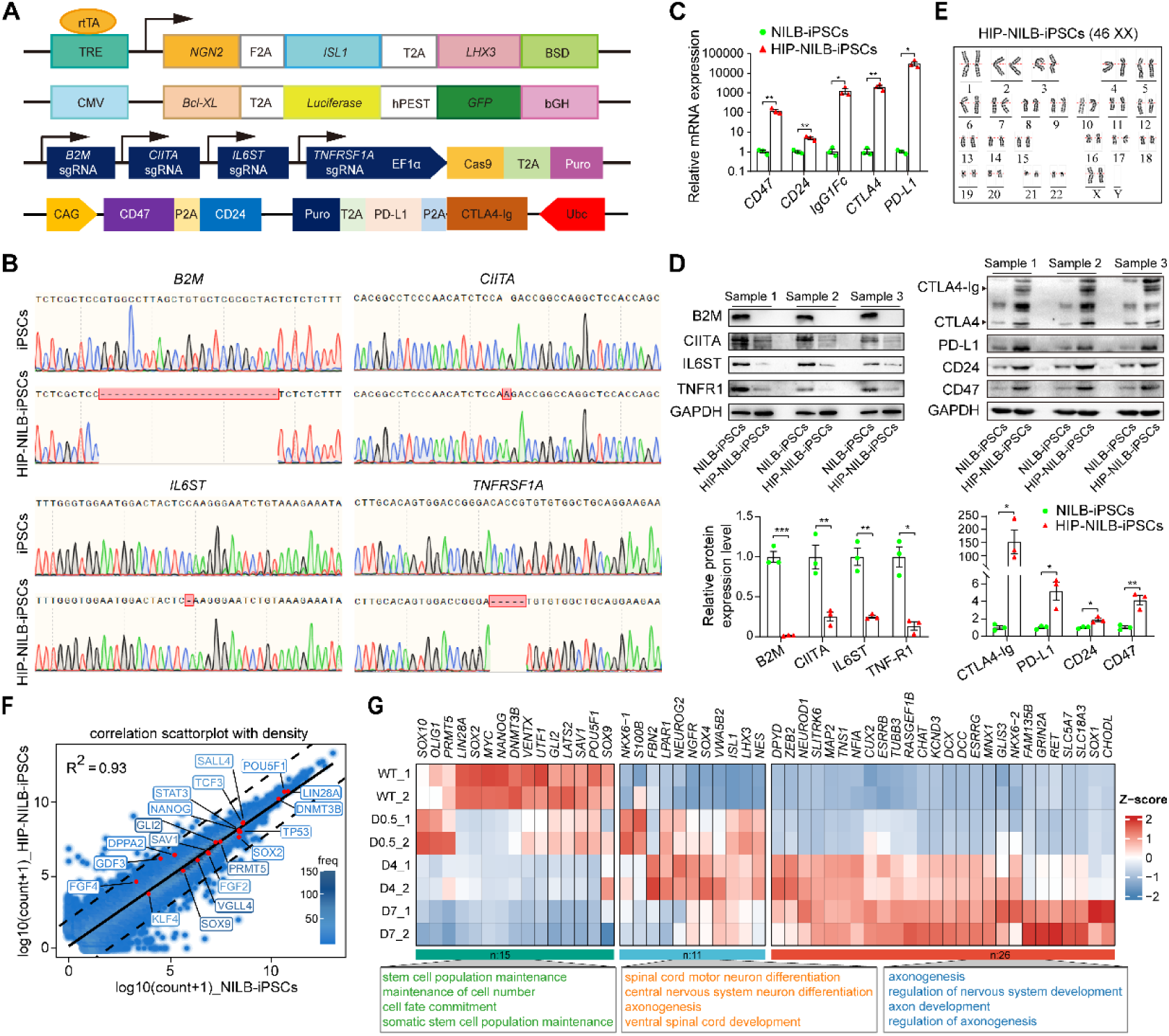
Engineered HIP-NILB-iPSCs via multiplex genetic-editing. **(A)** Schematic illustration of multiplex genetic-editing strategies for engineered HIP-NILB-iPSCs. **(B)** Indel analysis by Sanger sequencing for *B2M*, *CIITA*, *IL6ST* and *TNFRSF1A* in HIP-NILB-iPSCs. **(C)** Real-time PCR analysis of *CD47*, *CD24*, *IgG1-Fc*, *CTLA4* and *PD-L1* in HIP-NILB-iPSCs. **(D)** Western blot analysis and relative density quantification of B2M, CIITA, IL6ST, TNFR1, CTLA4-Ig, PD-L1, CD24 and CD47 in HIP-NILB-iPSCs of three independent samples. CIITA was detected in NILB-iPSCs and HIP-NILB-iPSCs upon IFN-γ and TNF-α treatment for 48 h. **(E)** Karyotyping of HIP-NILB-iPSCs. **(F)** Scatter plot comparison of transcriptomes for HIP-NILB-iPSCs and NILB-iPSCs. The raw counts for each gene were transformed to log10 value, and genes with more than 1 count in each sample were shown. The pluripotent-related genes were labeled in the diagram. Dashed lines depict the 10-fold changes. The R^2^ was determined by Pearson’s correlation. The NILB-iPSCs were used as a control. **(G)** Heatmap showing expression level of stem cells-related and neuron development-related genes together with the enrichment analysis in WT iPSCs and HIP-NILB-iPSCs upon Dox induced for 0.5-, 4- and 7-days. Data are mean ± SEM; p values were determined using a two-tailed, unpaired Student’s *t-test*; \**p* < 0.05; \*\**p*< 0.01.

**Fig. S2.**
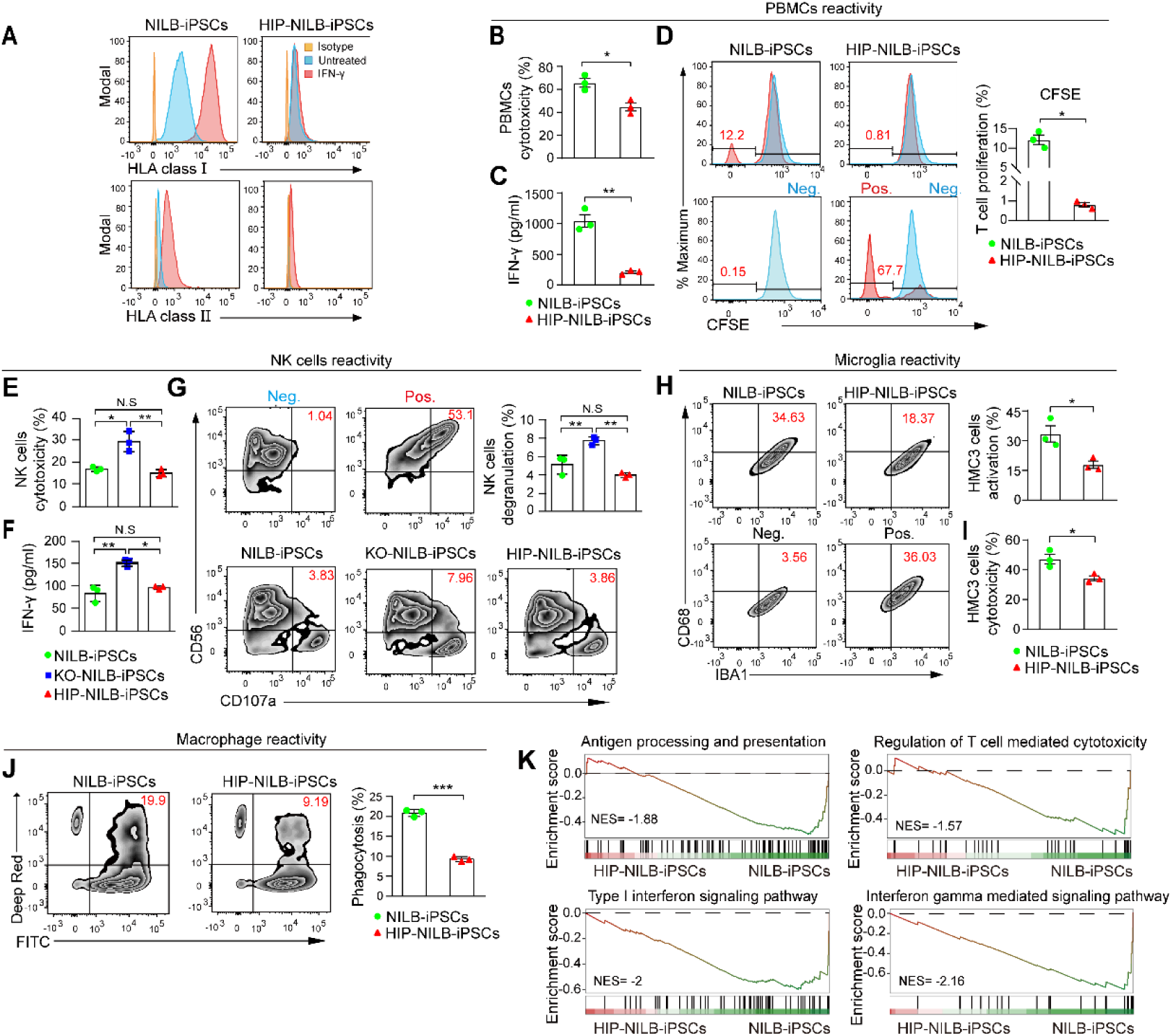
Immune response against HIP-NILB-iPSCs *in vitro.* **(A)** HLA class I and II expression in HIP-NILB-iPSCs with or without IFN-γ treatment. Isotype is a negative control with matched primary antibody. **(B)** PBMCs cytotoxicity against HIP-NILB-iPSCs by measuring LDH release. **(C)** ELISA assay for IFN-γ secretion in PBMCs cocultured with HIP-NILB-iPSCs. **(D)** CFSE analysis of T cells (CD3^+^) proliferation in PBMCs when cocultured with HIP-NILB-iPSCs. PBMCs is negative control (Neg.), PBMCs activated by PHA is positive control (Pos.). **(E)** Primary NK cells cytotoxicity against HIP-NILB-iPSCs by measuring LDH release. **(F)** ELISA assay for IFN-γ secretion in Primary NK cells cocultured with HIP-NILB-iPSCs. **(G)** NK degranulation assay by quantifying CD107a surface expression in CD56^+^ NK cells cocultured with HIP-NILB-iPSCs. NK cells alone as Neg., NK cells treated with PHA as Pos.. **(H)** Activation of HMC3 cells (CD68^+^) in IBA1^+^ when incubated with HIP-NILB-iPSCs. HMC3 alone as Neg., LPS-stimulated HMC3 as Pos.. **(I)** HMC3 cells cytotoxicity against HIP-NILB-iPSCs by measuring LDH release. **(J)** The phagocytic activity of THP-1 derived macrophages against HIP-NILB-iPSCs is characterized by the percentage of double positive of Deep Red and FITC. **(K)** Gene set enrichment analysis (GSEA) shows the enriched signaling pathway in HIP-NILB-iPSCs against NILB-iPSCs. NES represents the normalized enrichment score. All experiments were independently duplicated three times. Data are mean ± SEM, *p* values were determined using a two-tailed, unpaired Student’s *t-test* (b, c, e, f, g, h, i, j) or Welch’s t-test (D), \**p* < 0.05, \*\**p*< 0.01, \*\*\**p* < 0.001, N.S, not significant.

**Fig. S3.**
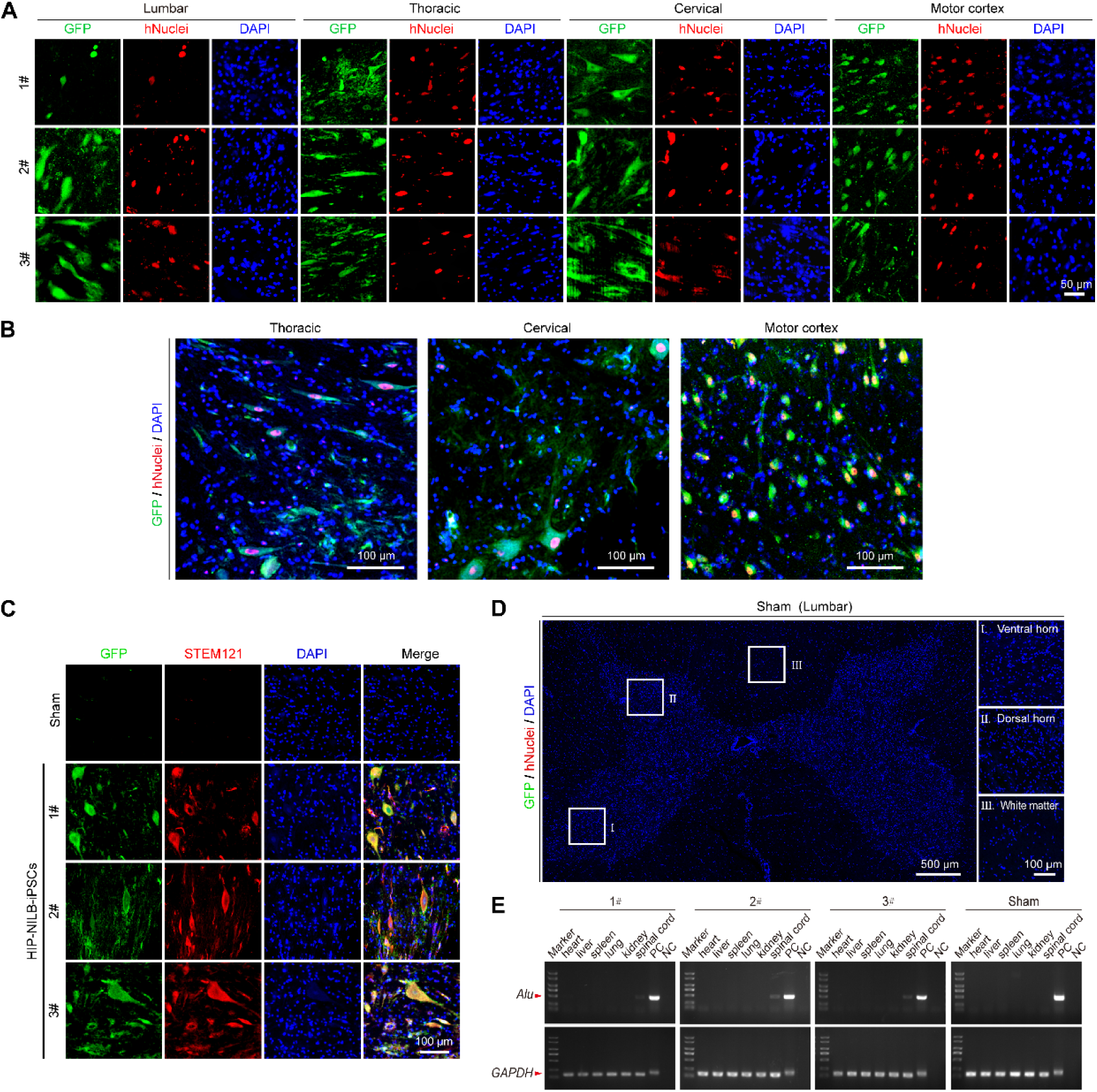
HIP-NILB-iPSCs survived in the CNS of ALS pigs. **(A)** The single channel images corresponding to Figure 2B. Scale bar, 50 μm. **(B)** GFP^+^hNuclei^+^ HIP-NILB-iPSCs migrated and survived in the thoracic, cervical spinal cords and motor cortex of ALS pig (2#) with a lower magnification (corresponding to Figure 2B). Scale bar, 100 μm. **(C)** Representative confocal images of GFP and STEM121 staining in HIP-NILB-iPSCs treated pigs. Scale bar, 100 μm. **(D)** Representative images of GFP and hNuclei staining in lumbar spinal cord sections of sham ALS pig. Scale bars, 500 μm or 10 μm. **(E)** *Alu* PCR analysis of various organs and spinal cords in treated pigs and the sham control.

**Fig. S4.**
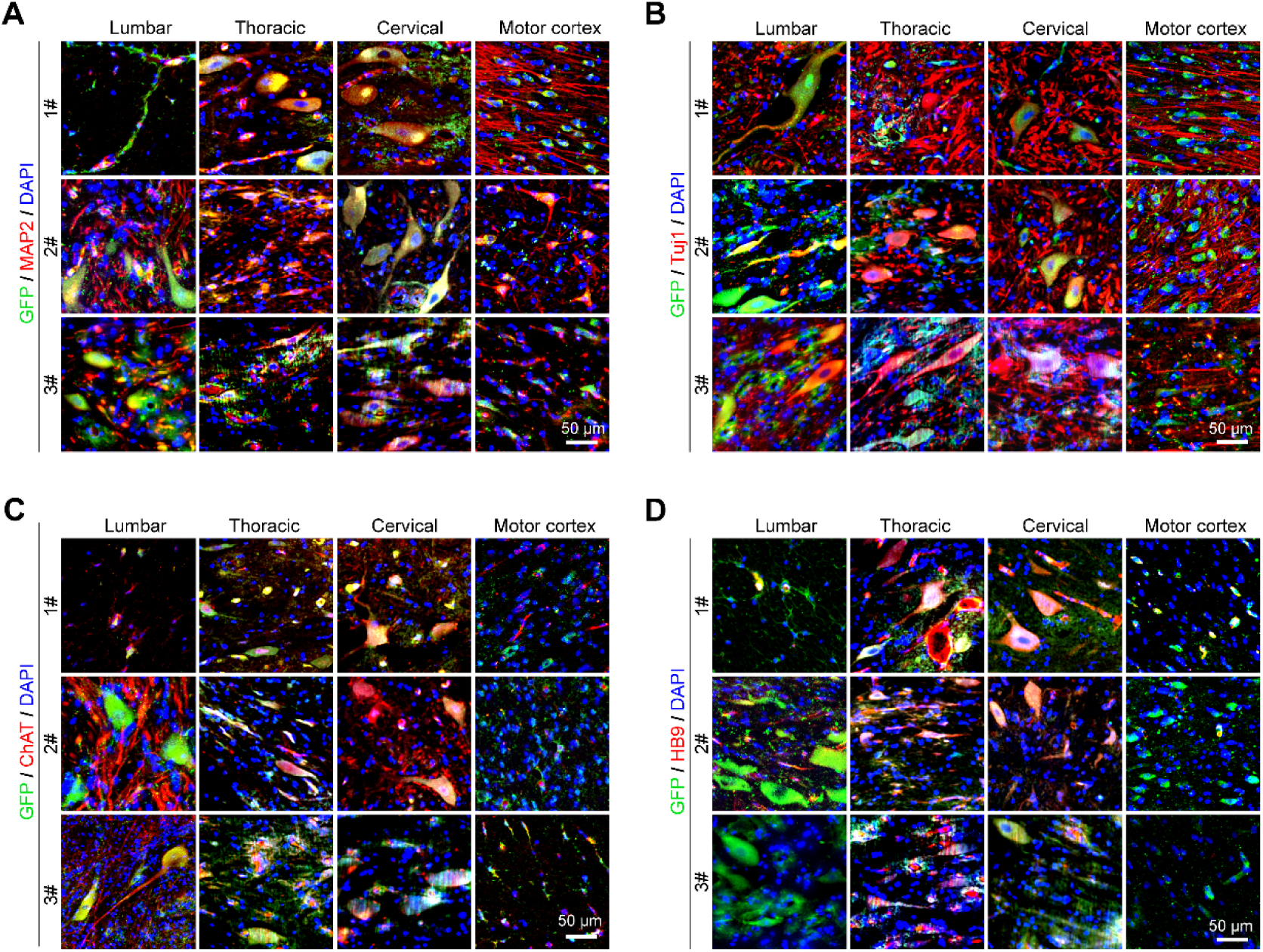
Widespread distribution of HIP-NILB-iPSCs-MNs. (A-D) Representative immunofluorescence images showed the expression of neuron markers (A) MAP2 and (B) TUJ1, and MNs markers (C) ChAT and (D) HB9 of HIP-NILB-iPSCs-derived MNs in lumbar, thoracic, cervical spinal cords, and motor cortex in three HIP-NILB-iPSCs treated ALS pigs. Scale bar, 50 μm.

**Fig. S5.**
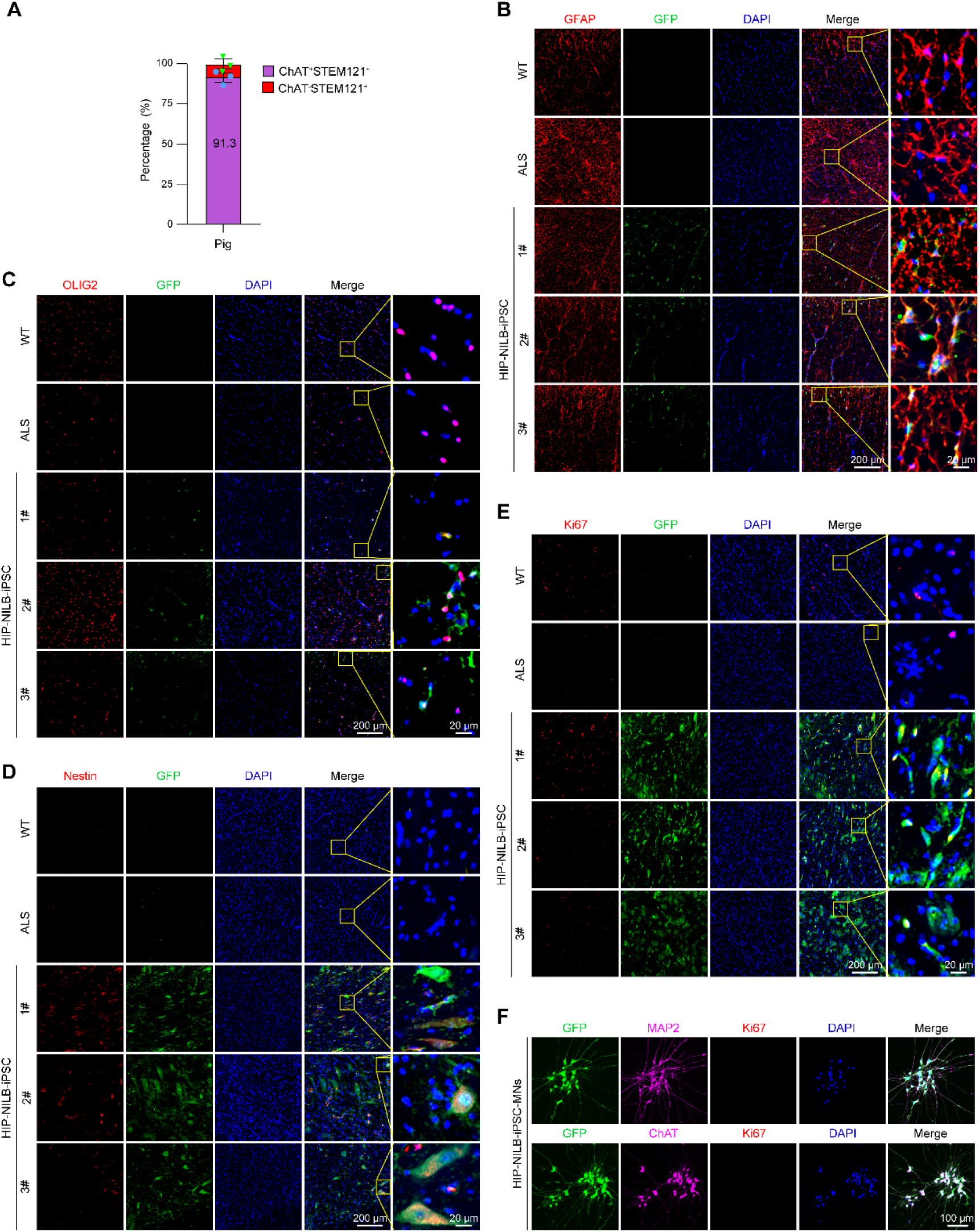
HIP-NILB-iPSCs mainly differentiated into MNs in ALS pigs. **(A)** Quantitative percentage of STEM121 positive HIP-NILB-iPSCs-derived MNs in HIP-NILB-iPSCs treated ALS pigs (n=3). **(B-D)** Representative images showing very few HIP-NILB-iPSCs differentiated into (B) astrocytes, (C) oligodendrocyte and (D) neural stem cells in the lumbar spinal cords of three HIP-NILB-iPSCs treated ALS pigs. Scale bars, 200 μm or 20 μm. **(E)** Representative images of Ki67 staining in the lumbar spinal cords of HIP-NILB-iPSCs grafted ALS pigs. Scale bars, 200 μm or 20 μm. **(F)** Representative images show no Ki67^+^ cells in HIP-NILB-iPSCs MNs *in vitro*. HIP-NILB-iPSCs were induced upon Dox for 5 days. Scale bar, 100 μm.

**Fig. S6.**
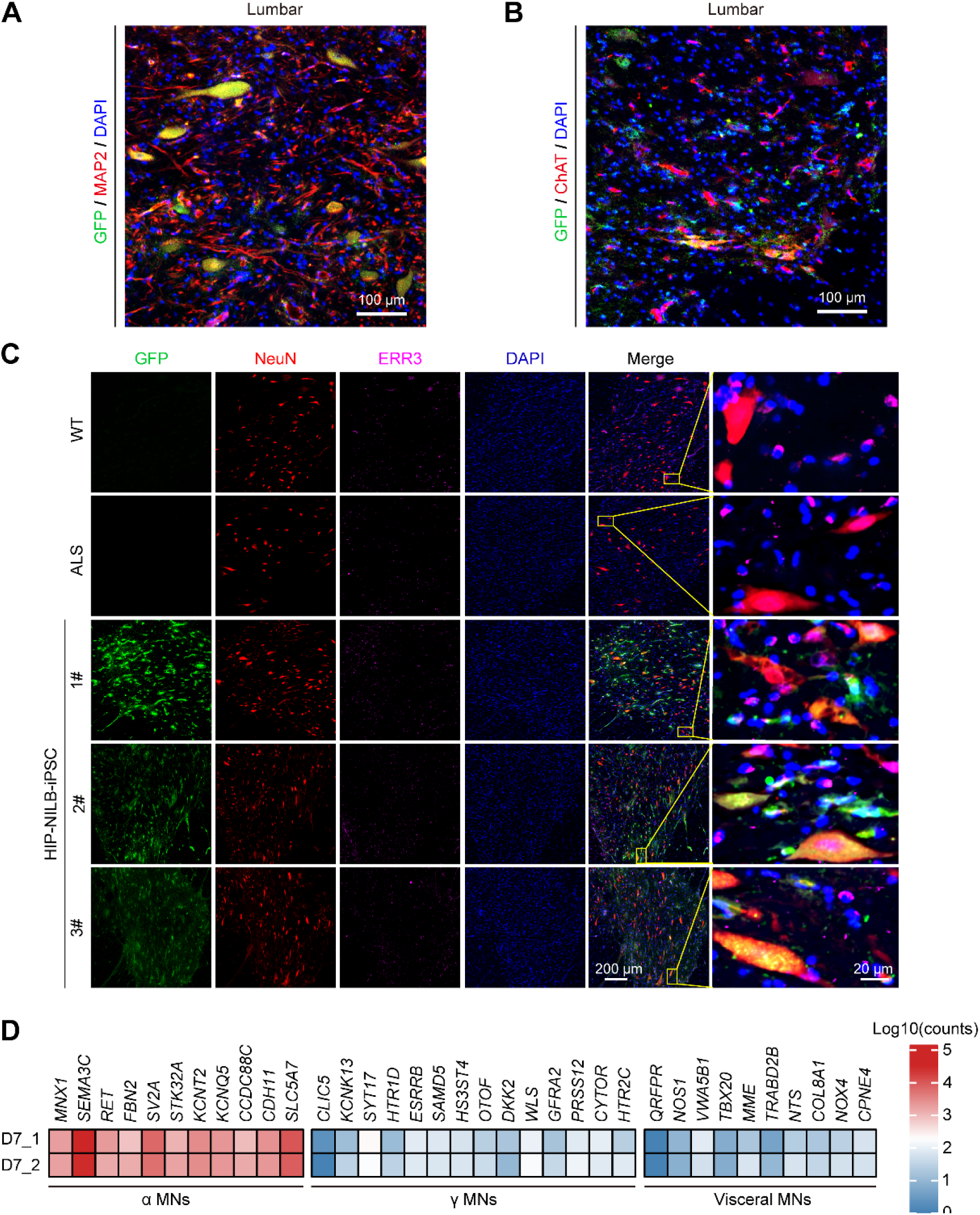
HIP-NILB-iPSCs mainly differentiated into α-MNs in the grafted ALS pigs. (A-B) HIP-NILB-iPSCs derivatives expressed mature neuron marker (A) MAP2 and MNs marker (B) ChAT in HIP-NILB-iPSCs treated ALS pigs with a lower magnification (correspond to Figure 2F). Scale bar, 100 μm. **(C)** Representative images showing HIP-NILB-iPSCs mainly differentiated into NeuN^+^ERR3^-^ α-MNs rather than NeuN^-^ERR3^+^ γ-MNs in all three HIP-NILB-iPSCs treated pigs. Scale bars, 200 μm or 20 μm. **(D)** Heatmap shows the expression pattern of α-MNs, γ-MNs and visceral MNs related genes in Dox-induced HIP-NILB-iPSCs-MNs (Day 7) *in vitro*.

**Fig. S7.**
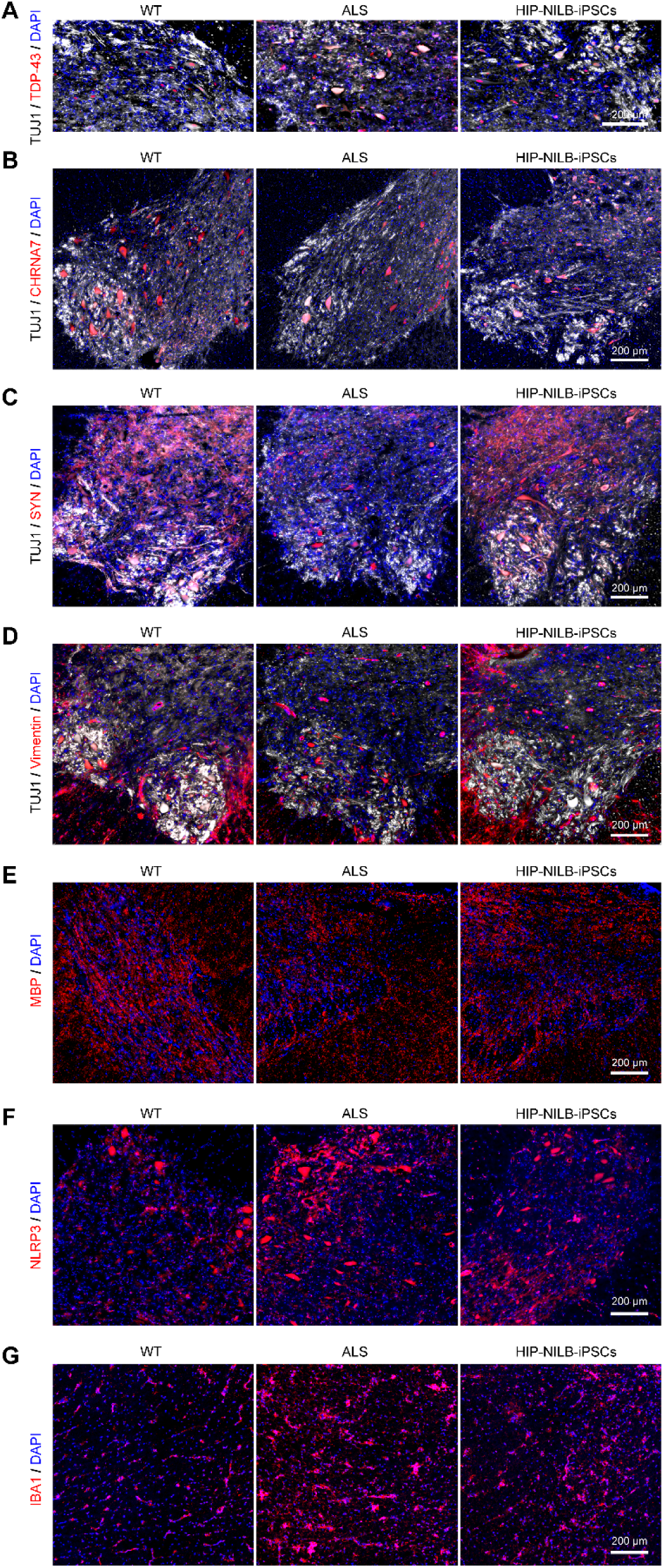
HIP-NILB-iPSCs improved ALS pathology and neural microenvironments in ALS pigs.(A) Representative images showing lower magnification images of TDP-43 in TUJ1^+^ neurons in WT, ALS and HIP-NILB-iPSCs treated pigs (correspond to Figure 2K). Scale bar, 200 μm. **(B-G)** Representative images showing lower magnification images of (B) CHRNA7 (correspond to Fig. 3b), (C) SYN (correspond to Fig. 3C), (D) Vimentin (correspond to Fig. 3D), (E) MBP (correspond to Fig. 3E), (F) NLRP3 (correspond to Fig. 3F), (G) IBA1 (correspond to Fig. 3G), respectively, in the lumbar spinal cord of WT, ALS and HIP-NILB-iPSCs treated pigs. Scale bar, 200 μm.

**Fig. S8.**
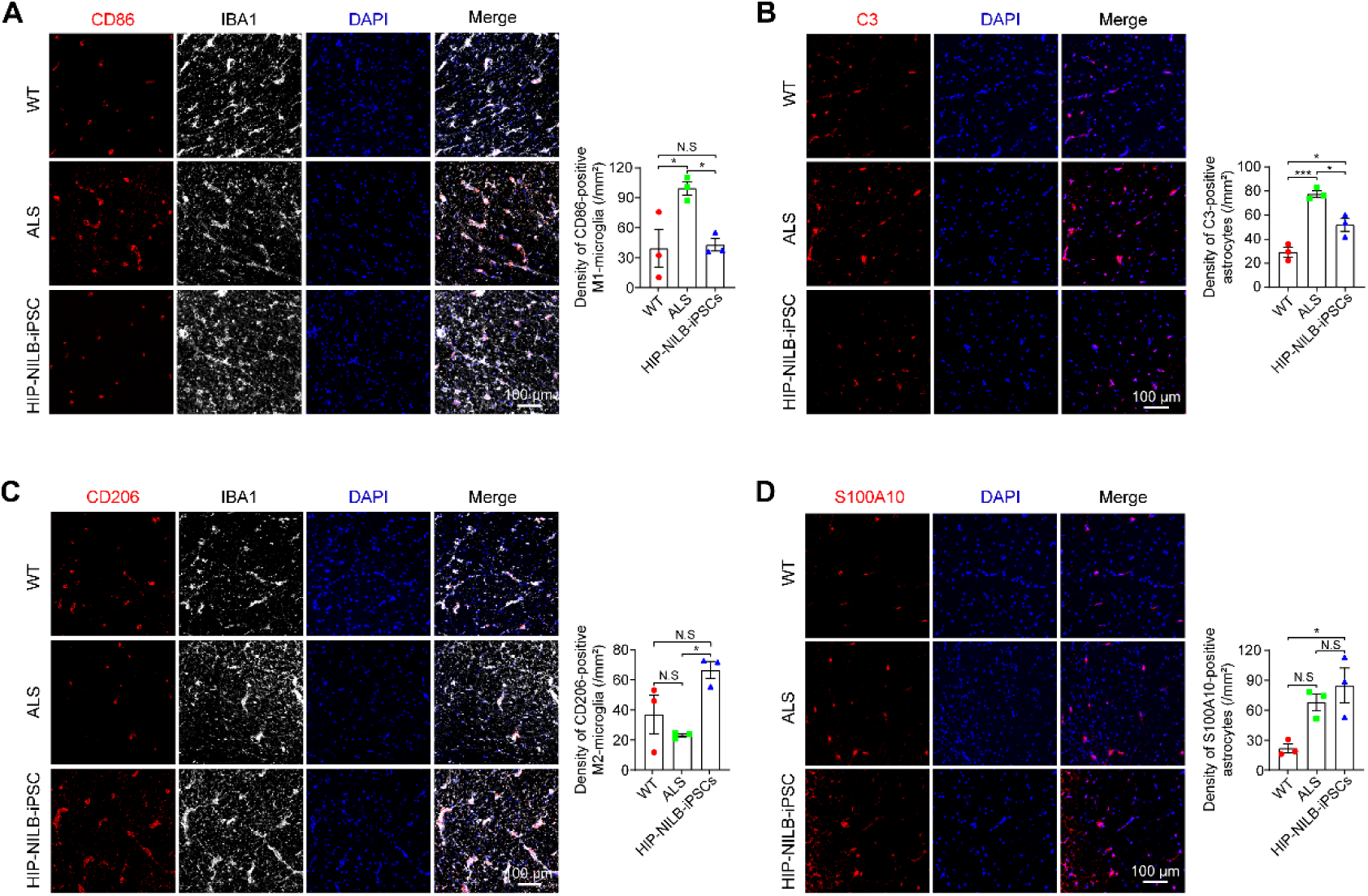
HIP-NILB-iPSCs attenuated neural inflammatory in ALS pigs. (A-B) Representative images of (A) CD86 and (B) C3 in WT, ALS and HIP-NILB-iPSCs treated pigs and the quantifications (n=3). Scale bar, 100 μm. **(C-D)** Representative images of (C) CD206 and (D) S100A10 in WT, ALS and HIP-NILB-iPSCs treated pigs and the quantifications (n=3). Scale bar, 100 μm. The data were analyzed by one-way ANOVA followed by Tukey’s multiple comparisons. Data was presented as mean ± SEM. \**p* < 0.05, \*\**p* < 0.01, \*\*\**p* < 0.001, N.S, not significant.

**Fig. S9.**
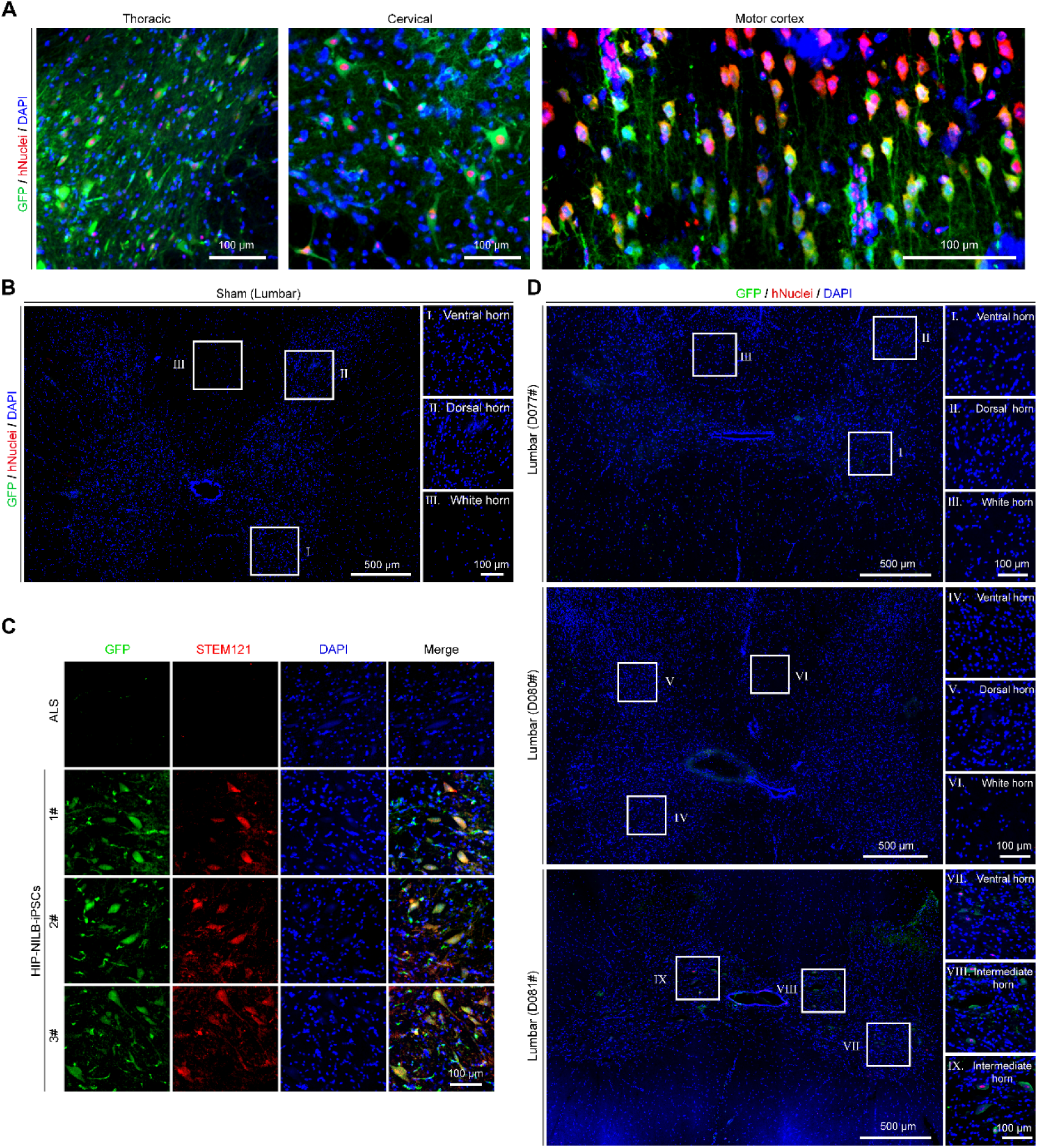
HIP-NILB-iPSCs survived in the CNS of ALS rabbits. **(A)** GFP^+^hNuclei^+^ HIP-NILB-iPSCs migrated and survived in the thoracic, cervical spinal cords and motor cortex of ALS rabbits (lower magnification images corresponding to Figure 4B). Scale bar, 100 μm. **(B)** Representative images of GFP and hNuclei staining in lumbar spinal cord sections of sham ALS rabbits. Scale bars, 500 μm or 100 μm. **(C)** Representative images of GFP and STEM121 staining in ALS and three HIP-NILB-iPSCs treated rabbits. Scale bar, 100 μm. **(D)** Representative images showing NILB-hiPSCs rarely survive in the lumbar spinal cords of three ALS rabbits. Scale bars, 500 μm or 100 μm.

**Fig. S10.**
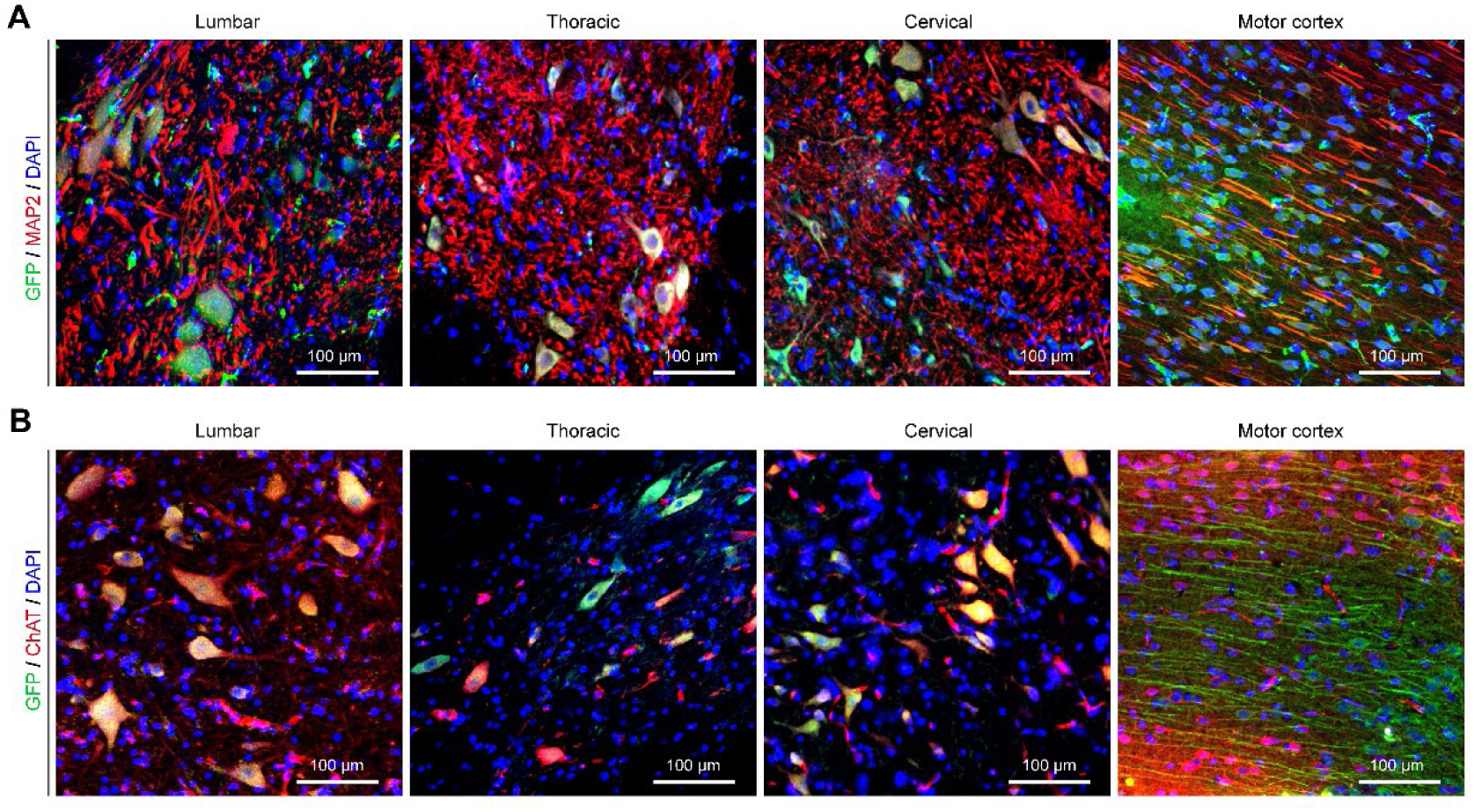
HIP-NILB-iPSCs-MNs migrated in the CNS of ALS rabbits. **(A)** Representative images of HIP-NILB-iPSCs derivatives expressed mature neuron marker MAP2 in the lumbar, thoracic, cervical spinal cords and motor cortex in ALS rabbits with lower magnification (corresponding to Fig. 4G). Scale bar, 100 μm. **(B)** Representative images of HIP-NILB-iPSCs derivatives expressed MNs-marker ChAT in the lumbar, thoracic, cervical spinal cords and motor cortex in ALS rabbits with lower magnification (corresponding to Fig. 4H). Scale bar, 100 μm.

**Fig. S11.**
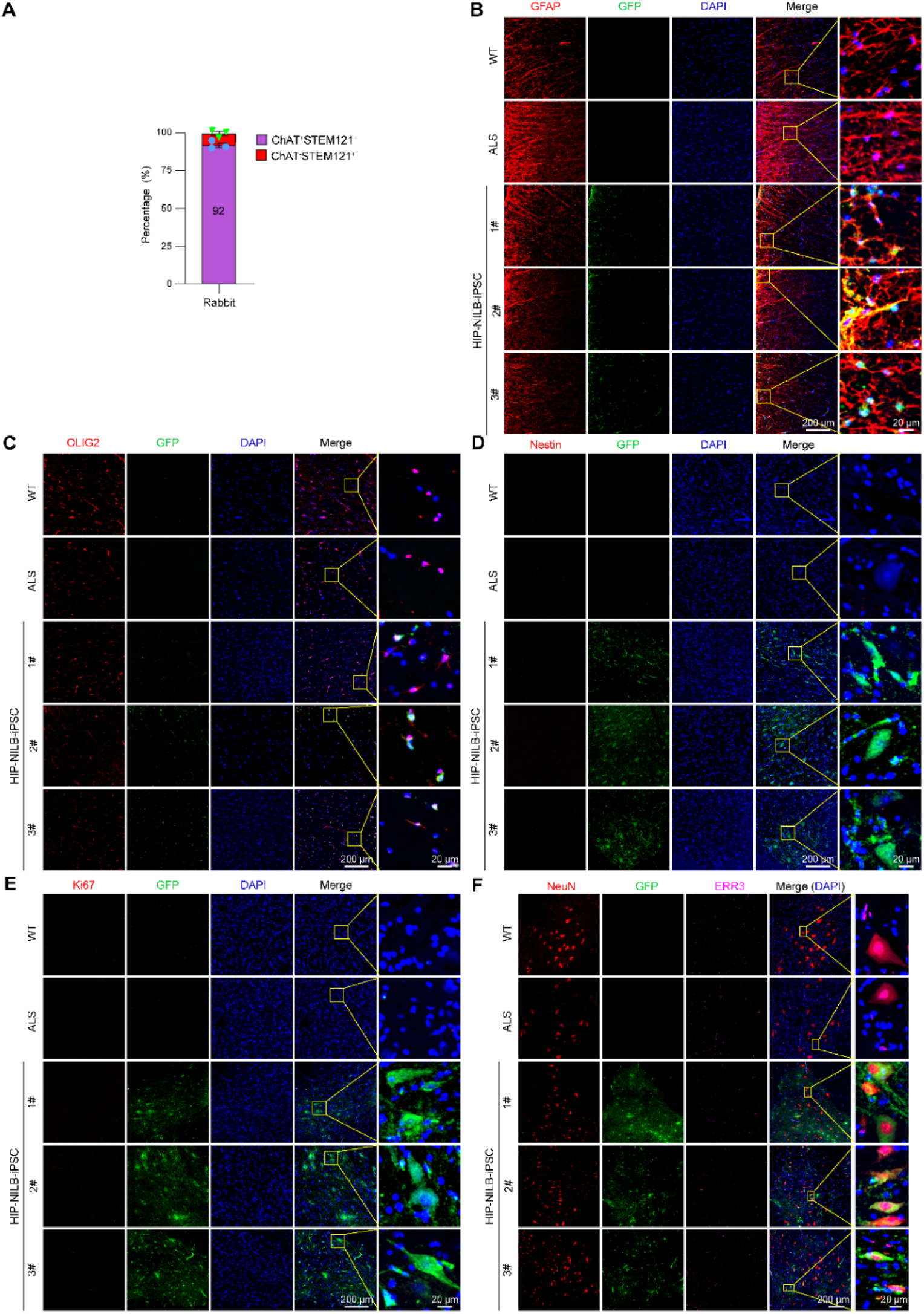
HIP-NILB-iPSCs mainly differentiated into MNs in ALS rabbits. **(A)** Quantitative percentage of STEM121 positive HIP-NILB-iPSCs-derived MNs in the grafted ALS rabbits (n=3). **(B-E)** Representative images showing the expression of (B) GFAP, (C) OLIG2, (D) Nestin and (E) Ki67 in the lumbar spinal cords of WT, ALS and three HIP-NILB-iPSCs-treated rabbits. Scale bars, 200 μm or 20 μm. **(F)** Representative immunofluorescence images and indicated high magnification zones showing HIP-NILB-iPSCs-derived α-MNs (NeuN^+^ERR3^-^) and γ-MNs (NeuN^-^ERR3^+^) in WT, ALS and three HIP-NILB-iPSCs-treated rabbits. Scale bars, 200 μm or 20 μm.

**Fig. S12.**
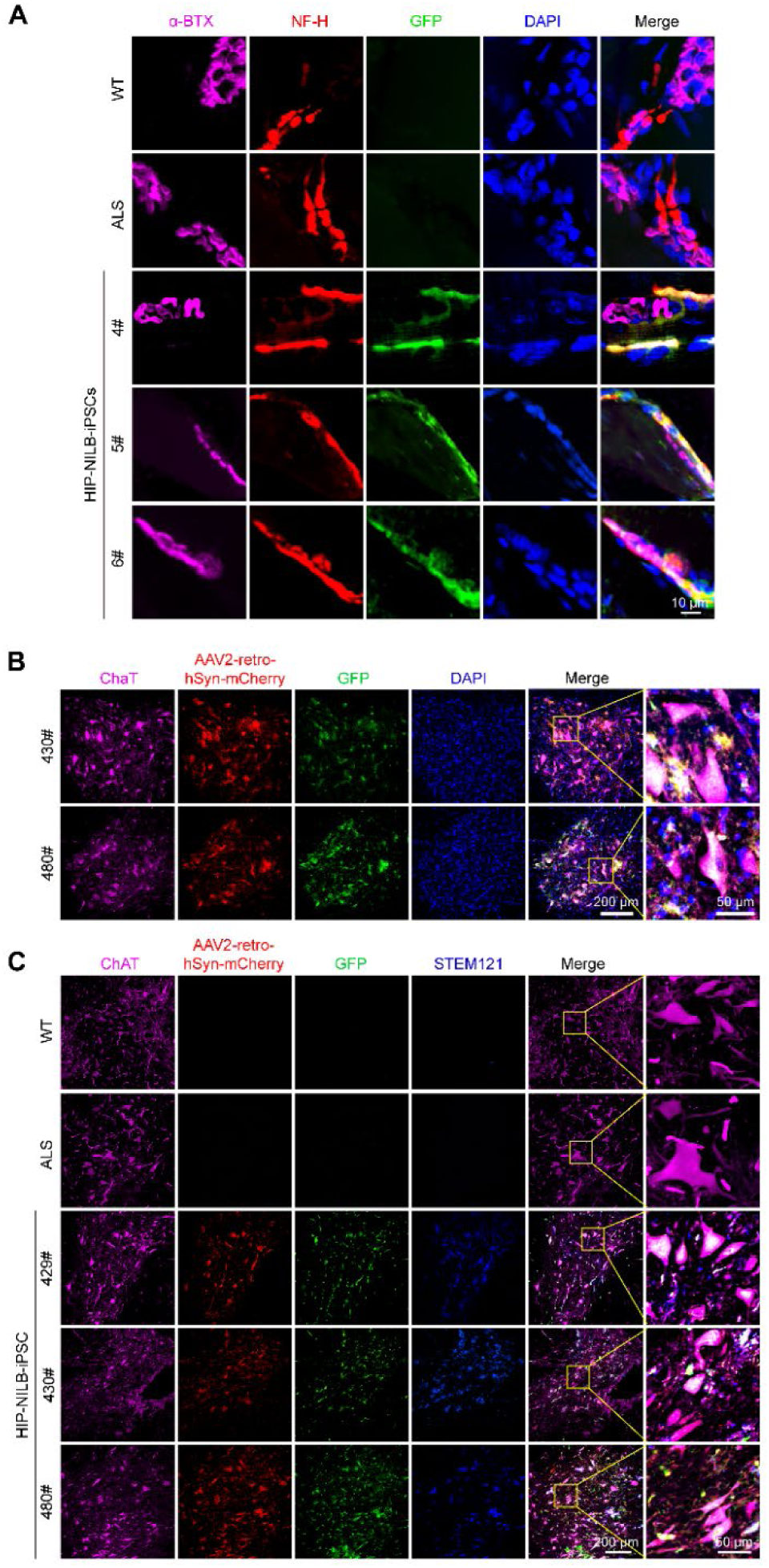
HIP-NILB-iPSCs-MNs integrated into host nerve circuits and formed new NMJs in ALS rabbits. **(A)** Representative immunofluorescence images of α-BTX^+^NF-H^+^GFP^+^DAPI^+^ showing reformed NMJs between HIP-NILB-iPSCs-derived and host gastrocnemius muscle fibers in HIP-NILB-iPSCs-treated three ALS rabbits (Rabbit 4#, 5# and 6#). Scale bar, 10 μm. **(B)** Representative images of retrograde tracing using AAV2-retro-hSyn-mcherry virus showing HIP-NILB-iPSCs-MNs (mCherry^+^ChAT^+^GFP^+^) in the lumbar spinal cord of HIP-NILB-iPSCs treated rabbits (Rabbit 430# and 480#). Scale bars, 200 μm or 50 μm. **(C)** Representative images showing STEM121^+^ HIP-NILB-iPSCs-MNs labeled by mcherry in the lumbar spinal cords of grafted rabbits (Rabbit 429#, 430# and 480#) (2 months post-transplantation) after gastrocnemius intramuscular injection rAAV2-retro-hSyn-mcherry viruses for 1 month. Scale bars, 200 μm or 50 μm.

**Fig. S13.**
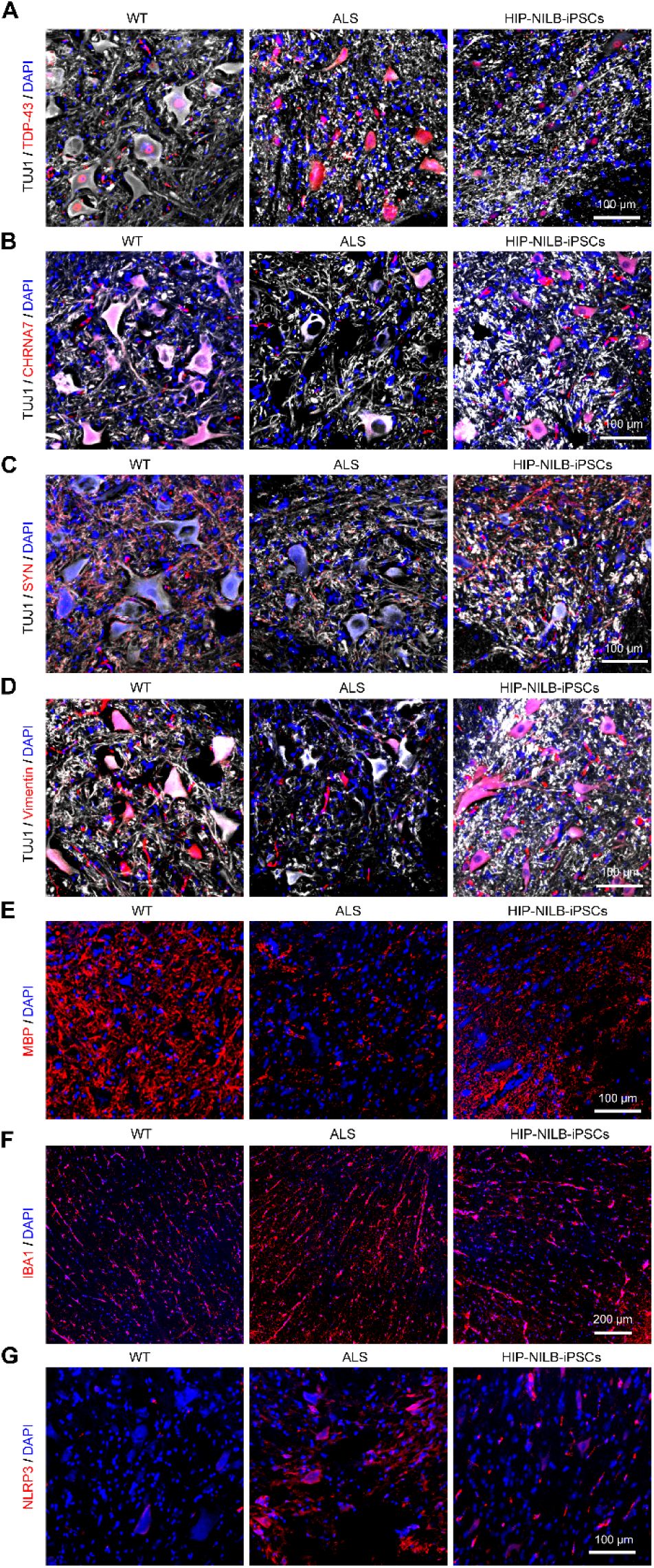
HIP-NILB-iPSCs improved ALS pathology and neural microenvironments in ALS rabbits. **(A)** Representative images showing lower magnification images of TDP-43 corresponding to Fig. 6E. Scale bar, 100 μm. **(B-G)** Representative images showing lower magnification images of (B-E) CHRNA7, SYN, Vimentin and MBP (corresponding to Fig. 6M), and (F-G) IBA1 and NLRP3 (corresponding to Fig. 6N), respectively. Scale bars, 200 μm or 100 μm.

**Fig. S14.**
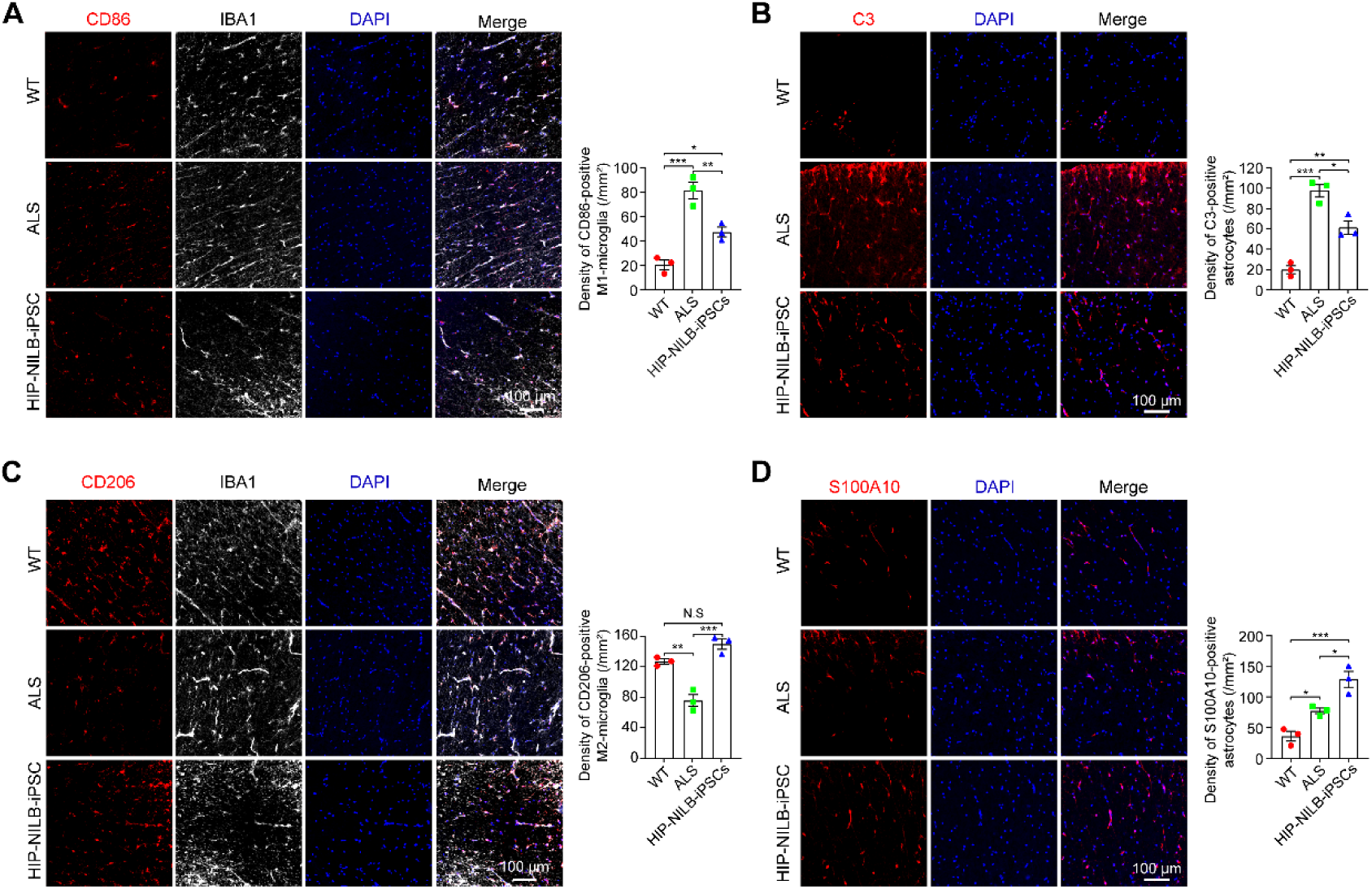
HIP-NILB-iPSCs attenuated neural inflammatory in ALS rabbits. (A-B) Representative images of (A) CD86 and (B) C3 in WT, ALS and HIP-NILB-iPSCs treated rabbits and the quantifications (n=3). Scale bar, 100 μm. **(C-D)** Representative images of (C) CD206 and (D) S100A10 in WT, ALS and HIP-NILB-iPSCs treated rabbits and the quantifications (n=3). Scale bar, 100 μm. The data were analyzed by one-way ANOVA followed by Tukey’s multiple comparisons. Data was presented as mean ± SEM. \**p* < 0.05, \*\**p* < 0.01, \*\*\**p* < 0.001, N.S, not significant.

**Table S1.**
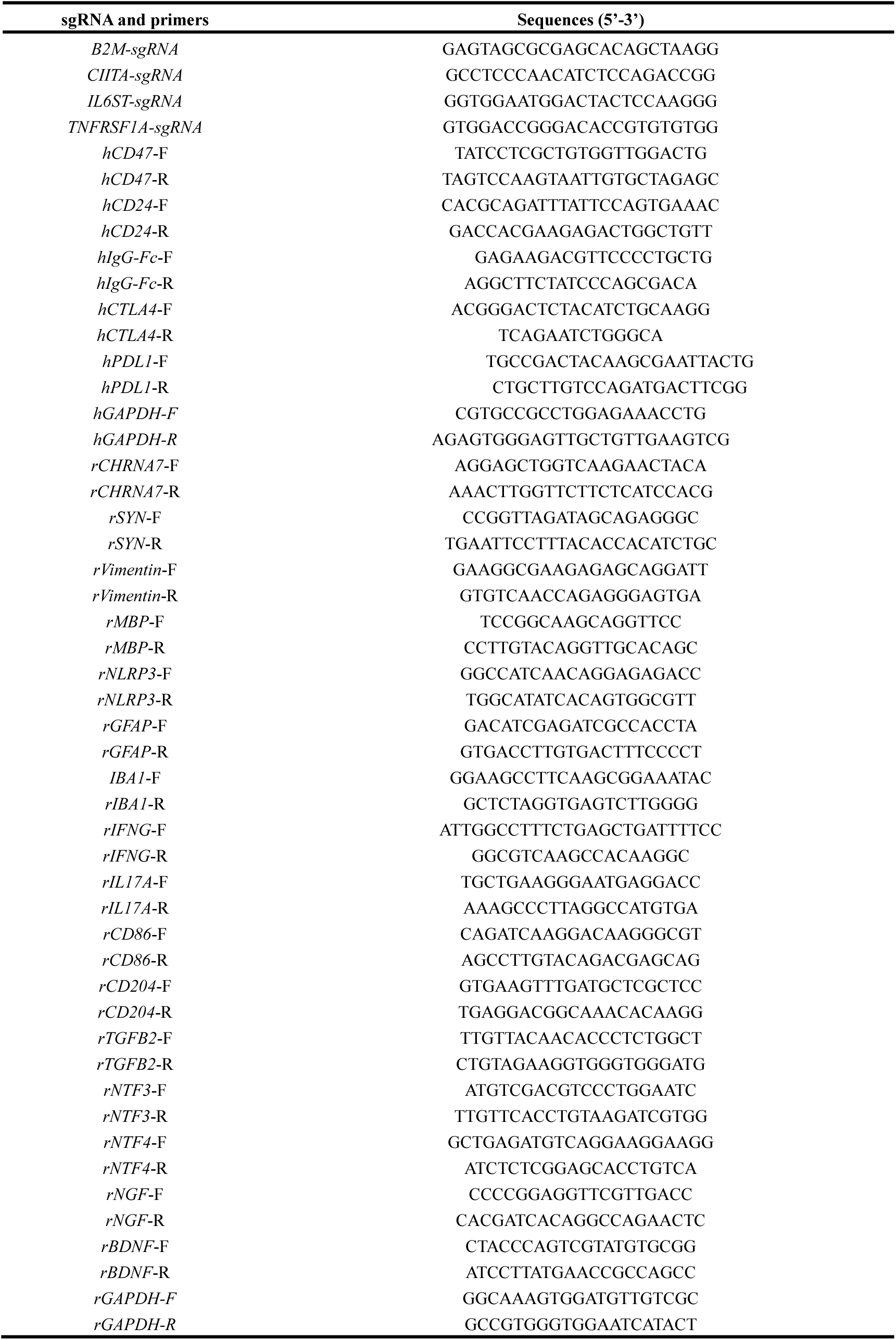
sgRNA sequences and real-time PCR primers were used.

