## supplementary figures and table for "Hypoimmunogenic human motor neurons induced from iPSCs in vivo substantially ameliorate ALS disease in large animal models"

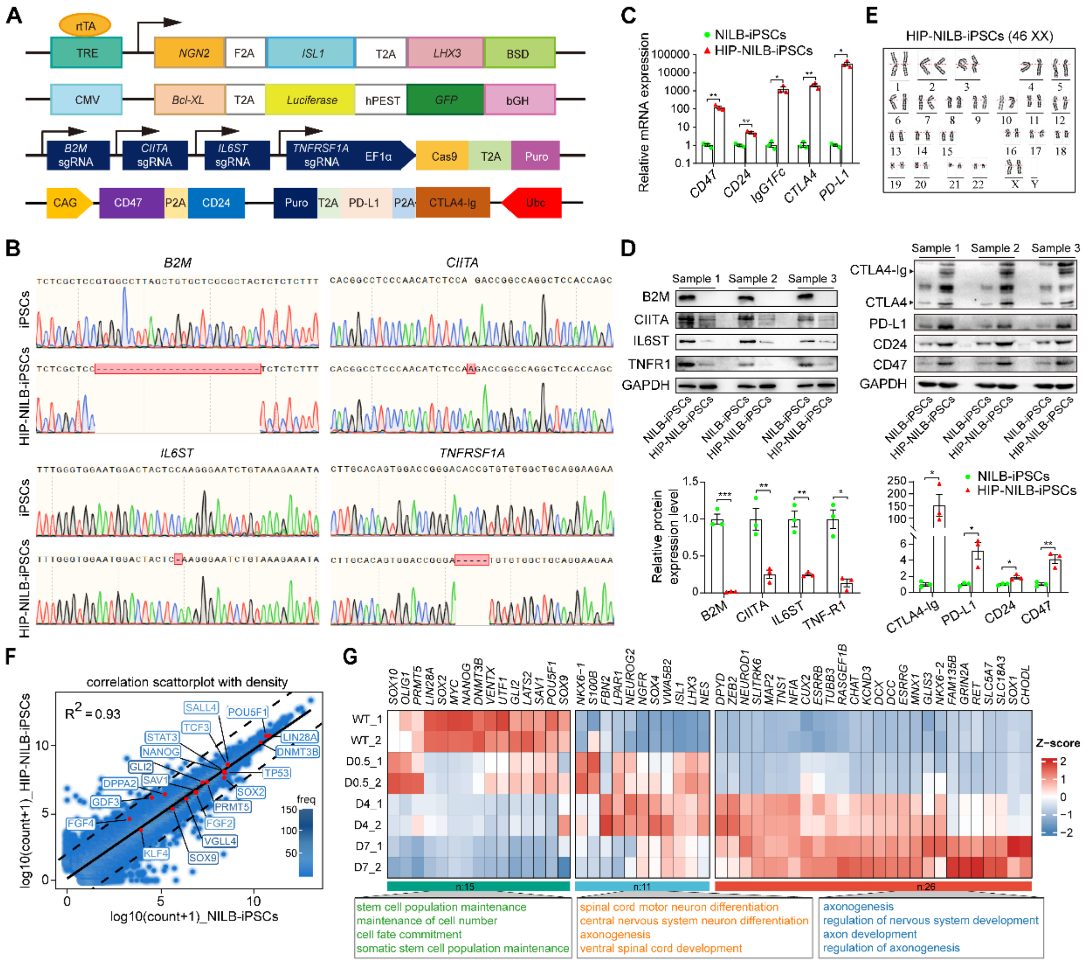

**Fig. S1. Engineered HIP-NILB-iPSCs via multiplex genetic-editing.**

**(A)** Schematic illustration of multiplex genetic-editing strategies for engineered HIP-NILB-iPSCs. **(B)** Indel analysis by Sanger sequencing for *B2M*, *CIITA*, *IL6ST* and *TNFRSF1A* in HIP-NILB-iPSCs. **(C)** Real-time PCR analysis of *CD47*, *CD24*, *IgG1-Fc*, *CTLA4* and *PD-L1* in HIP-NILB-iPSCs. **(D)** Western blot analysis and relative density quantification of B2M, CIITA, IL6ST, TNFR1, CTLA4-Ig, PD-L1, CD24 and CD47 in HIP-NILB-iPSCs of three independent samples. CIITA was detected in NILB-iPSCs and HIP-NILB-iPSCs upon IFN-γ and TNF-α treatment for 48 h. **(E)** Karyotyping of HIP-NILB-iPSCs. **(F)** Scatter plot comparison of transcriptomes for HIP-NILB-iPSCs and NILB-iPSCs. The raw counts for each gene were transformed to log10 value, and genes with more than 1 count in each sample were shown. The pluripotent-related genes were labeled in the diagram. Dashed lines depict the 10-fold changes. The $R^2$ was determined by Pearson's correlation. The NILB-iPSCs were used as a control. **(G)** Heatmap showing expression level of stem cells-related and neuron development-related genes together with the enrichment analysis in WT iPSCs and HIP-NILB-iPSCs upon Dox induced for 0.5-, 4- and 7- days. Data are mean ± SEM; p values were determined using a two-tailed, unpaired Student's *t-test*; *$p < 0.05$; **$p < 0.01$.

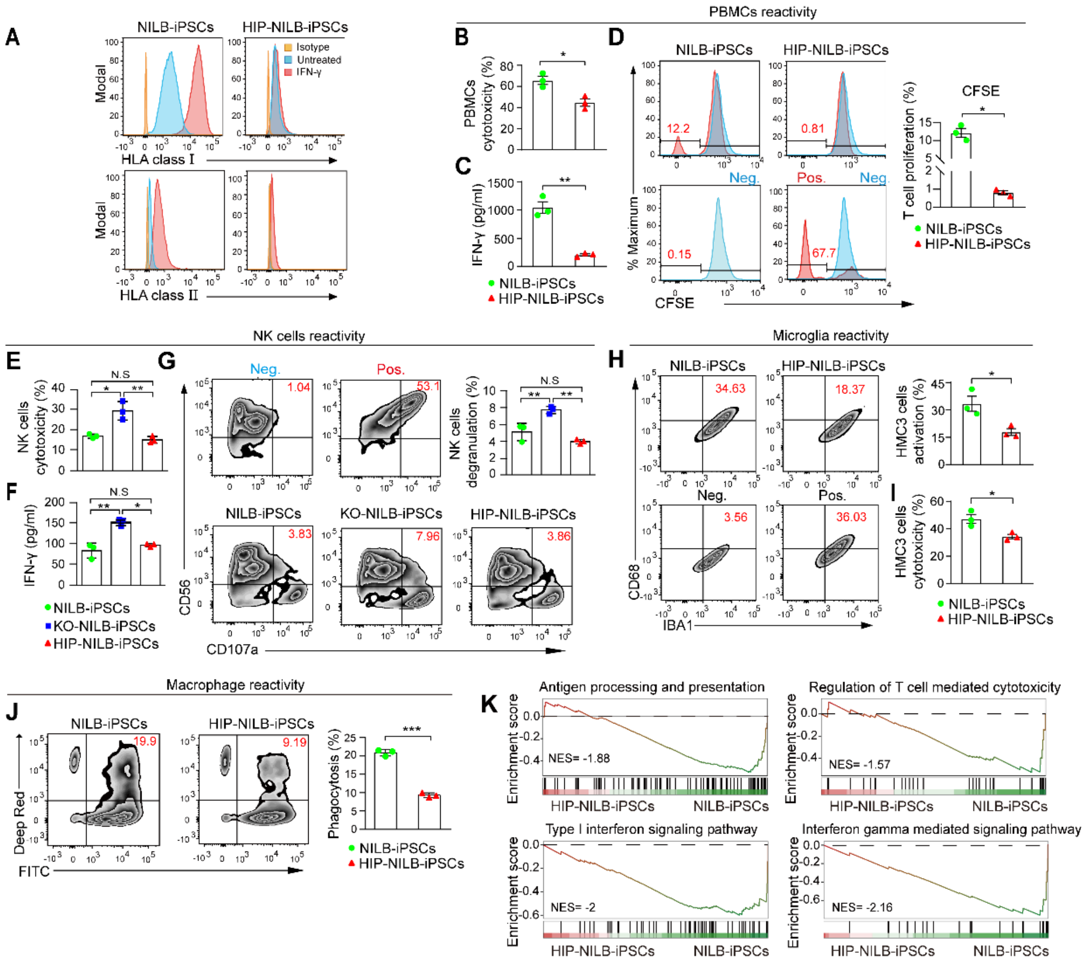


**Fig. S2. Immune response against HIP-NILB-iPSCs *in vitro.***

**(A)** HLA class I and II expression in HIP-NILB-iPSCs with or without IFN-γ treatment. Isotype is a negative control with matched primary antibody. **(B)** PBMCs cytotoxicity against HIP-NILB-iPSCs by measuring LDH release. **(C)** ELISA assay for IFN-γ secretion in PBMCs cocultured with HIP-NILB-iPSCs. **(D)** CFSE analysis of T cells ($CD3^+$) proliferation in PBMCs when cocultured with HIP-NILB-iPSCs. PBMCs is negative control (Neg.), PBMCs activated by PHA is positive control (Pos.). **(E)** Primary NK cells cytotoxicity against HIP-NILB-iPSCs by measuring LDH release. **(F)** ELISA assay for IFN-γ secretion in Primary NK cells cocultured with HIP-NILB-iPSCs. **(G)** NK degranulation assay by quantifying CD107a surface expression in $CD56^+$ NK cells cocultured with HIP-NILB-iPSCs. NK cells alone as Neg., NK cells treated with PHA as Pos.. **(H)** Activation of HMC3 cells ($CD68^+$) in $IBA1^+$ when incubated with HIP-NILB-iPSCs. HMC3 alone as Neg., LPS-stimulated HMC3 as Pos.. **(I)** HMC3 cells cytotoxicity against HIP-NILB-iPSCs by measuring LDH release. **(J)** The phagocytic activity of THP-1 derived macrophages against HIP-NILB-iPSCs is characterized by the percentage of double positive of Deep Red and FITC. **(K)** Gene set enrichment analysis (GSEA) shows the enriched signaling pathway in HIP-NILB-iPSCs against NILB-iPSCs. NES represents the normalized enrichment score. All experiments were independently duplicated three times. Data are mean ± SEM, *p* values were determined using a two-tailed, unpaired Student's *t-test* (b, c, e, f, g, h, i, j) or Welch's t-test (D), $*p < 0.05$, $**p < 0.01$, $***p < 0.001$, N.S, not significant.

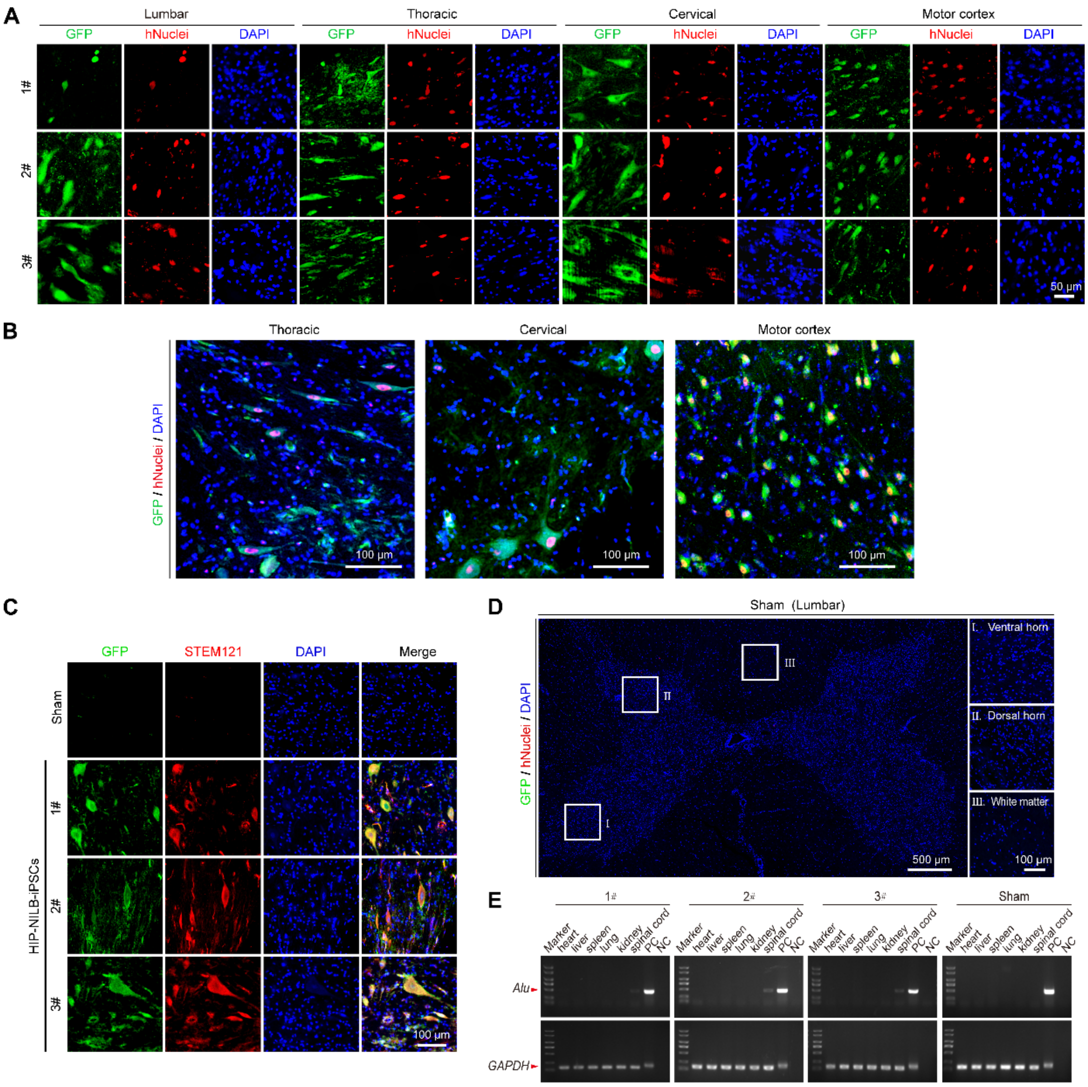


**Fig. S3. HIP-NILB-iPSCs survived in the CNS of ALS pigs.**
**(A)** The single channel images corresponding to Figure 2B. Scale bar, 50 µm. **(B)** $GFP^{+}hNuclei^{+}$ HIP-NILB-iPSCs migrated and survived in the thoracic, cervical spinal cords and motor cortex of ALS pig (2#) with a lower magnification (corresponding to Figure 2B). Scale bar, 100 µm. **(C)** Representative confocal images of GFP and STEM121 staining in HIP-NILB-iPSCs treated pigs. Scale bar, 100 µm. **(D)** Representative images of GFP and hNuclei staining in lumbar spinal cord sections of sham ALS pig. Scale bars, 500 µm or 10 µm. **(E)** *Alu* PCR analysis of various organs and spinal cords in treated pigs and the sham control.

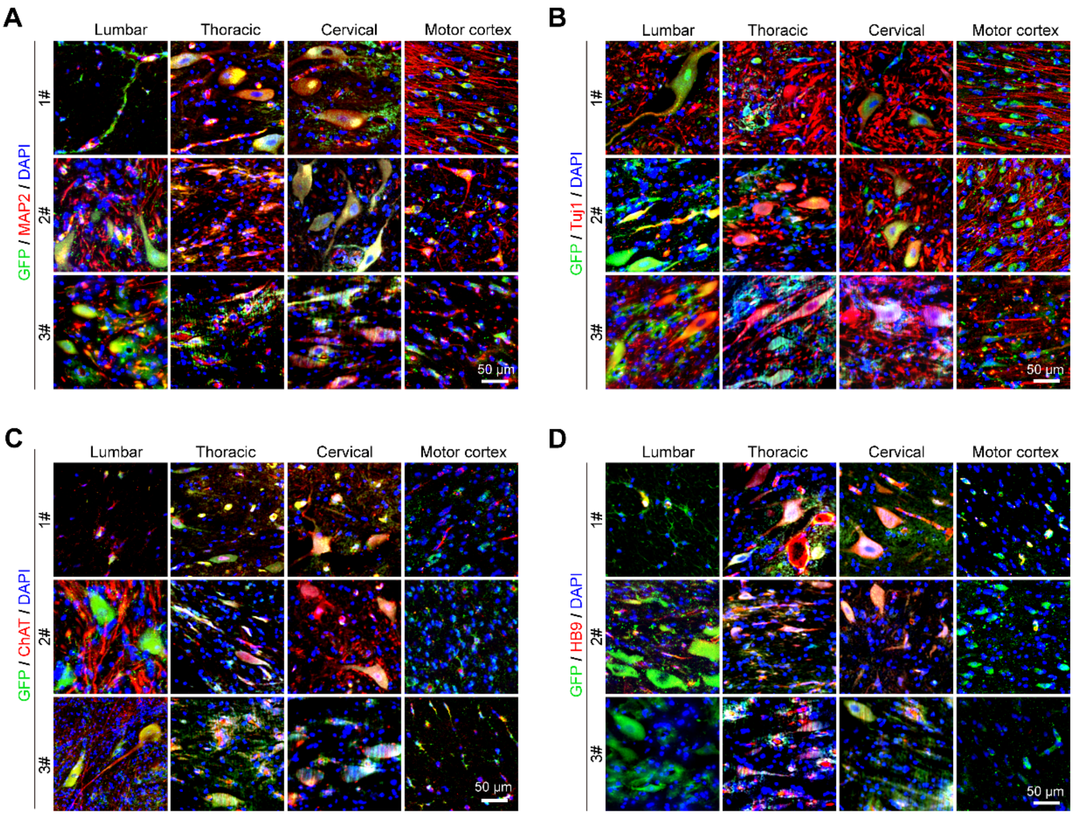


**Fig. S4. Widespread distribution of HIP-NILB-iPSCs-MNs.**

**(A-D)** Representative immunofluorescence images showed the expression of neuron markers (A) MAP2 and (B) TUJ1, and MNs markers (C) ChAT and (D) HB9 of HIP-NILB-iPSCs-derived MNs in lumbar, thoracic, cervical spinal cords, and motor cortex in three HIP-NILB-iPSCs treated ALS pigs. Scale bar, 50 μm.

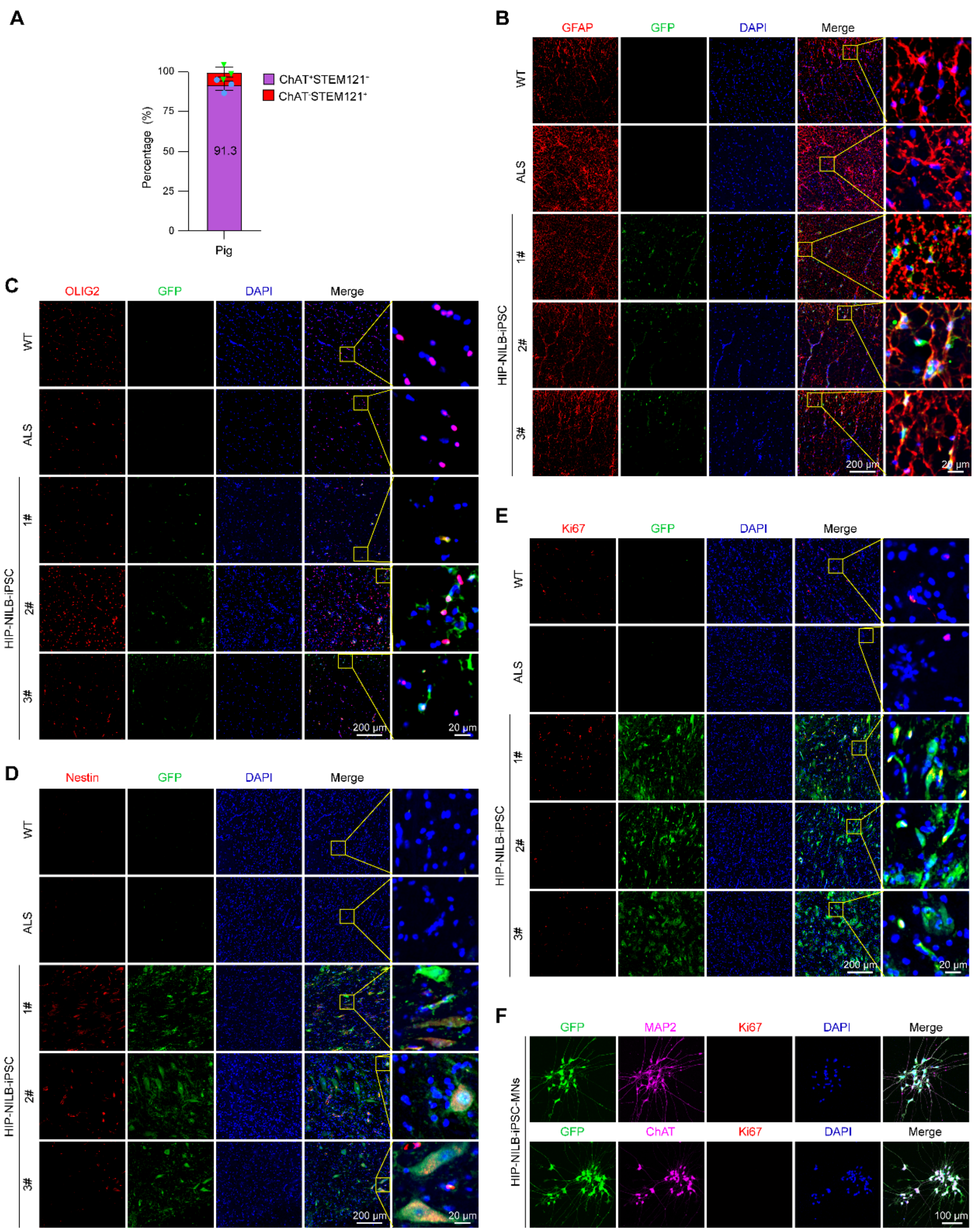


**Fig. S5. HIP-NILB-iPSCs mainly differentiated into MNs in ALS pigs.**

**(A)** Quantitative percentage of STEM121 positive HIP-NILB-iPSCs-derived MNs in HIP-NILB-iPSCs treated ALS pigs (n=3). **(B-D)** Representative images showing very few HIP-NILB-iPSCs differentiated into (B) astrocytes, (C) oligodendrocyte and (D) neural stem cells in the lumbar spinal cords of three HIP-NILB-iPSCs treated ALS pigs. Scale bars, 200 μm or 20 μm. **(E)** Representative images of Ki67 staining in the lumbar spinal cords of HIP-NILB-iPSCs grafted ALS pigs. Scale bars, 200 μm or 20 μm. **(F)** Representative images show no Ki67$^{+}$ cells in HIP-NILB-iPSCs MNs *in vitro*. HIP-NILB-iPSCs were induced upon Dox for 5 days. Scale bar, 100 μm.

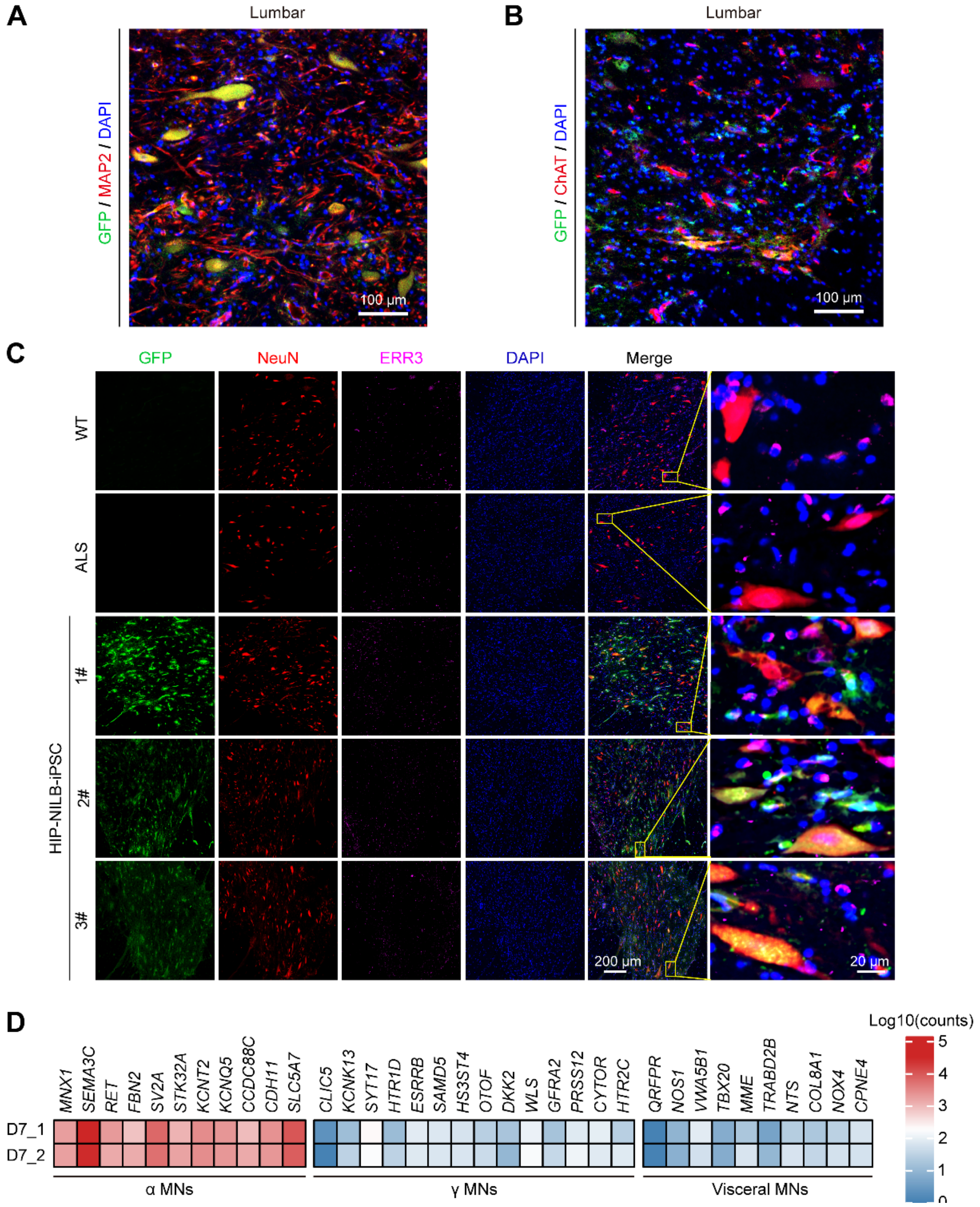


**Fig. S6. HIP-NILB-iPSCs mainly differentiated into α-MNs in the grafted ALS pigs.**

**(A-B)** HIP-NILB-iPSCs derivatives expressed mature neuron marker (A) MAP2 and MNs marker (B) ChAT in HIP-NILB-iPSCs treated ALS pigs with a lower magnification (correspond to Figure 2F). Scale bar, 100 μm. **(C)** Representative images showing HIP-NILB-iPSCs mainly differentiated into $NeuN^{+}ERR3^{-}$ α-MNs rather than $NeuN^{-}ERR3^{+}$ γ-MNs in all three HIP-NILB-iPSCs treated pigs. Scale bars, 200 μm or 20 μm. **(D)** Heatmap shows the expression pattern of α-MNs, γ-MNs and visceral MNs related genes in Dox-induced HIP-NILB-iPSCs-MNs (Day 7) *in vitro*.

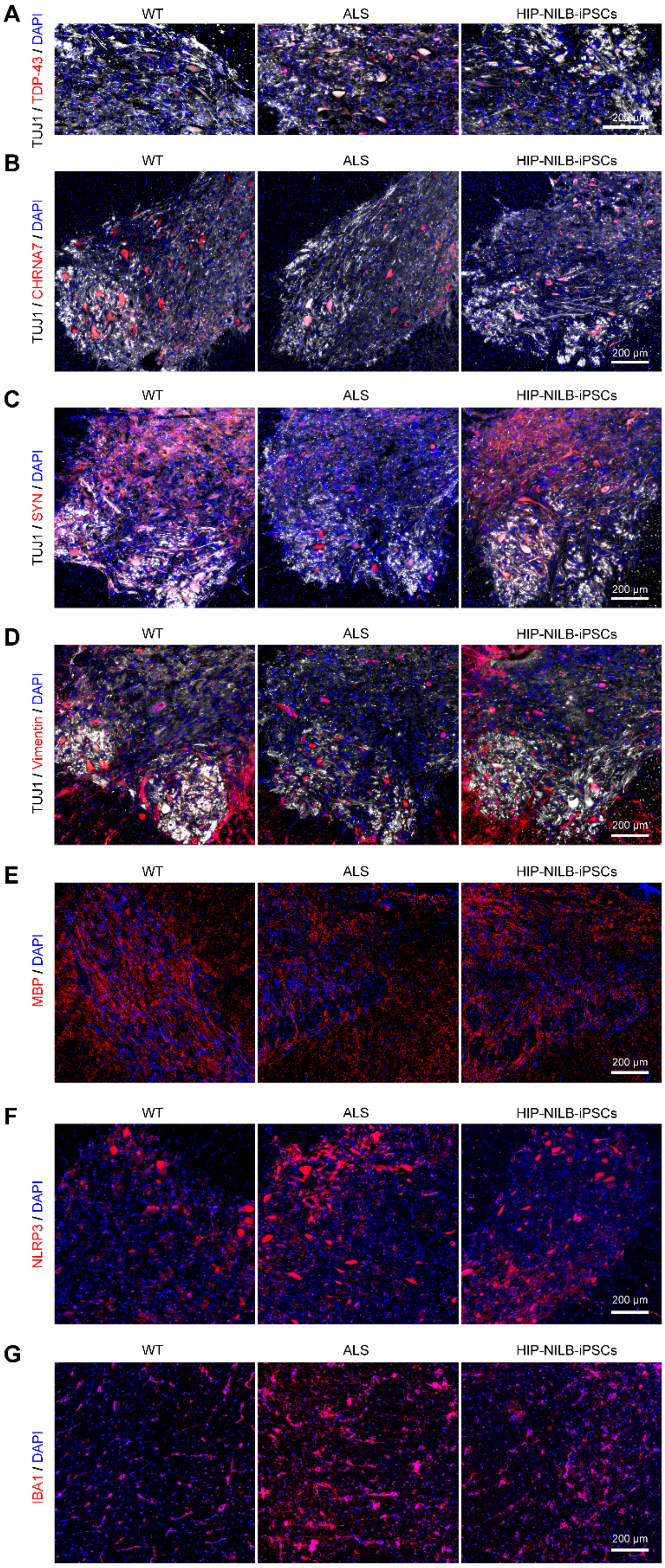


**Fig. S7. HIP-NILB-iPSCs improved ALS pathology and neural microenvironments in ALS pigs.**

**(A)** Representative images showing lower magnification images of TDP-43 in $TUJ1^{+}$ neurons in WT, ALS and HIP-NILB-iPSCs treated pigs (correspond to Figure 2K). Scale bar, 200 µm. **(B-G)** Representative images showing lower magnification images of (B) CHRNA7 (correspond to Fig. 3b), (C) SYN (correspond to Fig. 3C), (D)

Vimentin (correspond to Fig. 3D), (E) MBP (correspond to Fig. 3E), (F) NLRP3 (correspond to Fig. 3F), (G) IBA1 (correspond to Fig. 3G), respectively, in the lumbar spinal cord of WT, ALS and HIP-NILB-iPSCs treated pigs. Scale bar, 200 µm.

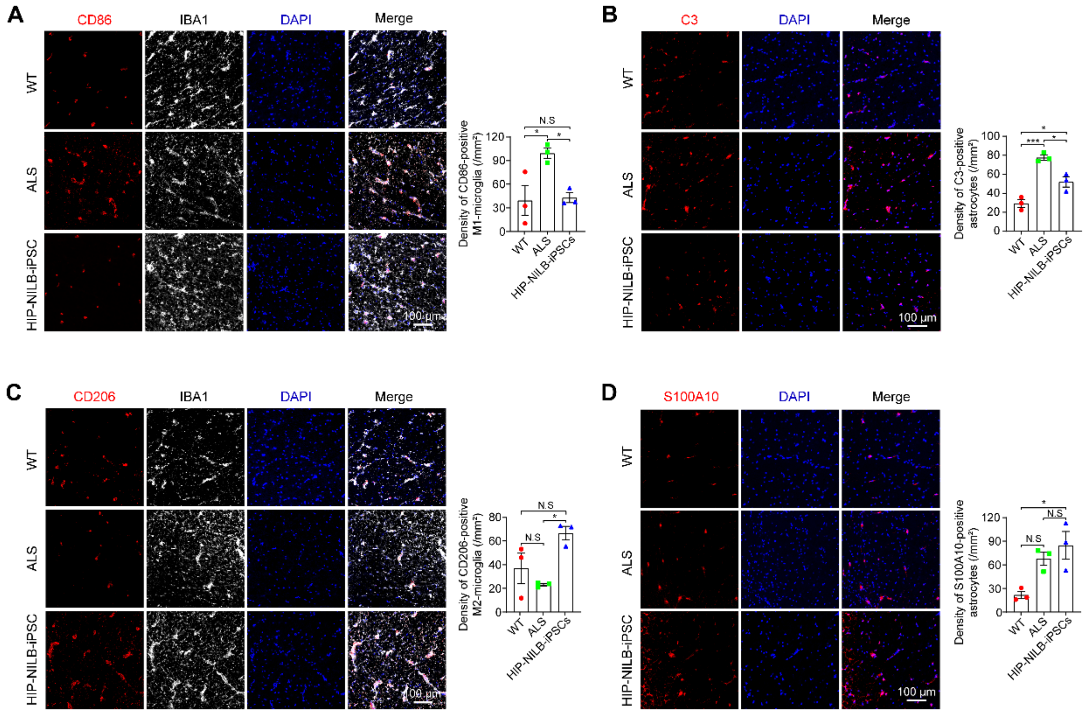


**Fig. S8. HIP-NILB-iPSCs attenuated neural inflammatory in ALS pigs.**
**(A-B)** Representative images of (A) CD86 and (B) C3 in WT, ALS and HIP-NILB-iPSCs treated pigs and the quantifications (n=3). Scale bar, 100 µm. **(C-D)** Representative images of (C) CD206 and (D) S100A10 in WT, ALS and HIP-NILB-iPSCs treated pigs and the quantifications (n=3). Scale bar, 100 µm. The data were analyzed by one-way ANOVA followed by Tukey's multiple comparisons. Data was presented as mean ± SEM. *$p < 0.05$, **$p < 0.01$, ***$p < 0.001$, N.S, not significant.

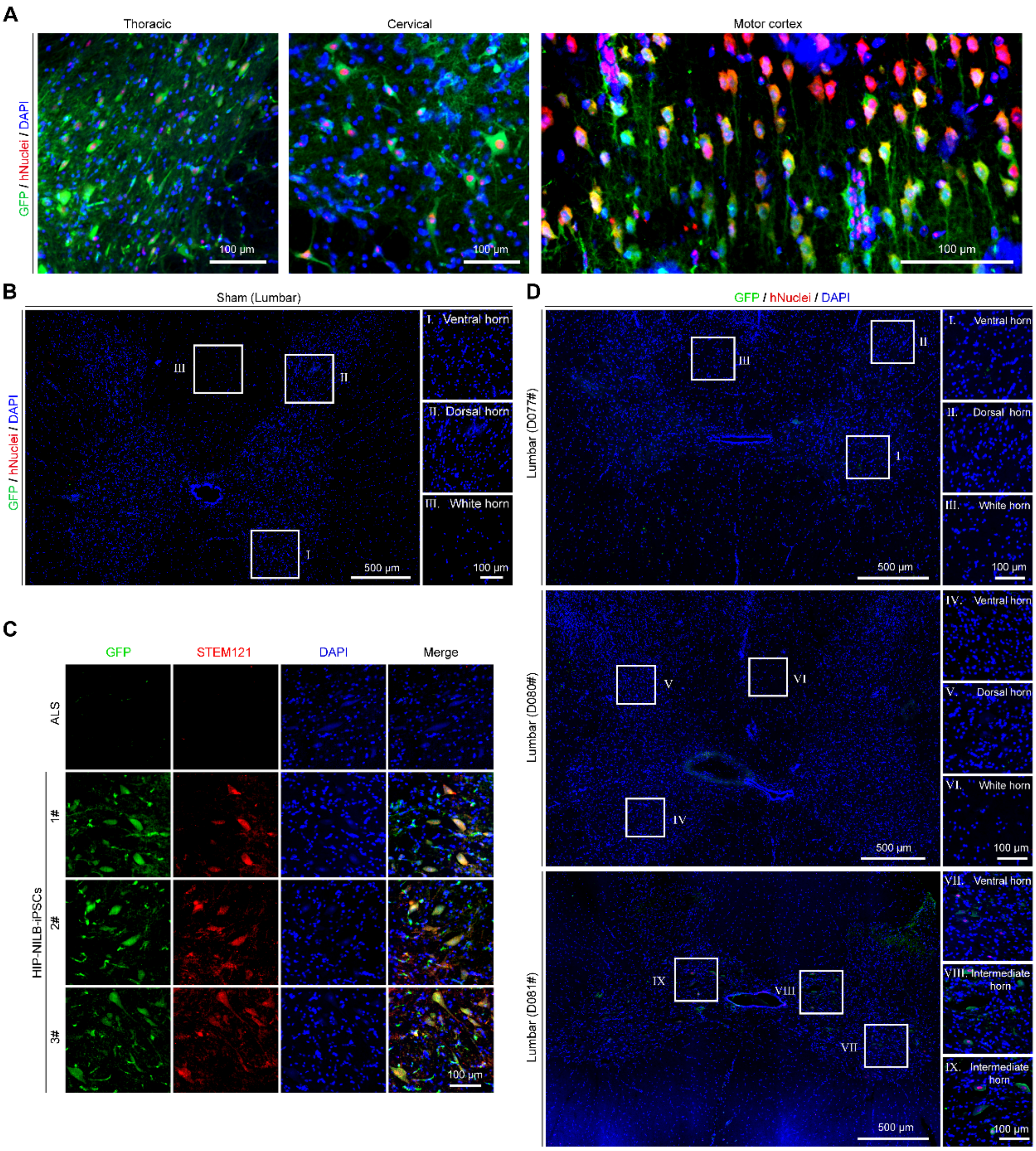


**Fig. S9. HIP-NILB-iPSCs survived in the CNS of ALS rabbits.**

**(A)** $GFP^{+}hNuclei^{+}$ HIP-NILB-iPSCs migrated and survived in the thoracic, cervical spinal cords and motor cortex of ALS rabbits (lower magnification images corresponding to Figure 4B). Scale bar, 100 µm. **(B)** Representative images of GFP and hNuclei staining in lumbar spinal cord sections of sham ALS rabbits. Scale bars, 500 µm or 100 µm. **(C)** Representative images of GFP and STEM121 staining in ALS and three HIP-NILB-iPSCs treated rabbits. Scale bar, 100 µm. **(D)** Representative images showing NILB-hiPSCs rarely survive in the lumbar spinal cords of three ALS rabbits. Scale bars, 500 µm or 100 µm.

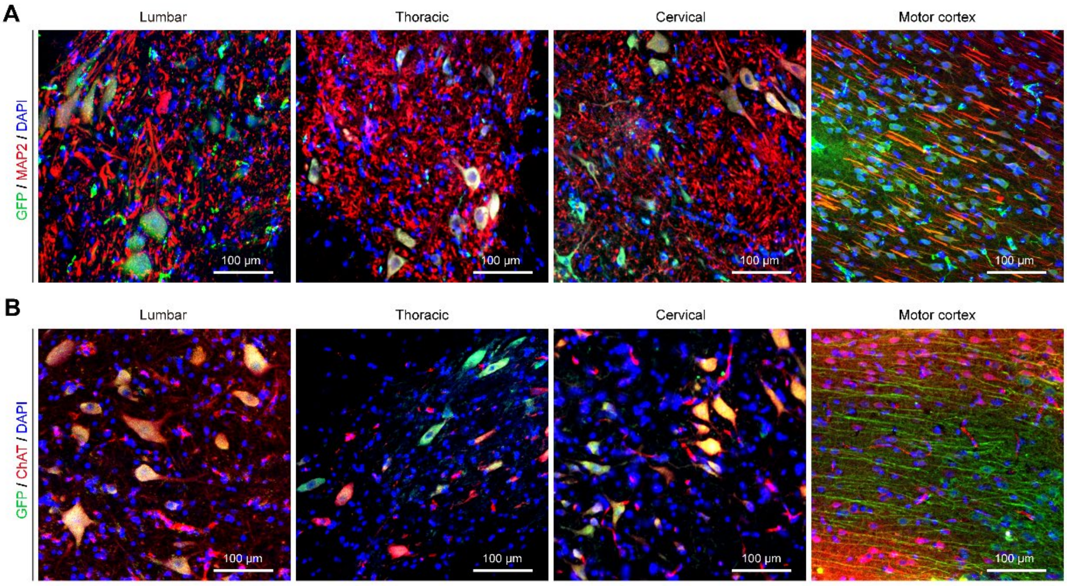


**Fig. S10. HIP-NILB-iPSCs-MNs migrated in the CNS of ALS rabbits.**

**(A)** Representative images of HIP-NILB-iPSCs derivatives expressed mature neuron marker MAP2 in the lumbar, thoracic, cervical spinal cords and motor cortex in ALS rabbits with lower magnification (corresponding to Fig. 4G). Scale bar, 100 μm. **(B)** Representative images of HIP-NILB-iPSCs derivatives expressed MNs-marker ChAT in the lumbar, thoracic, cervical spinal cords and motor cortex in ALS rabbits with lower magnification (corresponding to Fig. 4H). Scale bar, 100 μm.

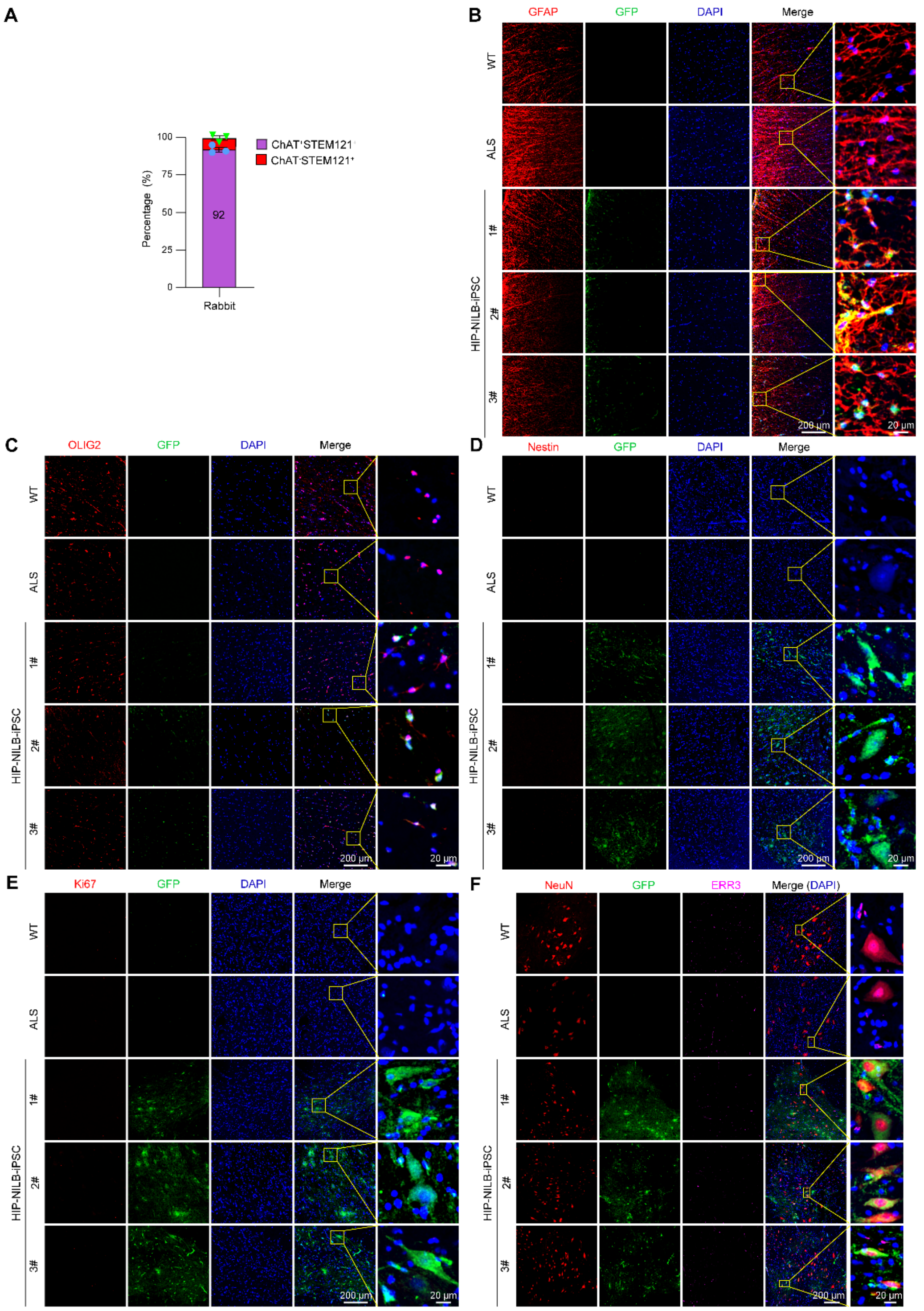


**Fig. S11. HIP-NILB-iPSCs mainly differentiated into MNs in ALS rabbits.**

**(A)** Quantitative percentage of STEM121 positive HIP-NILB-iPSCs-derived MNs in the grafted ALS rabbits (n=3). **(B-E)** Representative images showing the expression of (B) GFAP, (C) OLIG2, (D) Nestin and (E) Ki67 in the lumbar spinal cords of WT, ALS and three HIP-NILB-iPSCs-treated rabbits. Scale bars, 200 μm or 20 μm. **(F)** Representative immunofluorescence images and indicated high magnification zones showing HIP-NILB-iPSCs-derived α-MNs ($NeuN^{+}ERR3^{-}$) and γ-MNs ($NeuN^{-}ERR3^{+}$) in WT, ALS and three HIP-NILB-iPSCs-treated rabbits. Scale bars, 200 μm or 20 μm.

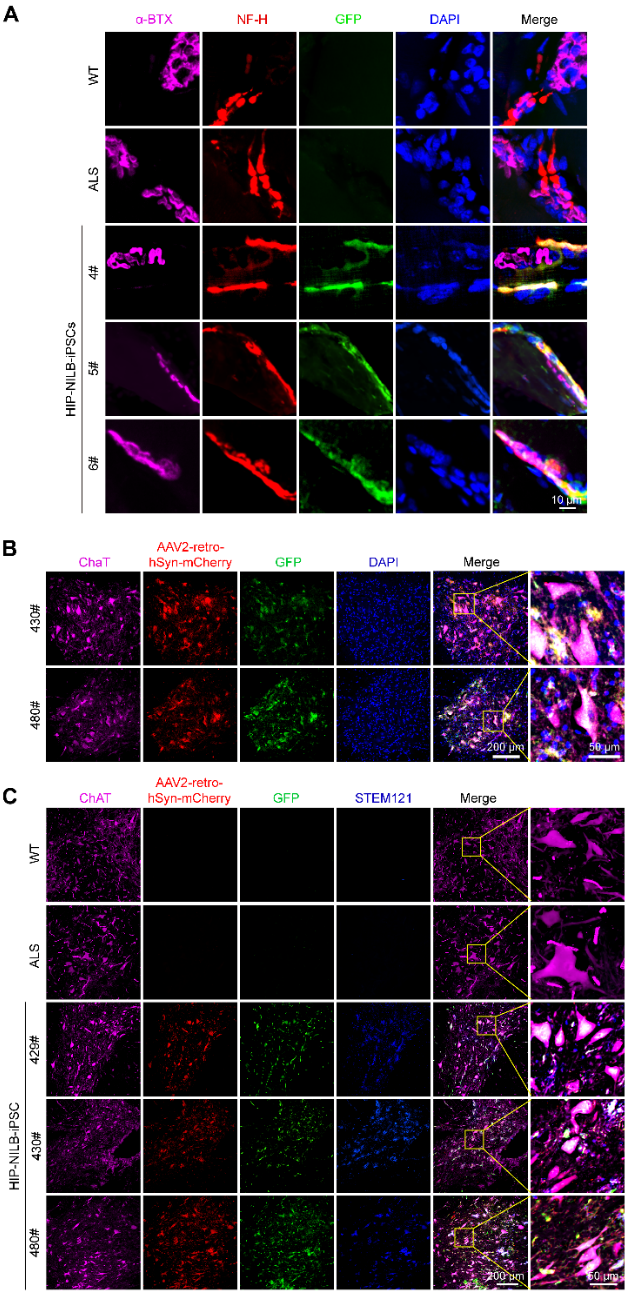


**Fig. S12. HIP-NILB-iPSCs-MNs integrated into host nerve circuits and formed new NMJs in ALS rabbits.**

**(A)** Representative immunofluorescence images of α-BTX$^{+}$NF-H$^{+}$GFP$^{+}$DAPI$^{+}$ showing reformed NMJs between HIP-NILB-iPSCs-derived and host gastrocnemius muscle fibers in HIP-NILB-iPSCs-treated three ALS rabbits (Rabbit 4#, 5# and 6#). Scale bar, 10 μm. **(B)** Representative images of retrograde tracing using AAV2-retro-hSyn-mcherry virus showing HIP-NILB-iPSCs-MNs (mCherry$^{+}$ChAT$^{+}$GFP$^{+}$) in the lumbar spinal cord of HIP-NILB-iPSCs treated rabbits (Rabbit 430# and 480#). Scale bars, 200 μm or 50 μm. **(C)** Representative images showing STEM121$^{+}$ HIP-NILB-iPSCs-MNs labeled by mcherry in the lumbar spinal cords of grafted rabbits (Rabbit 429#, 430# and 480#) (2 months post-transplantation) after gastrocnemius intramuscular injection rAAV2-retro-hSyn-mcherry viruses for 1 month. Scale bars, 200 μm or 50 μm.

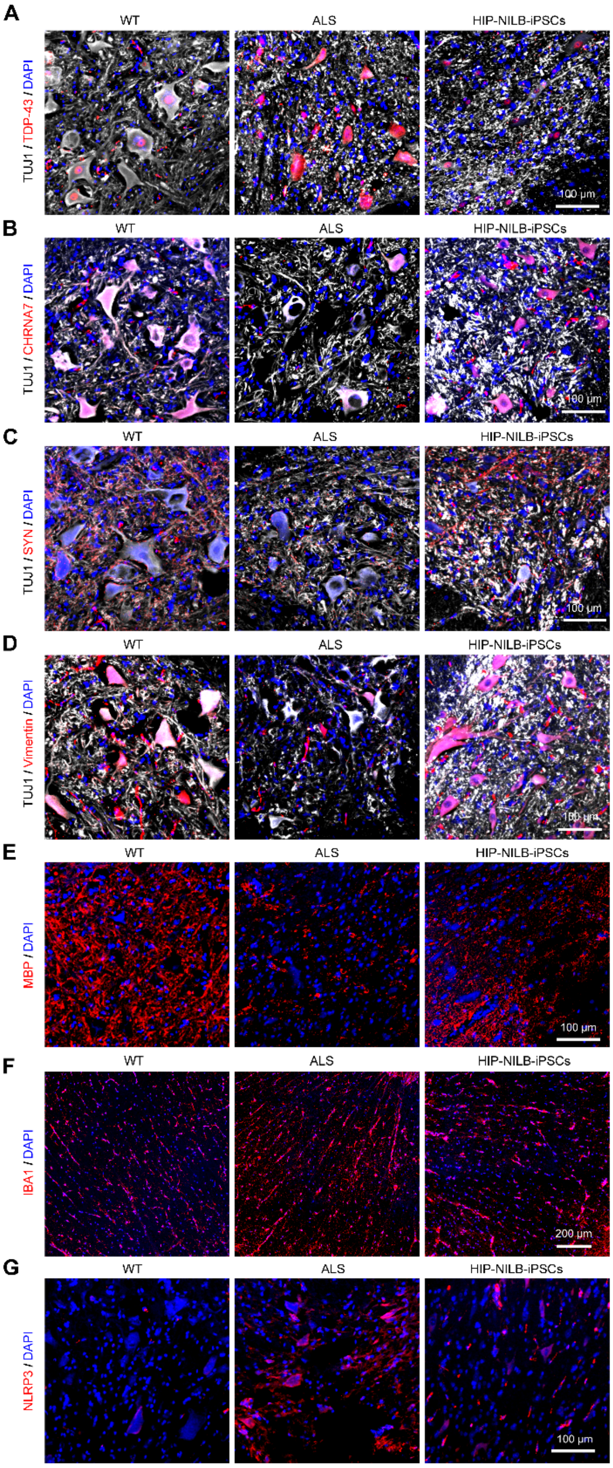


**Fig. S13. HIP-NILB-iPSCs improved ALS pathology and neural microenvironments in ALS rabbits.**

**(A)** Representative images showing lower magnification images of TDP-43 corresponding to Fig. 6E. Scale bar, 100 µm. **(B-G)** Representative images showing lower magnification images of (B-E) CHRNA7, SYN, Vimentin and MBP (corresponding to Fig. 6M), and (F-G) IBA1 and NLRP3 (corresponding to Fig. 6N), respectively. Scale bars, 200 µm or 100 µm.

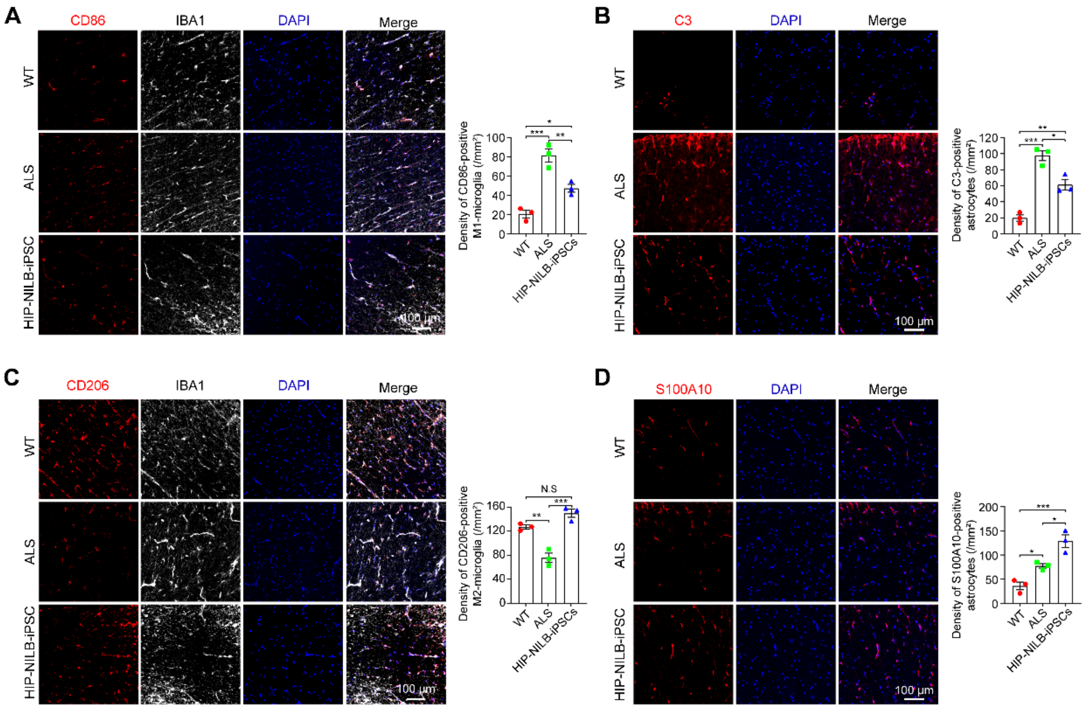


**Fig. S14. HIP-NILB-iPSCs attenuated neural inflammatory in ALS rabbits.**

**(A-B)** Representative images of (A) CD86 and (B) C3 in WT, ALS and HIP-NILB-iPSCs treated rabbits and the quantifications (n=3). Scale bar, 100 µm. **(C-D)** Representative images of (C) CD206 and (D) S100A10 in WT, ALS and HIP-NILB-iPSCs treated rabbits and the quantifications (n=3). Scale bar, 100 µm. The data were analyzed by one-way ANOVA followed by Tukey's multiple comparisons. Data was presented as mean ± SEM. *$p < 0.05$, **$p < 0.01$, ***$p < 0.001$, N.S, not significant.

**Table S1. sgRNA sequences and real-time PCR primers were used.**

| sgRNA and primers | Sequences (5'-3') |
|---|---|
| *B2M-sgRNA* | GAGTAGCGCGAGCACAGCTAAGG |
| *CIITA-sgRNA* | GCCTCCCAACATCTCCAGACCGG |
| *IL6ST-sgRNA* | GGTGGAATGGACTACTCCAAGGG |
| *TNFRSF1A-sgRNA* | GTGGACCGGGACACCGTGTGTGG |
| *hCD47*-F | TATCCTCGCTGTGGTTGGACTG |
| *hCD47*-R | TAGTCCAAGTAATTGTGCTAGAGC |
| *hCD24*-F | CACGCAGATTTATTCCAGTGAAAC |
| *hCD24*-R | GACCACGAAGAGACTGGCTGTT |
| *hIgG-Fc*-F | GAGAAGACGTTCCCCTGCTG |
| *hIgG-Fc*-R | AGGCTTCTATCCCAGCGACA |
| *hCTLA4*-F | ACGGGACTCTACATCTGCAAGG |
| *hCTLA4*-R | TCAGAATCTGGGCA |
| *hPDL1*-F | TGCCGACTACAAGCGAATTACTG |
| *hPDL1*-R | CTGCTTGTCCAGATGACTTCGG |
| *hGAPDH-F* | CGTGCCGCCTGGAGAAACCTG |
| *hGAPDH-R* | AGAGTGGGAGTTGCTGTTGAAGTCG |
| *rCHRNA7*-F | AGGAGCTGGTCAAGAACTACA |
| *rCHRNA7*-R | AAACTTGGTTCTTCTCATCCACG |
| *rSYN*-F | CCGGTTAGATAGCAGAGGGC |
| *rSYN*-R | TGAATTCCTTTACACCACATCTGC |
| *rVimentin*-F | GAAGGCGAAGAGAGCAGGATT |
| *rVimentin*-R | GTGTCAACCAGAGGGAGTGA |
| *rMBP*-F | TCCGGCAAGCAGGTTCC |
| *rMBP*-R | CCTTGTACAGGTTGCACAGC |
| *rNLRP3*-F | GGCCATCAACAGGAGAGACC |
| *rNLRP3*-R | TGGCATATCACAGTGGCGTT |
| *rGFAP*-F | GACATCGAGATCGCCACCTA |
| *rGFAP*-R | GTGACCTTGTGACTTTCCCCT |
| *IBA1*-F | GGAAGCCTTCAAGCGGAAATAC |
| *rIBA1*-R | GCTCTAGGTGAGTCTTGGGG |
| *rIFNG*-F | ATTGGCCTTTCTGAGCTGATTTTCC |
| *rIFNG*-R | GGCGTCAAGCCACAAGGC |
| *rIL17A*-F | TGCTGAAGGGAATGAGGACC |
| *rIL17A*-R | AAAGCCCTTAGGCCATGTGA |
| *rCD86*-F | CAGATCAAGGACAAGGGCGT |
| *rCD86*-R | AGCCTTGTACAGACGAGCAG |
| *rCD204*-F | GTGAAGTTTGATGCTCGCTCC |
| *rCD204*-R | TGAGGACGGCAAACACAAGG |
| *rTGFB2*-F | TTGTTACAACACCCTCTGGCT |
| *rTGFB2*-R | CTGTAGAAGGTGGGTGGGATG |
| *rNTF3*-F | ATGTCGACGTCCCTGGAATC |
| *rNTF3*-R | TTGTTCACCTGTAAGATCGTGG |
| *rNTF4*-F | GCTGAGATGTCAGGAAGGAAGG |
| *rNTF4*-R | ATCTCTCGGAGCACCTGTCA |
| *rNGF*-F | CCCCGGAGGTTCGTTGACC |
| *rNGF*-R | CACGATCACAGGCCAGAACTC |
| *rBDNF*-F | CTACCCAGTCGTATGTGCGG |
| *rBDNF*-R | ATCCTTATGAACCGCCAGCC |
| *rGAPDH-F* | GGCAAAGTGGATGTTGTCGC |
| *rGAPDH-R* | GCCGTGGGTGGAATCATACT |
